# Comparative analysis based on transcriptomics and metabolomics data reveal differences between emmer and durum wheat in response to nitrogen starvation

**DOI:** 10.1101/2020.02.03.931717

**Authors:** Romina Beleggia, Nooshin Omranian, Yan Holtz, Tania Gioia, Fabio Fiorani, Franca M. Nigro, Nicola Pecchioni, Pasquale De Vita, Urlich Schurr, Jaques David, Zoran Nikoloski, Roberto Papa

## Abstract

Mounting evidence indicates the key role of Nitrogen (N) on diverse processes in plant, including not only yield but also development and defense. Using a combined transcriptomics and metabolomics approach, we studied the response of seedlings to N starvation of two different tetraploid wheat genotypes from the two main domesticated subspecies, emmer (*Triticum turgidum* ssp. *dicoccum*) and durum wheat (*Triticum turgidum* ssp. *durum*). We found that durum wheat exhibits broader and stronger response in comparison to emmer as evidenced by the analysis of the differential expression pattern of both genes and metabolites and gene enrichment analysis. Emmer and durum wheat showed major differences in the responses to N starvation for transcription factor families. While emmer showed differential reduction in the levels of primary metabolites to N starvation, durum wheat exhibited increased levels of most metabolites, including GABA as an indicator of metabolic imbalance. The correlation-based networks including the differentially expressed genes and metabolites revealed tighter regulation of metabolism in durum wheat in comparison to emmer, as evidenced by the larger number of significant correlations. We also found that glutamate and GABA had highest values of centrality in the metabolic correlation network, suggesting their critical role in the genotype-specific response to N starvation of emmer and durum wheat, respectively. Moreover, this finding indicates that there might be contrasting strategies associated to GABA and Glutamate signaling modulating shoot *vs* root growth in the two different wheat subspecies.

## Introduction

Availability and uptake of nitrogen (N) is considered a major driver of growth (Lea and Azevedo, 2006). Indeed, N is an essential nutrient for all organisms, including plants, and is required for the biosynthesis of macromolecules, such as proteins, nucleic acids, and chlorophyll, and for the synthesis of many secondary metabolites with different roles in adaptation and signaling (Miller *et al.*, 2007). As a result, N deficiency (limited availability) and starvation (complete absence) dramatically affects plant growth and metabolism (Obata and Fernie, 2012).

However, only 30–50% of supplied N is taken up by crops (Raun and Johnson, 1999), and the remainder is lost by denitrification or leaching into terrestrial ecosystems, causing eutrophication and contamination of drinking water (Cassman *et al.*, 2003). Therefore, plant breeding efforts should be combined with improvement of crop management towards a more efficient use of N also to limit the use of fossil energy and environmental pollution (Ayadi *et al*., 2014; Ruisi *et al*., 2015). Towards this key objective, it is necessary to understand how plants react and cope with low N availability and identify the molecular basis of the natural genetic variation for adaptation to low N conditions.

Understanding the molecular mechanisms underlying the variation in traits responsible for the phenotypic plasticity in crop and wild species is a key step in addressing the challenges of modern agriculture, such as resilience to climate changes (Godfray *et al.*, 2010). In particular, understanding the genetic variation in N metabolism in major crop species, such as wheat, is expected to provide novel strategies for crop improvement (Kant *et al*., 2011; Xu *et al*., 2012; Hawkesford, 2017).

In an increasing number of model and crop species, transcriptome studies have highlighted the complexity of the regulatory mechanisms involved in the control of leaf or root gene expression under both N-limiting and non-limiting conditions (Krapp *et al*., 2011; Humbert *et al.*, 2013; Simons *et al.*, 2014; Curci *et al*., 2017). In addition, studies about the response of several cereal (e.g. rice, barley, sorghum, and wheat) to N starvation have highlighted differentially expressed genes (DEGs) involved in the response (Curci *et al*., 2017; Zuluaga *et al*., 2017; Yang *et al*., 2015, Gelli *et al*., 2014; Guo *et al.*, 2014). For instance, Gelli *et al.* (2014) compared transcriptomic levels in four tolerant and three sensitive sorghum genotypes to low N condition. Furthermore, Chen *et al.* (2011) and Hao *et al*. (2011) compared gene expression changes in response to N stress in two maize and soybean genotypes with contrasting low N tolerance.

Several works reported the combination of different ‘omics’ approaches in the evaluation of different crops responses to N starvation (Scheible *et al*., 2004; Amiour *et al*., 2012; Bielecka *et al*., 2015; Vicente *et al.*, 2016; Yu *et al*., 2017). Nevertheless, a limitation in these studies was the focus on a single genotype.

The analysis of gene expression can be complemented and expanded by using data on metabolite levels and their joint investigation with the help of network analysis approaches. The latter approaches have been useful in highlighting the role of metabolites in particular processes, but also for understanding the structure and regulation of the underlying metabolic and gene regulatory processes (Hirai *et al*., 2005; Caldana *et al*., 2011; Toubiana *et a*l., 2012, 2016; Beleggia *et al*., 2016).

The aim of this study was to investigate and compare the transcriptomic and metabolomics responses of two genotypes of tetraploid wheats (one emmer landrace and one elite durum wheat cultivar) to N starvation at the vegetative stage (seedling growth) that showed phenotypic responses to differences in N availability. Tetraploid wheats, (*Triticum turgidum* L. 2n=4x=28; AABB genome), alongside with einkorn and barley, were domesticated in the Fertile Crescent, and durum wheat derived from domesticated emmer (*Triticum turgidum* ssp. *dicoccum*) through a rather long human-driven selection process, including distinct and sequential domestication bottlenecks and continuous gene flow from wild emmer (*Triticum turgidum* ssp. *dicoccoides*) (Nesbitt and Samuel, 1998; Tanno and Willcox, 2006; Luo *et al*., 2007; Nevo, 2014). Here we obtained transcriptomics and metabolomics data from one emmer and one durum wheat genotype—the parents of a RIL population developed at CREA-CI Foggia (Russo *et al*., 2014). Our integrative analyses facilitated an in-depth molecular characterization and the comparison of tetraploid wheats responses to N starvation.

## Results

### Morphological and physiological differences under the two N conditions

First, we investigated the effect of N-starvation on plant growth by the evaluation of 13 complex traits, namely 12 morphological traits, including: Total leaf number (TLN); Total leaf area (TLA); Shoot fresh weight (SFW); Primary visible root length (PRL); Lateral visible root length (LRL); Total visible root length (TRL); Visible root system depth (RSD); Visible root system width (RSW); Root dry weight (RDW); specific root length (SRL); Total visible root length/total leaf area ratio (TRL/TLA); Lateral visible root length/ Primary visible root length ratio (LRL/PRL), as well as one physiological trait, Leaf chlorophyll content (SPAD) in emmer (Molise Sel. Colli) and durum wheat (Simeto). Table 1 shows the significant changes according to a two-way ANOVA due to genotype (G), N treatment (N) and their interaction (GxN). The traits TLA and SFW showed significant differences due to G (higher values in durum wheat) and N effect (higher values under optimal N condition). There were three measured traits, namely TLN, RDW and TRL/TLA which were significantly affected by N starvation. Finally, for SRL and SPAD, a significant effect due to the GxN interaction was observed. For instance, emmer at optimal N exhibited the largest value of SRL in comparison to emmer at N starvation and durum wheat in both N conditions. The opposite held for SPAD, for which durum wheat showed the highest value at optimal N compared to durum wheat at - N and for both treatments of emmer.

**Table 1.** Summary statistics and differential behavior for 12 morphological and one physiological trait in emmer and durum wheat under two N conditions: N starvation (-N) and optimal N (+N) condition. Data are reported as mean ± SE

|  | G |  |  | N |  |  | G x N |  |  |  |  |
| --- | --- | --- | --- | --- | --- | --- | --- | --- | --- | --- | --- |
|  | emmer | durum wheat | P value | -N | +N | P value | emmer x (-N) | emmer x (+N) | durum wheat x (-N) | durum wheat x (+N) | P value |
| **TLN** | 3.88±0.54 | 4.81±0.48 | n.s. | **3.50±0.19<sup>b</sup>** | **5.19±0.59<sup>a</sup>** | **0.0173** | 3.00±0.00 | 4.75±0.92 | 4.00±0.00 | 5.63±0.80 | n.s. |
| **TLA (cm<sup>2</sup>)** | **24.85±5.66<sup>b</sup>** | **41.92±8.39<sup>a</sup>** | **0.0304** | **19.98±1.43<sup>b</sup>** | **46.79±8.35<sup>a</sup>** | **0.0023** | 17.22±1.21 | 32.48±10.45 | 22.74±1.75 | 61.10±8.94 | n.s. |
| **SFW (g)** | **0.63±0.19<sup>b</sup>** | **1.17±0.30<sup>a</sup>** | **0.0453** | **0.42 ±0.05<sup>b</sup>** | **1.39 ±0.28<sup>a</sup>** | **0.0017** | 0.34±0.04 | 0.93±0.32 | 0.50±0.07 | 1.85±0.35 | n.s. |
| **PRL (cm)** | 155.58±26.79 | 172.72±25.42 | n.s. | 191.72±21.60 | 136.59±26.47 | n.s. | 189.27±31.72 | 121.89±39.82 | 194.16±34.17 | 151.28±39.25 | n.s. |
| **LRL (cm)** | 20.49±7.00 | 10.89±4.04 | n.s. | 15.28±5.19 | 16.10±6.70 | n.s. | 18.60±8.34 | 22.38±12.53 | 11.96±6.99 | 9.83±5.13 | n.s. |
| **TRL (cm)** | 176.07±32.17 | 183.62±27.96 | n.s. | 206.99±24.99 | 152.69±31.39 | n.s. | 207.87±39.64 | 144.27±50.83 | 206.12±36.65 | 161.11±44.36 | n.s. |
| **RSD (cm)** | 62.84±5.38 | 65.34± 3.66 | n.s. | 70.28±2.54 | 57.90±5.04 | n.s. | 68.69±4.09 | 56.98±9.76 | 71.86±3.41 | 58.82±4.74 | n.s. |
| **RSW (cm)** | 23.31±3.26 | 22.82±3.22 | n.s. | 24.79±3.08 | 21.34±3.27 | n.s. | 28.72±3.47 | 17.89±4.25 | 20.87±4.68 | 24.78±4.89 | n.s. |
| **RDW (g)** | 0.02±0.01 | 0.03±0.00 | n.s. | **0.03±0.00<sup>a</sup>** | **0.02±0.00<sup>b</sup>** | **0.0228** | 0.03±0.01 | 0.01±0.00 | 0.03±0.01 | 0.02±0.00 | n.s. |
| **SRL (m g<sup>-1</sup>)** | **107.31±15.77<sup>a</sup>** | **67.69±5.99<sup>b</sup>** | **0.0094** | **70.31±5.93<sup>b</sup>** | **104.69±16.65<sup>a</sup>** | **0.0201** | **75.75±7.06<sup>b</sup>** | **138.87±21.15<sup>a</sup>** | **64.87±9.72<sup>b</sup>** | **70.52±8.21<sup>b</sup>** | **0.0449** |
| **TRL/TLA (cm cm<sup>-2</sup>)** | 7.92±1.65 | 5.98±1.62 | n.s. | **10.57±1.28<sup>a</sup>** | **3.34±0.45<sup>b</sup>** | **0.0002** | 11.68±1.71 | 4.17±0.54 | 9.46±1.98 | 2.50±0.43 | n.s. |
| **LRL/PRL** | 0.11±0.04 | 0.06±0.02 | n.s. | 0.07±0.02 | 0.10±0.04 | n.s. | 0.09±0.04 | 0.14±0.07 | 0.06±0.03 | 0.05±0.02 | n.s. |
| **SPAD** | **26.55±1.22<sup>b</sup>** | **34.91±2.24<sup>a</sup>** | **<0.0001** | **26.91±1.24<sup>b</sup>** | **34.55±2.41<sup>a</sup>** | **0.0001** | **24.35±0.91<sup>b</sup>** | **28.75±1.72<sup>b</sup>** | **29.48±1.40<sup>b</sup>** | **40.35±1.31<sup>a</sup>** | **0.0353** |

**TLN:** Total leaf number; **TLA:** Total leaf area (cm<sup>2</sup>) Calculated on all the leaf as leaf length. maximum width.0.858 (Kalra and Dhiman. 1976); **SFW:** Shoot fresh weight (g); **PRL:** Primary visible root length (cm); **LRL:** Lateral visible root length (cm); **TRL:** Total visible root length (cm); **RSD:** Visible root system depth (cm); **RSW:** Visible root system width (cm); **RDW:** Root dry weight (g); **TRL/TLA:** Total visible root length/total leaf area (cm cm<sup>-2</sup>); **SRL:** specific root length (defined as TRL/RDW); **SPAD:** Leaf chlorophyll content (SPAD units). Values annotated in bold are significantly different (Tukey's test) and the character 'a'/'b' implies the higher/lower observed value for each significant change between the genotypes, the two N conditions and their interaction.

### Transcriptomic differences between the two N conditions

A global transcriptome analysis for the comparison of the two analyzed tetraploid wheat genotypes was performed using RNA-Seq Illumina technology resulting in 9.9 to 19.5 million reads per genotype (Table S1). These numbers were reduced after additional processing steps (see Methods) by 4.3 to 7.5%, depending on the sample. The cleaned reads were mapped on the bread wheat reference covering, on average, 70% of all reads in the analyzed genotypes (Table S1).

We used the mapped reads to assess the DEGs in each genotype between the two N conditions, i.e. N starvation and optimal N condition. The total number of genes expressed in emmer and durum wheat were 27,792 and 28,812, respectively. The number of significant DEGs for emmer was 1,788, while in durum wheat it was 3,129. The number of DEGs specific to durum wheat was ∼3.2-fold larger than in emmer, and the number of DEGs common to the two genotypes was 1,095 (Figure 1A). In addition, the number of the up-regulated DEGs specific to durum wheat was 2.5-fold larger than those specific to emmer, while the number of down-regulated DEGs specific to durum wheat was 3.5-fold larger than those specific to emmer (Figure 1B). Therefore, we found a stronger transcriptional response in durum wheat to the change in N availability in comparison to emmer.

**Figure 1.**
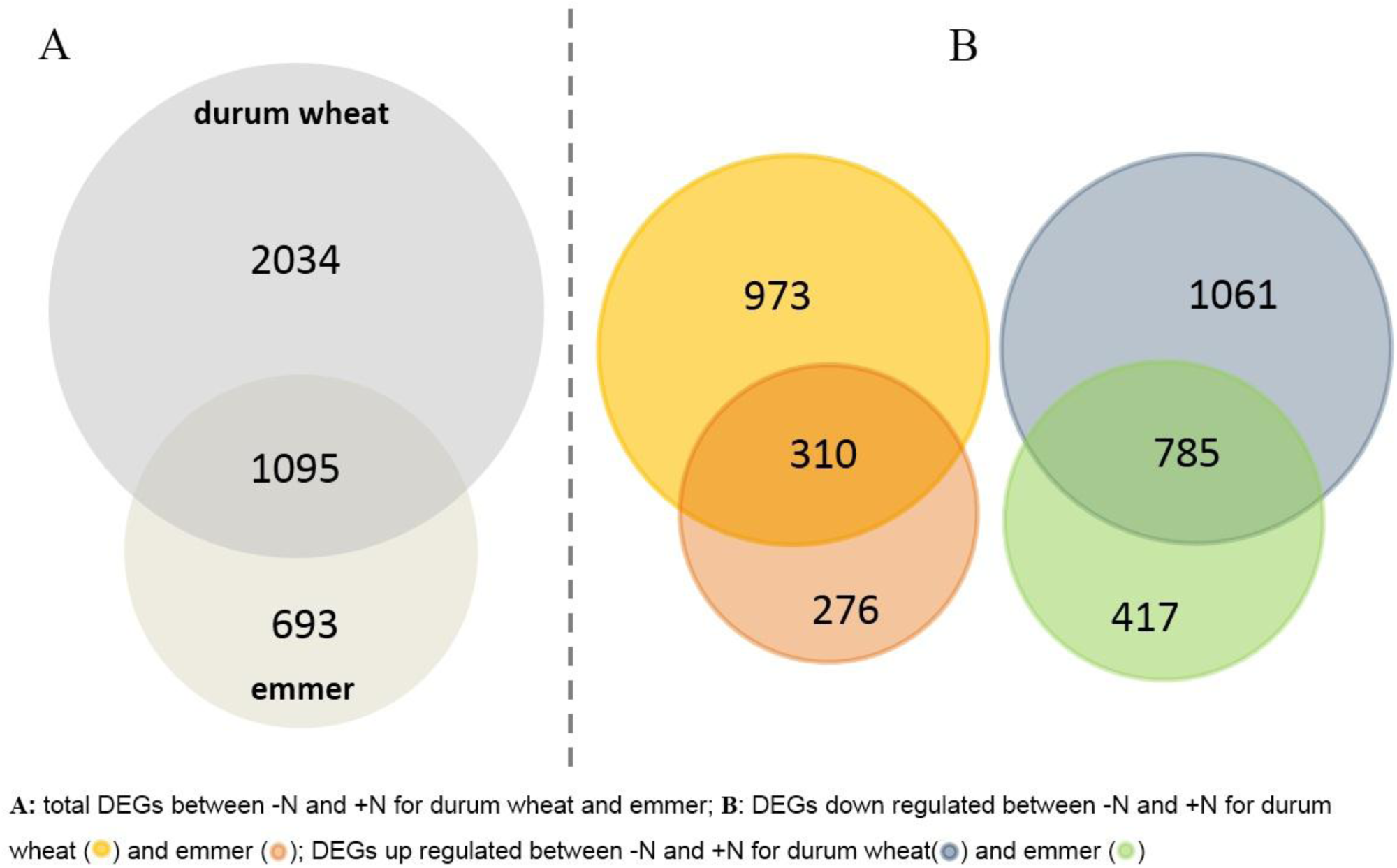
Differentially expressed genes between N starvation (-N) and optimal N (+N) conditions in durum wheat and emmer.

### Functions of DEGs in emmer and durum wheat between the two N conditions

The functional annotation of DEGs either common or specific to one of the genotypes, were reported in Table S2. Several DEGs were directly involved in N metabolism and transport. The key DEGs involved in nitrate assimilation, i.e. the gene coding for asparagine synthetase and aspartate aminotransferase, were up-regulated in both genotypes. In durum wheat, the gene coding for nitrate reductase was up-regulated, while the genes orthologous to *Arabidopsis* glutamine synthetase and glutamate dehydrogenase family were down-regulated in response to N-starvation (Table S2). A similar result was reported by Curci *et al*. (2017) for the response of durum wheat leaves to N chronic starvation during grain filling. One nitrate transporter and two ammonium transporters were found among the DEGs in emmer (Table S2). Interestingly, other DEGs associated with the translocation of other nutrient (potassium (8 genes), phosphate (1- [PhO1]), sulfate (1), zinc (1), calcium (8), copper (2), magnesium (3) and ABC transporter (6)) also changed under N starvation (Table S2).

A general alteration was observed for genes participating in carbon metabolism, especially for those involved in glycolysis, tricarboxylic acid cycle (TCA), photosynthesis and photorespiration, particularly in durum wheat (Table S2). Notably, gene coding for Phosphoglycerate kinase (PGK), Pyruvate kinase (PK), Glyceraldehyde 3-phosphate dehydrogenase (GAPDH), and Fructose bisphosphate aldolase were up-regulated and specific to durum wheat, while Pyruvate dehydrogenase E1-component subunit alpha (PDHA), Pyrophosphate--fructose 6-phosphate 1-phosphotransferase subunit alpha (PFP-ALPHA), and ATP-dependent 6-phosphofructokinase (PFK1) were up-regulated and specific to emmer. Concerning the Pentose phosphate pathway, one DEG encoding for glucose-6-phosphate dehydrogenase (G6PD) was up-regulated in both genotypes, while two orthologs to ribose-5-phosphate-isomerase (Rpi) were up-regulated only in durum wheat. Notably, orthologues to RuBisCO (5 DEGs) and ferrodoxin (3 DEGs) were up regulated only in durum wheat.

Transcription factors from the ARFs (5 DEGs) and NF-Y (3 DEGs) families were found to be down-regulated in both genotypes, while the MYB family (1 DEGs) was up-regulated in durum wheat and PTACs (5 DEGs) families were up-regulated in both genotypes (Table S2). In addition, 35 protein kinases (PKs) were identified as DEGs, of which 13 were common to the two genotypes, while six and 16 were found as DEGs specific to emmer and durum wheat, respectively. Generally, N starvation causes several stress responses. About two thirds of DEGs common or specific to each genotype were up-regulated, and among them there were several antioxidant enzymes encoding genes, such as: superoxide dismutase (SOD), catalase (CAT), glutathione peroxidase (GPX), peroxiredoxin (Prx), and lipoxygenases (LOXs), as well as enzymes of the ascorbate-glutathione cycle, such as: glutathione reductase (GR) or those involved in the biosynthesis of secondary metabolites (Table S2).

### GO enrichment analysis of DEGs

To investigate the transcriptomic changes in leaves of emmer and durum wheat under the two N conditions, we assessed the GO enrichment in the set of DEGs (see Methods).

The GO terms identified were categorized into 21 and 23 categories for emmer and durum wheat, respectively (Figure 2, Table S3). In both genotypes, the highest number of DEGs up-regulated were included in the categories ‘cellular process’, ‘metabolic process’, ‘binding’ and ‘catalytic’ while those that were differentially down-regulated were principally grouped into ‘binding’ and ‘catalytic’ categories. Differences between the two genotypes were observed with respect to the molecular function category ‘transcription regulator’ which was enriched in both the down- and up-regulated DEGs in emmer and durum wheat, respectively, and in the GO terms associated with biological process categories ‘regulation of biological process’ and ‘reproductive process’, which were only enriched in durum wheat for up-regulated DEGs.

**Figure 2.**
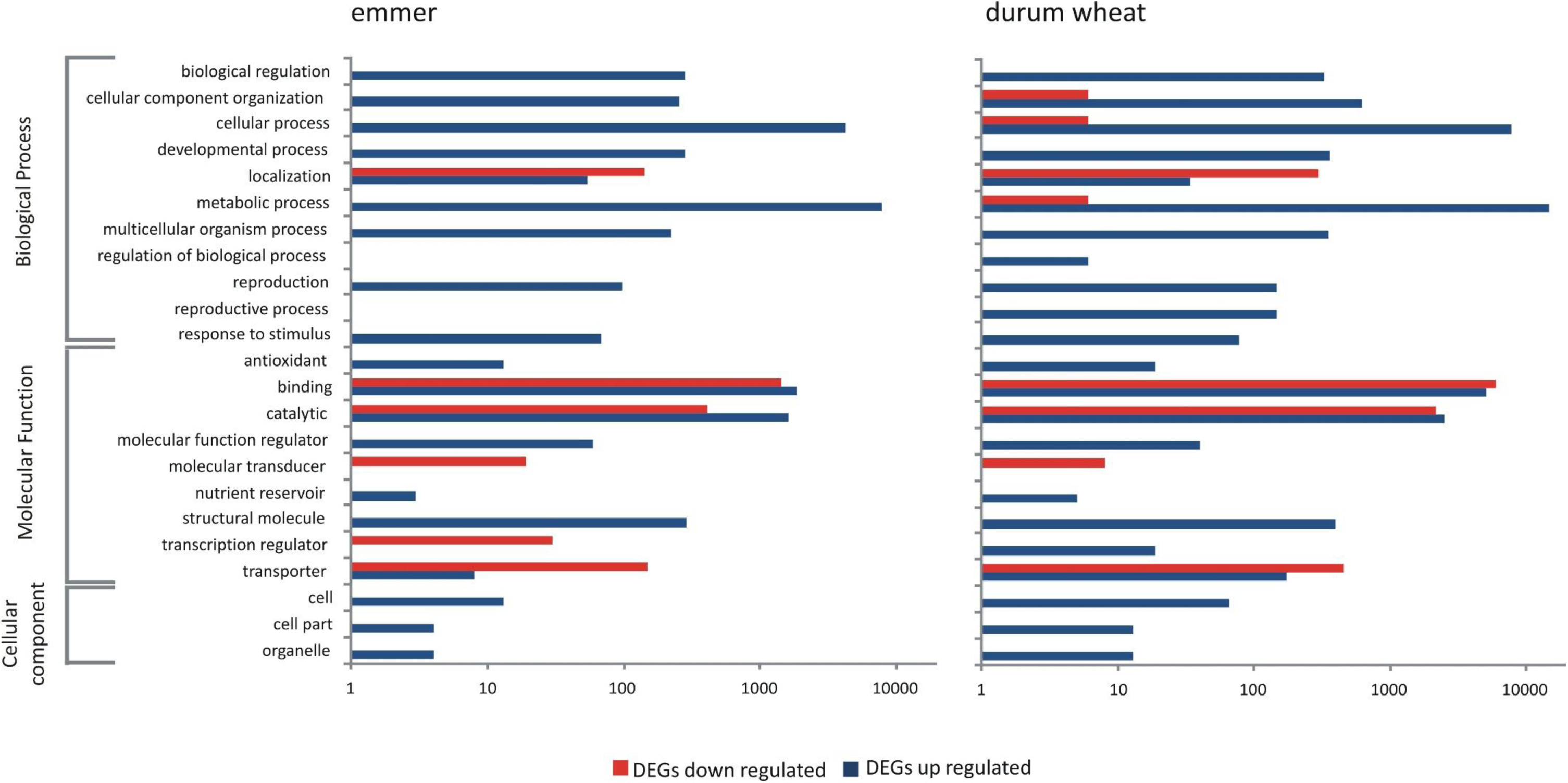
Comparison of Gene Ontology classifications of DEGs in emmer and durum wheat. Blue and red color indicates the number of up- and down-regulated DEGs, respectively. All DEGs are categorized into 21and 23 functional groups based on GO classification for emmer and durum wheat, respectively.

Extended list of over represented GO terms with the p-value of at most 10^−5^ for emmer and durum wheat is reported in Table S4. Notably, all GO terms of the categories ‘cellular process’, ‘metabolic process’, ‘binding’ and ‘catalytic’ (e.g. those involving the nitrogen) which were enriched in emmer were also found in durum wheat. Durum wheat showed also specific over-represented GO terms in several categories; for instance, these included the cellular amino acid, oxoacid or organic acid metabolic processes, or the metabolic/biosynthetic process of isopentenyl diphosphate (Table S4). In addition, regarding the categories ‘binding’ and ‘catalytic activity’, durum wheat showed different over-represented GO terms among the DEGs differentially up- and down-regulated. For example, GO terms of oxidoreductase, ligase, hydrolase (on glycosyl bond or O-glycosyl compounds), lyase and transferase activity were not enriched in the down-regulated DEGs, while GO terms of kinase, protein kinase, protein serine/threonine kinase and phosphotransferase activity were not enriched on the up-regulated DEGs (Table S4).

### Metabolic differences between the two N conditions

A total of 46 metabolites were identified and quantified using GC-MS (see Methods). These included 41 polar and five non-polar compounds, divided into the following compound classes: amino acids, organic acids, sugars and sugar alcohols, fatty acids, polycosanol and phytosterols. The data were analyzed using two-way ANOVA and significant differences (P ≤ 0.05) for 23 metabolites including the TCA cycle intermediates, some sugars, shikimic and quinic acids, several amino acids and GABA were reported (Table S5). A higher content of metabolites was found in emmer under optimal N condition in comparison to durum wheat (see Figure 3 for illustrative comparison). For all metabolites, a strong significant effect due to the interaction of the genotype and N treatment (GxN) was observed. Five metabolites (glutamic acid, aspartic acid, citric acid, saccharic acid and maltitol) showed also significant differences due to the effect of genotype (G) and treatment (N). Tryptophan showed only the genotype effect while aconitic and shikimic acids showed significant differences due to G apart from the GxN interaction.

**Figure 3.**
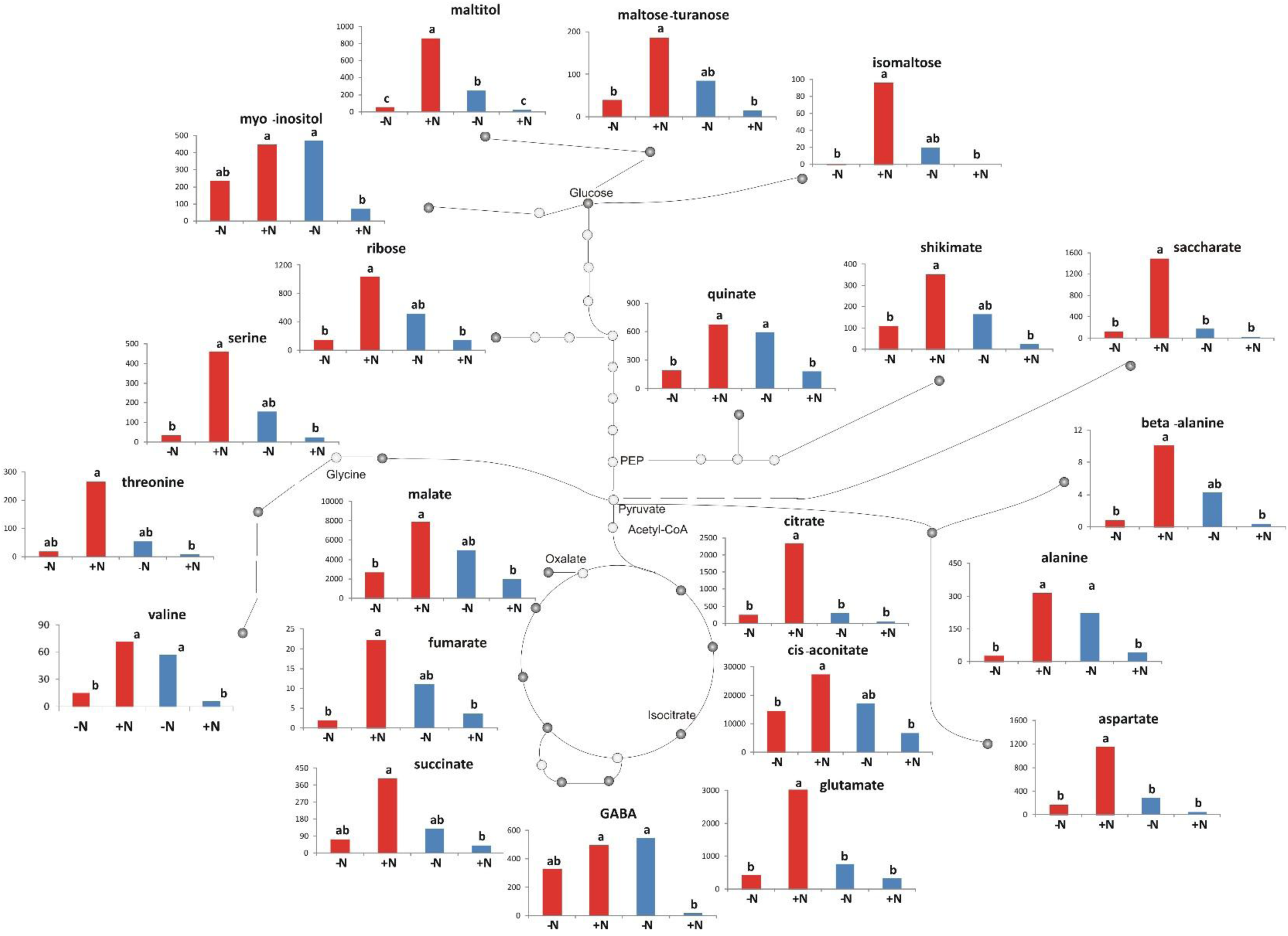
Metabolites exhibiting significant variation for emmer (red bars) and durum wheat (blue bars) under starvation (-N) and optimal (+N) levels due to the effect of GxN interaction. Bars with different letters are significantly different (p < 0.05)

### Network analysis of combined data sets

In general, the correlation structure among the combined data sets (transcripts and metabolites) of each genotype can be represented by a network, where a node denotes a transcript, or a metabolite and an edge stand for the presence of significant Pearson correlation between the data associated to the nodes. Overall, durum wheat showed 2.8-fold more significant correlations in comparison to emmer (Table 2). The intersection of the networks from the two genotypes included ∼397,000 edges, of which 99.3% did not demonstrate significant differences between the two networks (using Fisher’s z-transformation, see Methods). The latter set of edges (with no significant differences between the networks obtained for each genotype; by applying Fisher’s *z* transformation) is said to comprise the common network between the two genotypes that represents the 31.5% and 11.2% of the total correlations in the networks of emmer and durum wheat, respectively.

**Table 2.** Networks of emmer, durum wheat, intersection and common networks of transcripts and metabolites data.

|  | **emmer** | **durum wheat** | **(durum wheat /emmer)** | **intersection** | **Common**<br>(accepting the Fisher z test NULL hypothesis) |
| --- | --- | --- | --- | --- | --- |
| Number of edges in total | 1,249,637 | 3,500,971 | 2.8 | 396,571 | 393,779 |
| Number of edges DEG-DEG | 1,237,748 | 3,473,768 | 2.8 | 394,015 | 393,719 |
| Number of edges metabolite-metabolite | 185 | 157 | 0.85 | 65 | 60 |
| Number of edges DEG-metabolites | 11,704 | 27,046 | 2.3 | 2,491 | 0 |
| Number of nodes | 1,829 | 3,167 | 1.7 | 1,129 | 1,127 |
| Number of central nodes | 260 | 479 | 1.8 | 367 | 398 |
| Number of edges to the central nodes: DEGs – significantly behaved metabolites* | 1,898 | 4,590 | 2.4 | 1,217 | 0 |

\*significantly behaved metabolites considering the effect of G, N and GxN of the ANOVA model.

Because we are interested in understanding if the differences in correlation could reflect the differences in regulation of transcripts and metabolites, we considered only the significant correlations between DEGs and significantly altered metabolites under the two N conditions; the number of such correlations in durum wheat was 2.3-fold larger than in emmer. Focusing the attention only on those metabolites that showed differential behavior between the two N conditions, as reported above, we observed that for emmer GABA is involved in the smallest number of edges (12), while maltitol participates in the largest number of edges (1,667). In durum wheat we find almost contrasting situation, isomaltose was involved in the smallest number of edges (28), while GABA exhibited the largest number of edges (2,954) (Table S6).

The effect of the observed differences between the correlation structures (i.e. networks) obtained for both genotypes can be investigated for each node and can be summarized by its centrality in the network. In this context, we selected those nodes showing the centrality measures (i.e., degree and betweenness) greater than the corresponding mean values in each genotypic-specific network. Considering the nodes of the two genotype-specific networks, those with a central role included 260 and 479 genes in emmer and durum wheat, respectively (Table 2). In durum wheat also the metabolites: myo-inositol, quinic acid and valine showed high values for both centrality measures. To refine the network, we next included only DEGs with high values of centrality and only metabolites that were significantly contrasted between the two N conditions. In general, the total number of edges decreased of about 79% and 85% for emmer and durum wheat, respectively (Table S6). The total number of edges between central DEGs and differentially behaved metabolites in durum wheat is higher than those in emmer by 3.6-fold for alanine and 479-fold for GABA. In contrast, in emmer, the number of edges between central DEGs and significantly contrasted metabolites: glutamic acid, isocitric acid, isomaltose, saccharic acid, serine, succinic acid, and threonine, were higher than those in durum wheat. Noteworthy, with aspartic acid, citric acid, fumaric acid and maltitol the number of edges was the same in both genotype-specific networks.

### Function of DEGs having a central role in the networks

To evaluate the common or specific responses to N starvation in the two genotypes, we looked for the annotated functions of the DEGs shared between the two genotype-specific networks with a central role in at least one of the two networks (Table S7). Several DEGs related to photosynthesis were expressed in both genotypes but in some cases, they showed a central role only in emmer-specific network (e.g. Chlorophyll synthase (CHLG)) while, in contrast Carboxyl-terminal-processing peptidase 3 (CTPA3), Cytochrome c biogenesis protein (CCS1), and magnesium-chelatase subunit ChlD (ChlD) were found to have a central role in the durum wheat-specific network. In the network specific to emmer the most central nodes coded for Pyruvate phosphate dikinase 1 (PPDK) and Pyruvate dehydrogenase E1component subunit alpha-3 (PDH-E1 ALPHA) which were down- and up-regulated, respectively.

In durum wheat-specific network, DEGs related to proteolysis as well as the synthesis of the cofactor FMN, that were up-regulated, had a central role in the network, and at the same time, Allantoinase (ALN), a key enzyme for biogenesis and degradation of allantoin and its degradation derivatives, essential in the assimilation, metabolism, transport, and storage of nitrogen in plants, was among the central nodes.

In both genotype-specific networks, different DEGs involved in the chloroplast development showed central roles (Table S7). Among the central DEGs, there were several genes related to detoxification and plant stress responses caused by N starvation. Only one DEG (Traes_2BL_CCD296233, down-regulated) encoding for the Stress Enhanced Protein 2 [SEP2], showed a central role in both genotype-specific networks (Table S7).

To highlight the differences between emmer and durum wheat, we also considered the putative annotation of the central DEGs in each genotype-specific network (see Table S8). In emmer, several genes involved in C metabolism or related to stress conditions responses were up-regulated; at the same time a DEG related to carbonic anhydrase (EC 4.2.1.1), involved in N metabolism, was down-regulated. In contrast, in durum wheat, several DEGs related to photosynthesis were differently regulated, i.e. chlorophyll synthase and the ferritin were up-regulated while the ferrochelatase was down-regulated. Importantly in durum wheat-specific network, there is also a central DEG (Traes_3AS_3CB8A9C01) for glutamate decarboxylase [GAD] which was up-regulated.

Figure 4 represented the genotype-specific networks of DEGs-metabolites reported in Table S7 and Table S8 for emmer (A) and durum wheat (B), respectively. As illustrated, the network structure was different between the two genotypes; consistently emmer-specific network showed a higher number of negative correlations between DEGs and metabolites while durum wheat-specific network has higher number of positively correlated DEGs and metabolites pairs. Of note, glutamic acid and valine were the metabolites highly connected to the other nodes in emmer-specific network while GABA, quinic acid, *myo*-inositol and valine were highly connected to the rest of the nodes in durum wheat-specific network.

**Figure 4.**
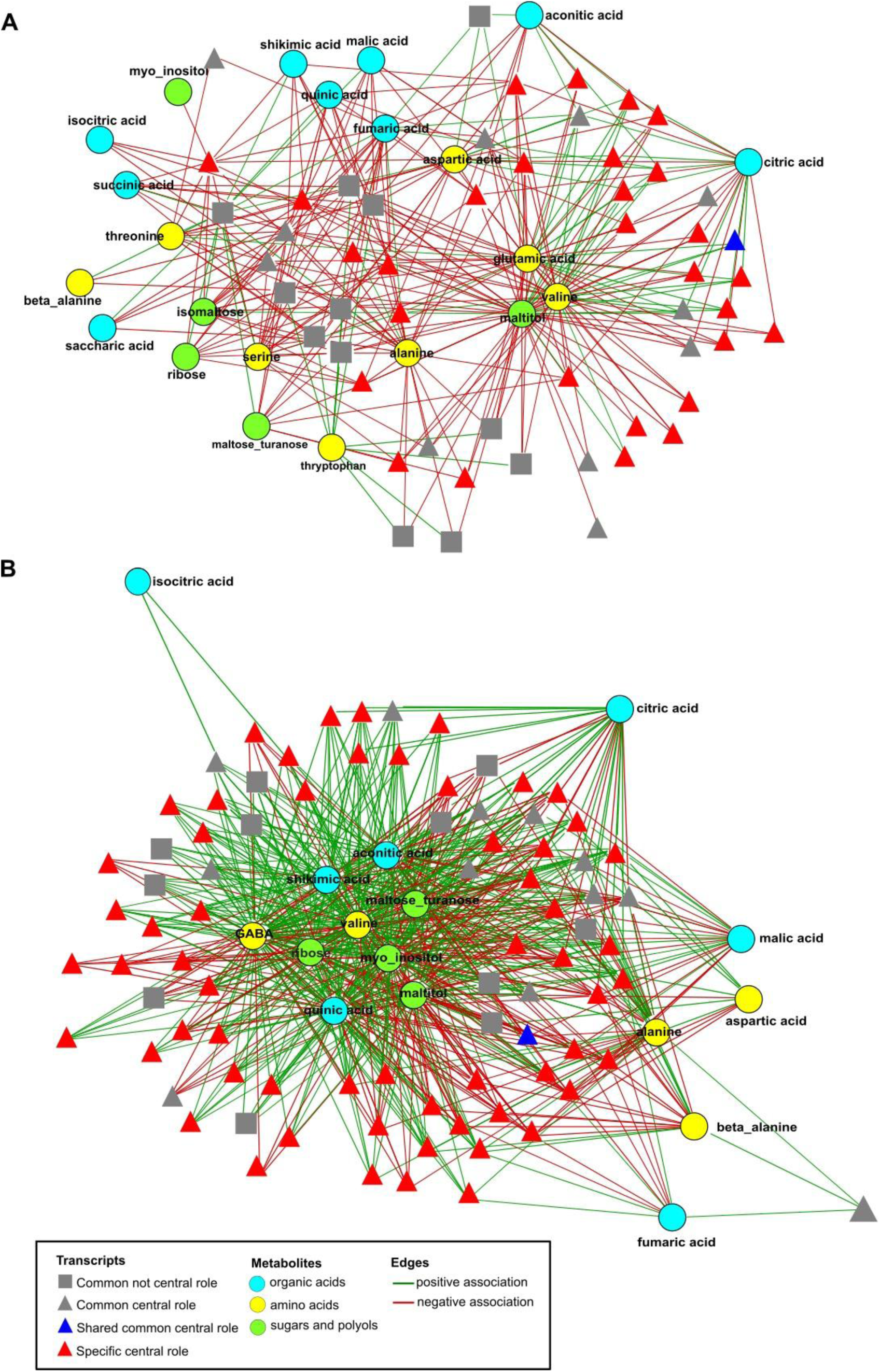
Genotype-specific DEGs-metabolites correlation networks, emmer (A) and durum wheat (B).

### DEGs position on the genome

We have also considered the position of DEGs in both genotypes on the physical map. In general, for each chromosome durum wheat showed a higher number of DEGs compared to emmer. In both genotypes, the larger number of DEGs was located on chromosome 2A, 2B, 4A, 5A and 5B, while lower number of genes was found in the chromosome 3B. Few genes were in chromosome 6B in emmer (Figure S1).

Figure 5 illustrates the location of down- and up-regulated DEGs with central role in the corresponding genotype-specific networks. Observing the results, the higher number of central nodes in the emmer-specific network was located on chromosome 2B, 4B, and 5A, while for durum wheat-specific network the higher number of central DEGs was located on chromosome 2A, 2B, 4A and 5B.

**Figure 5.**
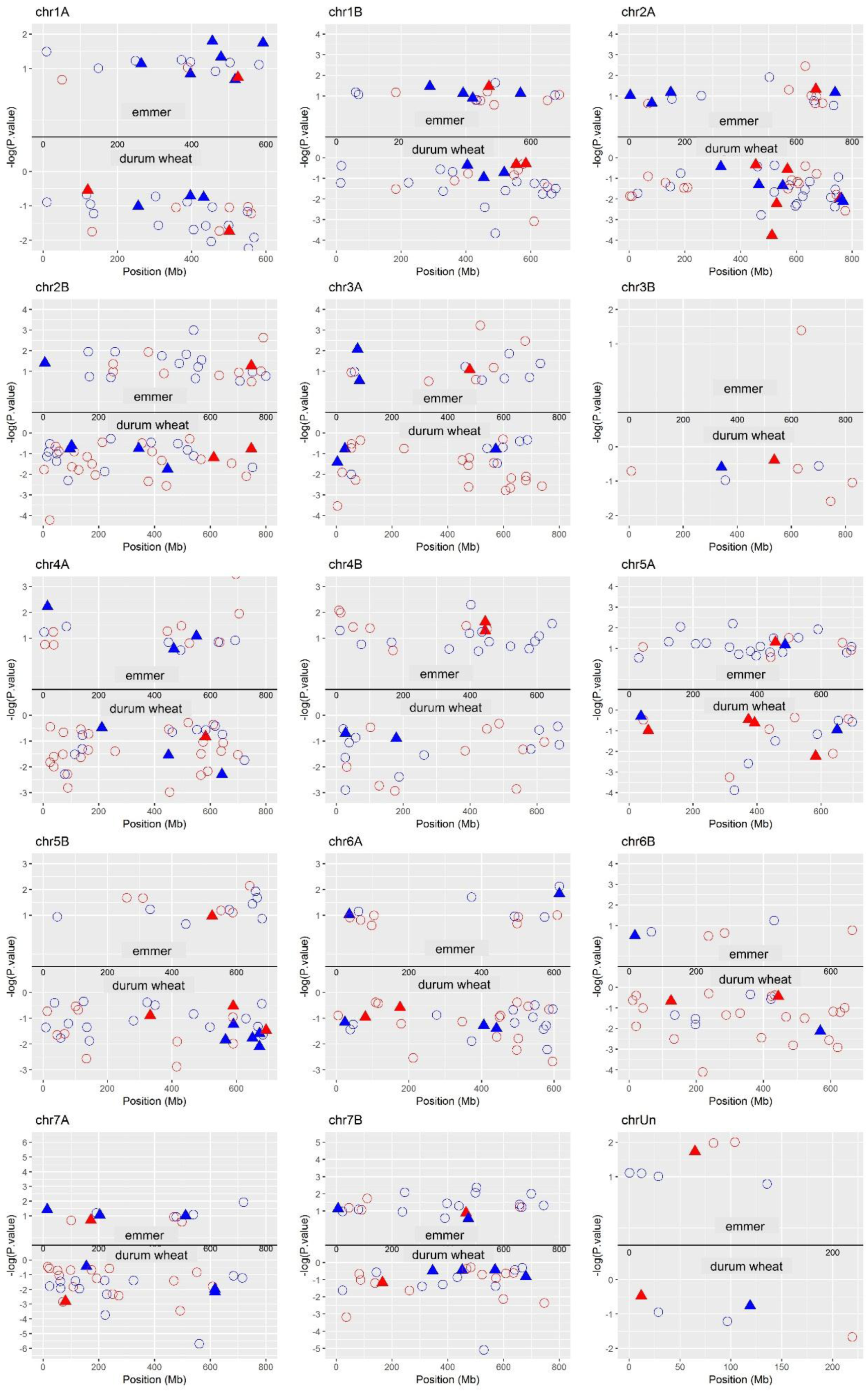
Position of the central DEGs in emmer- and durum wheat-specific networks on the physical map. DEGs having putative annotations were shown with filled triangles.

## Discussion

In a preceding work we found that emmer and durum wheat showed contrasting phenotypic responses associated to N starvation (Gioia *et al*., 2015). Here we present the results of gene expression and metabolites levels of emmer and durum wheat using two representative genotypes which were part of the previous investigation. Indeed, a striking result showed by our study is the major differences in the response to N starvation between our emmer and durum wheat genotypes based on their gene expression and metabolite levels. Emmer responded to the stress condition by slowing down all the metabolic functions, probably limiting his energy expenditure. On the contrary, durum wheat responded to the stress condition by activating a much larger number of genes (e.g. triggering more defense responsive pathways) and mechanisms resulting in an accumulation of metabolites in the investigated tissues (leaves) most likely associated to a metabolic imbalance. Moreover, evaluating the differences in plant growth, a significant growth variation under N starvation was observed in both genotypes according to the results reported by Gioia *et al*. (2015) which was more evident in the aerial part in durum wheat and in the below-ground part in emmer.

Durum wheat responded to N starvation with a much higher number of DEGs up-regulated. Some of these genes, directly involved in N metabolism, were differentially expressed exclusively in durum wheat (i.e. NR (up-regulated), GS and GDH (down-regulated)). In addition, the results of gene enrichment analysis indicate that emmer and durum wheat adapt to nitrogen starvation by a reprogramming of transcription. Transcription factors are important for controlling the expression of other genes in plant exposed to limited N condition or in complete starvation (Krapp *et al*., 2011; Yang *et al*., 2015; Curci *et al*., 2017) and, accordingly, our results showed as the regulation of transcripts was highly different and, in some case, with an opposite trend between emmer and durum wheat. In addition, some GO categories were only enriched for the DEGs in durum wheat, such as: the cellular amino acids, oxoacid or organic acids metabolism which were also highlighted by Huang *et al.* (2016) in their study on the transcriptomic evaluation in response to the imbalance of carbon: nitrogen ratio in rice seedling.

Moreover, the levels of metabolites showed significant differences in response to the N starvation in both emmer and durum wheat. In general, in stressed conditions a reduction in plant growth and photosynthesis is expected (Shaar-Moshe *et al*., 2018) and, consequently, this should lead to a decrease in monosaccharides content. Nevertheless, an increase in the starch and soluble sugars content was reported in the shoot of Arabidopsis thaliana under N starvation (Krapp *et al*., 2011). Accordingly, an increase of total sugars in both genotypes was observed, with a pronounced effect in durum wheat (which also showed a significant decrease of photosynthetic efficiency).

Consistently to the differences observed at transcriptomic level, the content of amino acids, under N starvation was lower in emmer (fold change = −2.7), while in durum wheat a higher accumulation (fold change= 1.7) of these metabolites was observed. Tschoep *et al*. (2009) showed that when Arabidopsis plants were grown under continuous N limitation, the total amino acids levels were found to be higher than under high N condition due to a metabolic imbalance. The results obtained in our conditions suggest a reduced use of amino acids for protein synthesis and growth in durum wheat that links with the reduction of photosynthetic activity under N starvation. On the other hand, the lower accumulation in emmer may indicate an earlier phase of the N starvation syndrome which could result in a drastically reduced, but still efficient, metabolism.

In this sense, it is also important to discuss carefully the behaviors of both glutamic acid and GABA, both altered in response to the N starvation condition in emmer and durum wheat. GABA is synthesized mainly from glutamate, closely associated with the TCA cycle, and having a signaling role (Bouchè and Fromm, 2004; Fait *et al*., 2008, Caldana *et al*., 2011). Two studies have suggested a signaling role of GABA during the nitrate uptake in both *Brassica napus* root (Beuve *et al*., 2004) and *Arabidopsis thaliana* (Barbosa *et al*., 2010). Moreover, Sulieman (2011) reported the important role of GABA in increasing of the efficiency of symbiotic N_2_ fixation in legumes. Michaeli and Fromm (2015), proposed that the metabolic and signaling functions of GABA has been evolved to be functionally entwined under nutrient starvation. Thus, it seems that GABA levels increase during plant nutrient starvation and energetically demanding stresses (Carillo, 2018), aspect that could be supported, from our data, by the negative correlation between the SPAD values (indicating reduced chlorophyll content) and the GABA content in durum wheat (r = −0.86; P = 0.0061). On the other hand, Forde and Lea (2007) reported the possible long-distance signaling role of glutamate between shoot and root as part of a network of N signaling pathways that enable the plant to monitor and adapt to changes in N status. In their model, when the shoot-derived glutamate arrive at the root tip, is sensed by plasma membrane glutamate receptors enabling meristematic activity in the root tip to respond to changes in the N/C status of the shoot. In our study the positive correlation in emmer between shoot glutamate and SRL (r = 0.97; P = 0.0001) could support this suggestion. In addition, the increase of the root morphological parameters in emmer under N starvation could be also sustained by a greater remobilization of the amino acids from the shoot to the root. The key role of GABA and glutamate is also supported by the results of the correlation-based network analysis integrating the information from both metabolites and transcripts. Indeed, durum wheat-specific network was characterized by the role of GABA that was associated to many (479) DEGs while in emmer-specific networks the glutamate was highly connected to many (201) other DEGs. This finding, on one hand, underlies their important role as signaling metabolites in stress conditions as those occurring during nitrogen starvation and, on the other hand, it may suggest the occurrence of two contrasting strategies based on GABA and glutamate signaling that appear associated to shoot and root growth, respectively.

The genotype-specific networks of the two tetraploid wheats showed different structures. Overall, only one DEG (down-regulated) common to both emmer and durum wheat showed a central role in the corresponding networks (i.e. Stress Enhanced Protein 2 -[SEP2]) which is a light-inducible gene as showed in *Arabidopsis thaliana* and rice (Umate, 2010). A previous work reported that the regulation of SEP gene expression by light stress is very specific while other physiological stresses, such as: cold, heat, wounding, desiccation, salt or oxidative stress, did not promote accumulation of SEP transcripts indicating that they were not triggered by photooxidative damage itself (Heddad and Adamska, 2000). Therefore, based on our results we can speculate that SEP2 is inducible by both light and N starvation.

Among the genes having a central role in the durum wheat-specific network, there were some transcription factors (i.e. DEAD-box ATP-dependent RNA helicase 3[DEAD-box RH3] and the MIKC-type MADS-box transcription factor) as well as some stress responsive genes (i.e. peroxidase and protein detoxification). For example, as well documented, the DEAD-box RNA helicases are involved in RNA metabolism and have important roles in diverse cellular functions (e.g. plant growth and development, and in response to biotic and abiotic stresses (Vashisht and Tuteja, 2006; Li *et al*., 2008; Linder and Jankowsky, 2011; Zhu *et al*., 2015). Recently, Gu *et al*. (2014) demonstrated the relevant role of the chloroplast DEAD-box RH3 on the growth and stress response in *Arabidopsis thaliana*. Interestingly, it is reported that in bread wheat the MIKC-type MADS-box TFs have key roles in plant growth (Ma *et al*., 2017; Li *et al*., 2018); however, even if one of these transcription factors (Traes_5AL_13E2DEC48) was a central node in the durum wheat-specific network, it was down-regulated under N starvation in comparison to the optimal N condition.

Moreover, several studies reported that in wheat, grown in either in field or greenhouse conditions, activities of many enzymes in the antioxidant defense system (i.e. SOD, CAT, GPX, GR, Prx and LOX) are altered to control the oxidative stress induced by other factors and to maintain the balance between ROS production and detoxification which avoid potential damage to cellular components, metabolism, development and growth system (Mittler *et al*., 2004; Caverzan *et al*., 2016 and reference therein). For example, Kumar *et al*. (2013) reported an increase of SOD transcript in wheat in response to heat shock treatment that may indicate greater tolerance to environmental stresses. Also, in this study, many important genes related to the antioxidant defense system were up-regulated in both genotypes but with a ratio of 1:2 between emmer and durum wheat.

The up-regulation of genes involved in the defense-system and the increase in the content of metabolites under starvation observed in durum wheat suggest that a possible mechanism of response to the starvation may be linked to the autophagy. This process is inducible in different and multiple stress condition or development stages, and it is defined as a non-specific degradation process for the recycling of intracellular material that might be used as building blocks to temporarily overcome the absence of nutrients (Liu and Bassham, 2012; Pérez-Pérez *et al*., 2012). Nutrient limitation also increases ROS production, which in turn may stimulate autophagy functioning as signaling molecules as suggested by Liu *et al*. (2009). Taken together, these findings indicate that the absence of nutrients is a primary signal leading to autophagy activation in eukaryotes, but this stress signal is tightly associated with the production and accumulation of ROS. Because in plants the chloroplasts are primary source of ROS, their degradation through autophagic processes may be highly possible as also reported under carbon-limited conditions (Wada *et al*., 2009).

To face environmental constrains, according to the plant-life history (the distribution of resources between growth, reproduction and defense), plants can combine acclimation mechanisms from different strategies defined as escape or resistance (Shaar-Moshe *et al*., 2018 and references therein). In this sense, probably, emmer as adaptive strategy to N starvation relied mainly on the below-ground part while the durum wheat reacts on the up-ground part. Although our experiment did not analyze the transcriptomic and/or metabolomics responses of the roots, it provides important information with respect to differential response on the level of gene and metabolites involving in the efforts of this crop to retain homeostasis under nutrient stress conditions. Indeed, the responses of emmer appear more plastic with enhanced activation of root growth under N starvation then durum which trigger to maintain growth rate even in absence of available N.

## Experimental Procedures

### Plant materials and experimental design

Two genotypes of *Triticum turgidum* were considered: one emmer (*T. turgidum* ssp. *dicoccum*) named ‘Molise Selezione Colli’, a pure line selected from a local population, and one modern durum wheat cultivar (*T. turgidum* ssp. *durum*) named ‘Simeto’ (derived from Capeiti/Valnova), released in ltaly in 1988. They showed many contrasting traits including differences in grain yield (GY), heading date (HD), plant height (PH), test weight (TW), thousand kernel weight (TKW), protein content (PC), yellow index (YI), gluten index (GI), roots and shoot morphological parameters (De Vita *et al*., 2006, 2007; Iannucci *et al*., 2017).

Both genotypes were previously purified by two cycles of Single Seed Descent (SSD). The samples were part of a larger four-week-long experiment conducted in 2012 under N-optimal and N-starvation conditions in the PhyTec Experimental Greenhouse at the Institute of Biosciences and Geosciences (IBG-2): Plant Sciences Institute, Forschungszentrum Julich GmbH, Germany (50°54’36’’N, 06°24’49’’E) which included 12 genotypes for each tetraploid wheat subspecies. The resulting 36 genotypes were grown under two different N conditions with two replicates per genotype in two subsequent growing conditions. Thus, for each treatment genotypes were replicated four times using two plants per replicate with overall 8 plants per genotype per treatment. Each rhizobox contained two different genotypes of the same subspecie, each represented by two plants arranged to avoid contacts between roots of different genotypes. This means that the two genotypes considered here were grown in four different rhizoboxes for each N condition. Before sowing, for each genotype, grains of uniform size were visually selected, surface sterilized (1% NaClO (w/v) for 15 min and rinsed 10 times with deionized water), pre-germinated and then transplanted into the soil-filled rhizoboxes. The soil used to fill the rhizoboxes (volume of ∼18 l) was a ‘Typ 0’ manually sieved peat soil (Nullerde Einheitserde; Balster Einheitserdewerk, Frondenberg, Germany), which provided low nutrient availability, with a pH of 6.1, and the available phosphate, potassium, magnesium, ammonium nitrogen, and nitrate nitrogen concentrations of 7.0, 15.0, 98.0, <1.0, and <1.0 mg l–1, respectively. All plants were watered regularly twice a day with 400 ml of tap water and supplied three times per week with 200 ml of modified Hoagland solution (Hoagland and Arnon, 1950) with or without added nitrogen. For the optimal nitrogen condition, the stock solution included 5 mM KNO_3_, 5 mM Ca(NO_3_)_2_, 2 mM MgSO_4_, 1 mM KH_2_PO_4_, plus trace elements while for the nitrogen-starvation solutions, 1 mM KNO_3_ and 5 mM Ca(NO_3_)_2_ were replaced by 2.5 mM K_2_SO_4_ and 5 mM CaCl_2_6(H_2_O), respectively. The experiments were carried out under natural lighting in a greenhouse, with the air temperature kept between 18 and 24 °C, and the relative humidity between 40 and 60%. For more details concerning the experiment and growth conditions see Gioia *et al*. (2015). At the end of the experiment, for each replicate, leaves of the two plants were pooled and immediately frozen in liquid nitrogen to obtain leaves tissues for RNA and metabolites extraction.

### Phenotypic traits

The following traits were scored for both genotypes: the total leaf area (TLA), the total number of leaves (TLN), and the principal parameters of the root system architecture, such as: visible primary root length (PRL), visible lateral root length (LRL), total root length (TRL) of all visible roots, root system depth (RSD), and root system width (RSW). At the end of the experiment, at 28 days after sowing (DAS) (Zadoks stage 14–18 for optimal N; Zadoks stage 12–14 for N starvation; Zadoks *et al.*, 1974), the chlorophyll content (SPAD units) was estimated with a SPAD-502 chlorophyll meter (Minolta Corp., Ramsey, NJ, USA). In addition, wheat plants were harvested to determine the shoot fresh weight (SFW) and the root biomass (root dry weight; RDW) after a careful washing and oven drying. More details of each determination were reported in Gioia *et al*. (2015).

### Transcriptomic analysis

RNA extraction was performed using 100 mg of frozen ground tissue (leaves) and treated with RNase-Free DNase by the On-Column DNase I Digestion Set (Sigma-Aldrich). For the subsequent analysis only RNA samples with integrity greater than 8.0 were used. Library construction and RNA sequencing were carried out at the Montpellier Genomix (http://www.mgx.cnrs.fr) sequencing facility. Libraries quantification, RNA-Seq data filtering and processing used in this study were essentially as those described previously by David *et al.* (2014). The bread wheat chromosome survey sequence for the cv. Chinese Spring (http://plants.ensembl.org/triticum_aestivum) generated by the International Wheat Genome Sequencing Consortium (IWGSC) was used as the reference assembly. The Biomart package of EnsEMBL were used to acquire the transcripts, and the physical genomic location of the 66,307 genes was predicted from the IWGSC on the genome A and B (Ensembl release 22, http://plants.ensembl.org/biomart/martview/). Since these sequences were obtained by separately sequencing each bread wheat chromosome arm, the bread wheat reference helped to distinguish paralogous durum wheat copies.

RNA-Seq reads were mapped on the bread wheat reference transcriptome using BWA (Li and Durbin, 2009) while allowing 3 errors (-n 3 in the alignment step). Picard tools (http://picard.sourceforge.net) were used to remove PCR and optical duplicates. Rough read counts were computed at all sites for each individual using the idxstats function of the Samtools.

### Metabolite Profiling

After collection, part of the frozen leaves of each replicate were freeze-dried and successively milled using a Pulverisette 7 Planetary Micro Mill (Classic Line, Fritsch) with an agate jar and balls, and stored a −20°C until analysis.

A total of 30 mg dry weight (dw) of each replicate was used for the extraction, derivatisation and analysis by gas chromatography–mass spectrometry (GC-MS) of the polar and non-polar metabolites as previously described (Beleggia *et al*., 2013). Metabolites were identified by comparing the mass spectrometry data with those of a custom library obtained with reference compounds and with those of the National Institute of Standards and Technology (NIST 2011) database. The chromatograms and mass spectra evaluation and quantification were performed using the Mass Hunter software.

The standards and all the chemicals used were HPLC grade (Sigma-Aldrich Chemical Co., Deisenhofen, Germany).

### Statistical analysis

Analysis of variance (ANOVA) was carried out with respect to each morphological trait and metabolite detected in the shoot of emmer and durum wheat lines considered. Mean discrimination between emmer and durum was performed applying Tukey’s test and statistically significant differences were determined at the significance level of *α*= 0.05. Statistical analysis of the data was performed using the JMP software (SAS Institute Inc., Cary, NC, USA version 8).

### Bioinformatics analysis and network construction

#### Data preprocessing

First, genes for which the count per million (cpm) for a single sample was smaller than one and the sum of cpms across all samples was smaller than the total number of samples were filtered out. Raw counts were first normalized using trimmed mean of M-values normalization method (R package edgeR (Robinson *et al*., 2010) and then voom normalized using the R package limma (Smyth, 2005).

#### Analysis of differential expression

Analysis of differential expression was conducted on the data after data preprocessing. DEGs were determined between N starvation and a control with optimal N level for the following scenarios: (*i*) for each genotype and (*ii*) between the two genotypes. For the two scenarios, a linear model was employed to determine differential behavior. To this end, we applied the R package limma (Smyth, 2005).

#### GO enrichment analysis

Annotations were extracted from EnsemblPlants (Kinsella *et al*., 2011) (http://plants.ensembl.org/biomart/martview/2ace56daacae40bad4af00cc25d51e4f) and agriGO (http://bioinfo.cau.edu.cn/agriGO/download.php) (Du *et al*., 2010). We used hypergeometric test (Kachitvichyanukul and Schmeiser, 1985) to identify enriched terms in the list of DEGs. The cut-off value for significance level was considered as 0.05 after FDR correction.

#### Network analysis

Co-expression networks were extracted by applying Pearson correlation on all pairs of data profiles, resulting in a similarity matrix S_m×m_. We then build a network G = (V, E) with m nodes, corresponding to the DEGs; there is an edge between two nodes i, j ∈ V(G) if and only if the entrys_ij_ of S is significant at the level of 0.05, after FDR correction.

Co-expression networks are separately reconstructed for the data from each genotype (*i.e.*, Molise Sel. Colli and Simeto), by identifying significant correlation coefficients between each pair of genes in the network (p value<0.05, FDR corrected). For each pair of genes in the network, the Fisher Z-score test is used to assess the significance of the difference between the correlation coefficients obtained from emmer and durum wheat data. The edge between a pair of genes is referred to as a *significantly different edge* if the obtained p-value from Fisher Z-score test is smaller than 0.05 after FDR correction. The degree and the betweenness centralities of all nodes (i.e. DEGs) in co-expression networks and the differential networks were calculated using the R package igraph (Csardi and Nepusz, 2006). The same analysis was repeated for metabolite data, and integration of metabolite and transcript data; however, the entire metabolite profiles were used in these cases.

To find the nodes (i.e., genes and metabolites) which capture the differences between the two genotypes, we scored the nodes by the number of correlations of value larger than *τ* (*τ* was considered to be 0.6 and 0.8) present in emmer but not in durum wheat co-expression network and vice versa.

## Author Contributions

RB and RP conceived and designed the study. TG, FF, US carried out the experiments and performed the morphological analysis; RB and FN performed the metabolite analyses. YH and JD performed the RNAseq analysis; NO and ZN performed the bioinformatics and network analyses. RB, RP, NO and ZN analyzed data and wrote the paper. NP and PDV reviewed the manuscript. All authors read and approved the final manuscript.

## Acknowledgments

The authors are thankful to Assaf Distelfeld for providing genomic information and for a critical reading of the manuscript. TG thanks the SIR-MIUR grant number [RBSI14SJ81]. NO and ZN acknowledge the support from Horizon 2020 Teaming project PlantaSyst (EU). RP thanks the Università Politecnica delle Marche financial support.

