## Supplementary material for "Comparative analysis based on transcriptomics and metabolomics data reveal differences between emmer and durum wheat in response to nitrogen starvation"

**Figure S1.** Position of down- (red) and up-regulated (blue) DEGs in emmer (Molise) and durum wheat (Simeto) on the physical map. Central DEGs in the transcript-metabolite correlation networks were shown by triangles.

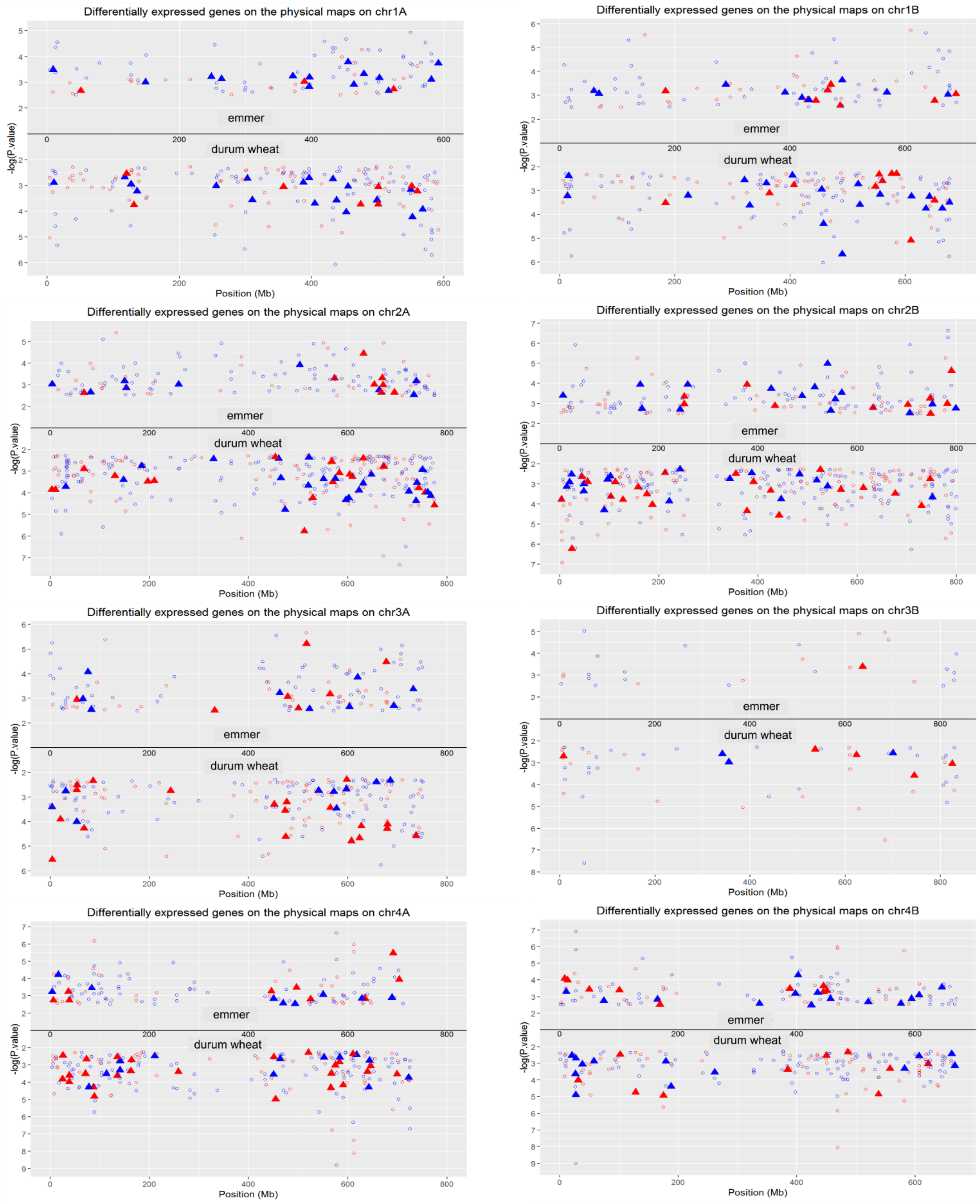

**Table S1** Reads and mapping results

| cie | Genotype | Nitrogen Level | Reads Total Number | Reads Total Number (cleaned) | Reads Mapped | Reads mean | Reads Mapped mean | % |
|---|---|---|---|---|---|---|---|---|
| *Triticum turgidum* ssp. *dicoccum* | Molise | +N | 12,426,327 | 11,729,149 | 8,634,292 | 14,514,858 | 10,578,245 | 72,9 |
| | | | 15,048,113 | 13,952,143 | 10,324,849 | | | |
| | | | 14,903,919 | 14,109,719 | 9,863,795 | | | |
| | | | 19,540,053 | 18,268,422 | 13,490,043 | | | |
| | | -N | 14,549,043 | 13,877,108 | 9,621,513 | 12,123,763 | 8,512,069 | 70,2 |
| | | | 15,385,878 | 14,271,168 | 10,653,558 | | | |
| | | | 10,419,996 | 9,732,724 | 6,506,270 | | | |
| | | | 11,085,787 | 10,614,052 | 7,266,933 | | | |
| [illegible] *turgidum* convar *durum* | Simeto | +N | 11,679,027 | 10,989,390 | 7,657,401 | 12,161,459 | 8,247,526 | 67,8 |
| | | | 14,556,720 | 13,748,679 | 9,989,064 | | | |
| | | | 13,016,891 | 12,037,935 | 8,523,540 | | | |
| | | | 12,548,405 | 11,869,831 | 6,820,097 | | | |
| | | -N | 14,047,180 | 13,176,130 | 9,423,828 | 13,197,161 | 9,291,518 | 70,4 |
| | | | 16,677,090 | 15,519,511 | 10,966,290 | | | |
| | | | 15,471,231 | 14,636,407 | 10,024,095 | | | |
| | | | 9,905,802 | 9,456,597 | 6,751,858 | | | |

**Table S2. Differentially expressed genes in common or specific in emmer (cv. Molise Sel. Colli) and in durum wheat (cv. Simeto) under nitrogen starvation. Key genes involved in nitrogen metabolism, carbon metabolism, transporters, transcription factors, kinases, and stress-related genes are reported. In blue are represented DEGs up regulated and in red DEGs down regulated.**

| N metabolism and key enzymes | Gene ID | Description | common | Molise Sel. Colli | Simeto |
|---|---|---|---|---|---|
| N | Traes_6AS_AB7CDF374 | Nitrate reductase | | | x |
| | Traes_6AS_148C79718 | Nitrate reductase | | | x |
| | Traes_5BL_7E69810D5 | Asparagine synthetase [glutamine-hydrolyzing]1 | x | | |
| | Traes_6AL_0F8A39C3D | Aspartate aminotransferase, chloroplastic | x | | |
| | Traes_3AL_B137CC856 | Aspartate aminotransferase 3, chloroplastic | | | x |
| | Traes_7BL_0029CE4EA | Aspartate aminotransferase | | | x |
| | Traes_6AL_2017727C4 | Glutamine synthetase | | | x |
| | Traes_6BL_51A18A82E | Glutamine synthetase | | | x |
| | Traes_6BL_95C7F7123 | Glutamine synthetase | | | x |
| | Traes_2AL_3F84846F1 | Glutamate dehydrogenase | | | x |
| Amino acids metabolism | Traes_2BL_6552196A1 | 3-isopropylmalate dehydrogenase | | | x |
| | Traes_2AL_B4C264AB8 | 3-isopropylmalate dehydrogenase | x | | |
| | Traes_5AL_0002B669C | Beta-ureidopropionase | | | x |
| | Traes_5BL_AA36BC1E0 | Beta-ureidopropionase | | | x |
| | Traes_2BL_0EF162538 | Agmatine deiminase | | | x |
| | Traes_2BL_2768AE3B1 | Aminomethyltransferase | | | x |
| | Traes_2BL_79F69A456 | Aminomethyltransferase | | | x |
| | Traes_6BS_8C5C8E9E2 | Aminopeptidase | x | | |
| | Traes_6AS_ADCD6AB01 | Aminopeptidase M1 | x | | |
| | Traes_4BL_F15D3655D | Arginine biosynthesis bifunctional protein ArgJ, chloroplastic | x | | |
| | Traes_4AS_641F6B847 | Arginine biosynthesis bifunctional protein ArgJ, chloroplastic | x | | |
| | Traes_1BS_DB2EFB90D | Arginine decarboxylase | x | | |

| | | | | |
|---|---|---|---|---|
| Traes_1BL_708C02E25 | Branched-chain-amino-acid aminotransferase | | x | |
| Traes_2BL_FFF96039E | Branched-chain-amino-acid aminotransferase | | | x |
| Traes_5BL_537388393 | Carbamoyl-phosphate synthase small chain, chloroplastic | | x | |
| Traes_4AL_BB919E3D0 | Cysteine desulfurase 1, chloroplastic | | | x |
| Traes_4AL_45E27280D | Cysteine desulfurase 1, chloroplastic | x | | |
| Traes_3B_3890BB4DE | Cysteine synthase | | | x |
| Traes_3AS_A3FC93C65 | Cysteine synthase | | | x |
| Traes_3B_B8A2DB156 | Cysteine synthase | | | x |
| Traes_3AL_50A1008AB | Cysteine synthase | x | | |
| Traes_5AS_EAE5D853B | Cysteine synthase | x | | |
| Traes_5BS_1AC8D3009 | Cysteine synthase | x | | |
| Traes_7BL_B5491A48C | D-3-phosphoglycerate dehydrogenase | | | x |
| Traes_7AL_95C81BBBF | D-3-phosphoglycerate dehydrogenase | | | x |
| Traes_1AL_5E4EECFBC | Delta-1-pyrroline-5-carboxylate dehydrogenase 12A1, mitochondrial | x | | |
| Traes_7BS_5781FD481 | Dihydroxy-acid dehydratase, chloroplastic | | | x |
| Traes_7BS_36E4AE13A | Dihydroxy-acid dehydratase, chloroplastic | | x | |
| Traes_1BL_91B1BCB59 | Folylpolyglutamate synthase | | | x |
| Traes_4AL_9BCB37710 | Glutamate decarboxylase | x | | |
| Traes_4BS_D94FACCE2 | Glutamate decarboxylase | | | x |
| Traes_3AS_3CB8A9C01 | Glutamate decarboxylase 2 | | | x |
| Traes_2AL_73A59BC32 | Glutamate receptor | | | x |
| Traes_3AS_31089CA19 | Histidine biosynthesis bifunctional protein hisIE, chloroplastic | | | x |
| Traes_4AS_E54C88DDB | Imidazole glycerol phosphate synthase hisHF, chloroplastic | x | | |
| Traes_4BL_CF4451CAA | Imidazole glycerol phosphate synthase hisHF, chloroplastic | x | | |
| Traes_6BS_AE8D595A3 | L-aspartate oxidase, chloroplastic | | | x |
| Traes_2AL_F8F55F0A3 | Methionine aminopeptidase | | | x |
| Traes_5AL_8349DE248 | Ornithine aminotransferase, mitochondrial | | | x |
| Traes_5BL_1D3F1BCC2 | Ornithine aminotransferase, mitochondrial | | | x |
| Traes_6AL_727AEFCD4 | Ornithine carbamoyltransferase, chloroplastic | x | | |
| Traes_6BL_0A6B5575F | Ornithine carbamoyltransferase, chloroplastic | x | | |
| Traes_6AS_0C1D497EA | S-adenosylmethionine synthase 4 | | | x |

| | | | | | |
|---|---|---|---|---|---|
| | Traes_6BL_93357D848 | Serine/threonine-protein phosphatase | | | x |
| | Traes_7AS_F3CC01C3D | Serine--glyoxylate aminotransferase | | x | |
| | Traes_7BL_28E519867 | Tryptophan synthase | x | | |
| | Traes_7AL_7C0CB1D06 | Tryptophan synthase | | x | |
| Purine/pyrimidine metabolism | Traes_2AL_DB64E18A1 | Allantoinase | x | | |
| | Traes_2BL_55D540224 | Allantoinase | | | x |
| | Traes_6BL_C433D70A3 | Dihydropyrimidine dehydrogenase (NADP(+)), chloroplastic | | x | |
| | Traes_5AS_BA57B5B56 | Ureidoglycolate hydrolase | | | x |
| | Traes_5BS_E660355D6 | Ureidoglycolate hydrolase | | | x |
| | Traes_3AL_690595C9D | CTP synthase | x | | |
| | Traes_1BL_D633DA4E8 | CTP synthase | x | | |
| | Traes_1AL_B68AA4C68 | CTP synthase | | x | |
| | Traes_1AL_341E2A7A8 | CTP synthase | | | x |
| | Traes_1BL_3A901257A | CTP synthase | | | x |
| | Traes_3AL_2A172A963 | CTP synthase | | | x |
| | Traes_5AL_6FB526190 | Carbonic anhydrase | | x | |
| | Traes_3AL_CAE12F07B | Phosphotransferase | x | | |

| **Transporter** | **Gene ID** | **Description** | **common** | **Molise Sel. Colli** | **Simeto** |
|---|---|---|---|---|---|
| Ammonium transporter | Traes_2AL_CF862CF2D | Ammonium transporter 1 member 4 (AMT1-4) | | x | |
| | Traes_2BL_F1A254464 | Ammonium transporter 1 member 4 (AMT1-4) | | x | |
| Nitrate transporter | Traes_7AS_9D057AD42 | Protein NRT1/ PTR FAMILY 3.1 | | x | |
| Potassium transporter | Traes_1AL_B7757AFDC | Potassium channel AKT2/3 | | x | |
| | Traes_5BL_F112FA40E | Potassium transporter | x | | |
| | Traes_7AS_9A01E6B17 | Potassium transporter | | x | |
| | Traes_3AL_9E0B84E84 | Potassium transporter | | | x |
| | Traes_7BS_E3BDDBFEF | Potassium transporter | | | x |

| | | | | | |
|---|---|---|---|---|---|
| | Traes_5AL_B64648FE6 | Putative potassium transporter 12 | x | | |
| | Traes_5BL_F43BDBCB9 | Putative potassium transporter 12 | | | x |
| | Traes_5AL_91132361F | K(+) efflux antiporter 5 | | | x |
| Phosphate transporter | Traes_7AS_735085EFB | Phosphate transporter PHO1 homolog 1 | | | x |
| Zinc transporter | Traes_2AL_3983FD077 | Zinc transporter ZIP | | | x |
| Sulfate transporter | Traes_3AL_224FB10D3 | Probable sulfate transporter 3.5 | x | | |
| Magnesium transporter | Traes_1AL_C04849EE3 | Probable magnesium transporter | | | x |
| | Traes_3AL_58B8BF68F | Probable magnesium transporter | | | x |
| | Traes_4AL_3C35261D2 | Magnesium/proton exchanger | | | x |
| Sugar transporter | Traes_1AL_039C1D96D | Bidirectional sugar transporter SWEET | | | x |
| | Traes_3AL_5A716129C | Bidirectional sugar transporter SWEET | | | x |
| | Traes_3AL_729C392FB | Bidirectional sugar transporter SWEET | x | | |
| | Traes_4AS_6D625D7BF | Bidirectional sugar transporter SWEET | x | | |
| | Traes_6AL_58B6B7319 | Bidirectional sugar transporter SWEET | | | x |
| | Traes_6BL_BE49FDA0B | Bidirectional sugar transporter SWEET | | | x |
| | Traes_4AS_A322DBCB0 | Sucrose transporter SUT1A | x | | |
| | Traes_4AS_C4001163C | GDP-mannose transporter GONST3 | | | x |
| Calcium transporter | Traes_1BL_9772C6F45 | Calcium-transporting ATPase | | | x |
| | Traes_2BL_90036AF45 | Calcium-transporting ATPase | x | | |
| | Traes_2BL_E2DCFA60B | Calcium-transporting ATPase | | x | |
| | Traes_4AL_8CAEF3C85 | Calcium-transporting ATPase | | x | |
| | Traes_5AL_FA3033A40 | Calcium-transporting ATPase | | | x |
| | Traes_6AS_9B913DF6F | Calcium-transporting ATPase | | x | |
| | Traes_1BL_9772C6F45 | Calcium-transporting ATPase | | | x |
| | Traes_5AL_FA3033A40 | Calcium-transporting ATPase | | | x |

| | | | | | |
|---|---|---|---|---|---|
| Sodium transporter | Traes_3AL_9F1E0BAE1 | Sodium/pyruvate cotransporter BASS2, chloroplastic | | x | |
| Copper transporter | Traes_2AL_D0EABF355 | Probable copper-transporting ATPase HMA5 | | | x |
| | Traes_2BL_19B3E60AA | Probable copper-transporting ATPase HMA5 | | | x |
| ABC transporter | Traes_3AS_BB185F2EB | ABC transporter G family member 7 | x | | |
| | Traes_3B_5FDDEC00B | ABC transporter G family member 32 | | x | |
| | Traes_5AS_C606A8A1F | PDR-type ABC transporter | | | x |
| | Traes_3B_77C6EE076 | ABC transporter G family member 7 | | | x |
| | Traes_4AS_6A57D15D9 | ABC transporter I family member 6, chloroplastic | | | x |
| | Traes_3AS_E18F065A3 | Protein ABCI7, chloroplastic | | | x |
| Amino acid transporter | Traes_6AL_1E68C6606 | Cationic amino acid transporter 9, chloroplastic | x | | |
| | Traes_4AL_1B3E7B791 | WAT1-related protein | | x | |
| | Traes_5BL_CBB9D7E7F | WAT1-related protein | | | x |
| Other transporter | Traes_2AL_2612F72B9 | 3-oxoacyl-[acyl-carrier-protein] synthase | x | | |
| | Traes_2AL_341C03FBF | 3-oxoacyl-[acyl-carrier-protein] synthase | x | | |
| | Traes_2AL_5745581C6 | 3-oxoacyl-[acyl-carrier-protein] synthase | x | | |
| | Traes_1BS_8C10A5D1D | Acyl carrier protein (ACP) | | | x |
| | Traes_5AL_D9BFBD63A | Acyl carrier protein (ACP) | x | | |
| | Traes_5BL_E9C760122 | Acyl carrier protein (ACP) | x | | |
| | Traes_5AS_178DFC4E3 | Acyl carrier protein (ACP) | x | | |
| | Traes_5BS_4C0AD2649 | Acyl carrier protein (ACP) | x | | |
| | Traes_2AL_E80950527 | Chloride channel protein | | | x |
| | Traes_5BL_B8C3AE4A5 | Chloride channel protein | | | x |
| | Traes_6AS_0F6151540 | Chloride channel protein | | | x |
| | Traes_6AS_28752C23C | Chloride channel protein | | | x |
| | Traes_2BL_1356B2300 | Chloride channel protein | x | | |
| | Traes_4AL_6240D6D66 | Folate transporter 1, chloroplastic | | | x |

| | | | | |
|---|---|---|---|---|
| Traes_2BL_E2C38B73C | Gamma-soluble NSF attachment protein | | | x |
| Traes_6BL_2B67C5E30 | Glycolipid transfer protein 1 | | | x |
| Traes_6BL_2B67C5E30 | Glycolipid transfer protein 1 | | | x |
| Traes_7BL_C46BC291C | Heavy metal transporting P1B-ATPase 2 | | x | |
| Traes_7AL_8304348B7 | Heavy metal transporting P1B-ATPase 2 | x | | |
| Traes_5BL_68C95B396 | Mitochondrial import inner membrane translocase subunit TIM10 | | x | |
| Traes_1BS_0A858EB86 | Mitochondrial pyruvate carrier | | | x |
| Traes_1BS_0A858EB86 | Mitochondrial pyruvate carrier | | | x |
| Traes_4AS_5828A4E4D | Peroxisomal nicotinamide adenine dinucleotide carrier | | | x |
| Traes_2AS_F3FE36E3E | Phospholipid-transporting ATPase | | | x |
| Traes_2BS_2317C066E | Phospholipid-transporting ATPase | | | x |
| Traes_4AS_D950EBDB0 | Phospholipid-transporting ATPase | | | x |
| Traes_4BL_CA03BD203 | Phospholipid-transporting ATPase | | | x |
| Traes_4AL_D7702A458 | Plasma membrane ATPase | | | x |
| Traes_5BS_7DDBE5B89 | Plasma membrane ATPase | | x | |
| Traes_4AL_D7702A458 | Plasma membrane ATPase | | | x |
| Traes_1AL_C544E6828 | Plastidal glycolate/glycerate translocator 1, chloroplastic | | | x |
| Traes_2AL_E7D1C8AA5 | Probable inositol transporter 2 | | | x |
| Traes_2BL_DE4FAAC5E | Probable inositol transporter 2 | | | x |
| Traes_2AL_7F21D24BA | Protein TIC 40, chloroplastic | | | x |
| Traes_2BL_A4FF3C90C | Protein TIC 40, chloroplastic | | | x |
| Traes_5BL_28CCE5325 | Protein TIC 62, chloroplastic | | | x |
| Traes_7BL_95C7C23DD | Sec-independent protein translocase protein TATB, chloroplastic | | x | |
| Traes_1BL_5F74C58ED | Secretory carrier-associated membrane protein (SCAMPs) | | x | |
| Traes_1AL_090571678 | Secretory carrier-associated membrane protein (SCAMPs) | | x | |
| Traes_3AL_697247353 | Secretory carrier-associated membrane protein (SCAMPs) | | | x |
| Traes_7AS_D1A84E231 | Trigger factor-like protein TIG, Chloroplastic | x | | |
| Traes_7BS_B18D7717E | Trigger factor-like protein TIG, Chloroplastic | x | | |
| Traes_5BL_2C3037F54 | Vacuolar protein sorting-associated protein 2 homolog 2 | x | | |
| Traes_6AL_2C61320D2 | Photosynthetic NDH subunit of lumenal location 4, chloroplastic | | x | |

| | Traes_4BL_F37C90CB8 | Photosynthetic NDH subunit of subcomplex B 2, chloroplastic | | x | |
|---|---|---|---|---|---|
| **C metabolism** | **Gene ID** | **Description** | **common** | **Molise Sel. Colli** | **Simeto** |
| Glycolisis | Traes_2AS_D9D9A9ADB | [Pyruvate dehydrogenase (acetyl-transferring)] kinase, mitochondrial | | x | |
| | Traes_6BS_C9B216D97 | Acetyltransferase component of pyruvate dehydrogenase complex | | x | |
| | Traes_2AL_2469A32A8 | Aldose 1-epimerase | | | x |
| | Traes_2AL_CA75501A81 | Aldose 1-epimerase | | | x |
| | Traes_2AL_729CB1040 | Aldose 1-epimerase | | | x |
| | Traes_5AS_C7C69D278 | ATP-dependent 6-phosphofructokinase | X | | |
| | Traes_4AL_EBD5A433B | ATP-dependent 6-phosphofructokinase (PFK1) | | x | |
| | Traes_4AL_EBD5A433B | ATP-dependent 6-phosphofructokinase (PFK1) | | x | |
| | Traes_5BL_DECE49DFC | Dihydrolipoamide acetyltransferase component of pyruvate dehydrogenase complex | | | x |
| | Traes_5AL_17458C47B | Dihydrolipoamide acetyltransferase component of pyruvate dehydrogenase complex | | | x |
| | Traes_5AS_3B966EA46 | Dihydrolipoamide acetyltransferase component of pyruvate dehydrogenase complex | x | | |
| | Traes_5AL_32B5C730F | Dihydrolipoamide acetyltransferase component of pyruvate dehydrogenase complex | x | | |
| | Traes_5BL_BA19E1CE3 | Dihydrolipoamide acetyltransferase component of pyruvate dehydrogenase complex | x | | |
| | Traes_1BS_15C828137 | Dihydrolipoyl dehydrogenase | | | x |
| | Traes_5AL_AB5831D39 | Enolase 1, chloroplastic | | x | |
| | Traes_5BL_B4537ED84 | Enolase 1, chloroplastic | | | x |
| | Traes_4BS_D12DBE6D3 | Fructose-bisphosphate aldolase | | | x |
| | Traes_3AL_441C0AE1B | Fructose-bisphosphate aldolase | | | x |
| | Traes_5BS_017F8702A | Fructose-bisphosphate aldolase | | | x |
| | Traes_4BS_4404098F1 | Fructose-1,6-bisphosphatase, chloroplastic | | | x |
| | Traes_2AL_783CF383F | Glyceraldehyde-3-phosphate dehydrogenase | | | x |
| | Traes_2BL_5D64E8C87 | Glyceraldehyde-3-phosphate dehydrogenase | | | x |
| | Traes_4BL_F32809B15 | Glyceraldehyde-3-phosphate dehydrogenase | | | x |
| | Traes_4BL_A9FAA75A9 | Glyceraldehyde-3-phosphate dehydrogenase | | | x |
| | Traes_2AL_783CF383F | Glyceraldehyde-3-phosphate dehydrogenase | | | x |

| | | | | | |
|---|---|---|---|---|---|
| | Traes_2BL_5D64E8C87 | Glyceraldehyde-3-phosphate dehydrogenase | | | x |
| | Traes_4BL_F32809B15 | Glyceraldehyde-3-phosphate dehydrogenase | | | x |
| | Traes_4BL_A9FAA75A9 | Glyceraldehyde-3-phosphate dehydrogenase | | | x |
| | Traes_1BL_F7884B670 | Phosphoglycerate kinase (PGK) | | | x |
| | Traes_1BL_F7884B670 | Phosphoglycerate kinase (PGK) | | | x |
| | Traes_2BS_167A31F21 | Plastidial pyruvate kinase 1, chloroplastic | | | x |
| | Traes_3B_631CEF092 | Plastidial pyruvate kinase 2 | | | x |
| | Traes_3AL_81435CD25 | Plastidial pyruvate kinase 2 | x | | |
| | Traes_5AS_4C84934FE | Pyrophosphate--fructose 6-phosphate 1-phosphotransferase subunit alpha | | x | |
| | Traes_7AS_25F9FEF52 | Pyruvate dehydrogenase E1 component subunit alpha | | x | |
| | Traes_2AS_F485758C1 | Pyruvate dehydrogenase E1 component subunit alpha-3, chloroplastic | x | | |
| | Traes_2BL_9B9195C2E | Pyruvate kinase | | | x |
| | Traes_5AL_89F608111 | Pyruvate kinase | | | x |
| | Traes_5BL_6E109E397 | Pyruvate kinase | | | x |
| | Traes_2AL_D6D188AA7 | Pyruvate kinase | | | x |
| | Traes_2BL_9B9195C2E | Pyruvate kinase | | | x |
| | Traes_5AL_89F608111 | Pyruvate kinase | | | x |
| | Traes_5BL_6E109E397 | Pyruvate kinase | | | x |
| | Traes_2AL_D6D188AA7 | Pyruvate kinase | | | x |
| | Traes_1AL_E5E41283E | Pyruvate, phosphate dikinase 1, chloroplastic | x | | |
| | Traes_1BL_AAC4D956E | Pyruvate, phosphate dikinase 1, chloroplastic | x | | |
| Pentose phosphate shunt | Traes_2AS_67222868A | Glucose-6-phosphate 1-dehydrogenase | x | | |
| | Traes_5AL_C14B0076E | Probable ribose-5-phosphate isomerase 4, chloroplastic | | | x |
| | Traes_5BL_063A12847 | Probable ribose-5-phosphate isomerase 4, chloroplastic | | | x |
| Tricarboxilic acid cycle (TCA) | Traes_2BL_CEC7F2D7B | Isocitrate dehydrogenase [NADP] | | | x |
| | Traes_2AL_3668C746C | D-2-hydroxyglutarate dehydrogenase, mitochondrial | | | x |
| | Traes_2BL_A42EAB22C | D-2-hydroxyglutarate dehydrogenase, mitochondrial | | | x |

| | | | | | |
|---|---|---|---|---|---|
| Photosynthesis | Traes_5AL_7E134CAB9 | [Fructose-bisphosphate aldolase]-lysine N-methyltransferase, chloroplastic | | | x |
| | Traes_5BL_3CD624CE9 | [Fructose-bisphosphate aldolase]-lysine N-methyltransferase, chloroplastic | | | x |
| | Traes_4AL_70D393C3B | 2-carboxy-1,4-naphthoquinone phytyltransferase, chloroplastic | x | | |
| | Traes_2BS_4AE914BE2 | Chlorophyll a-b binding protein, chloroplastic | | | x |
| | Traes_5BL_2C06592B7 | Chlorophyll a-b binding protein, chloroplastic | | | x |
| | Traes_5BL_5081CE9EC | Chlorophyll a-b binding protein, chloroplastic | | | x |
| | Traes_7AS_C05C7D06B1 | Chlorophyll a-b binding protein, chloroplastic | | | x |
| | Traes_7BS_91EA4C6E5 | Chlorophyll a-b binding protein, chloroplastic | | | x |
| | Traes_2BS_F6A5E8996 | Chlorophyll a-b binding protein, chloroplastic | | | x |
| | Traes_5AL_75780E5B0 | Chlorophyll a-b binding protein, chloroplastic | | x | |
| | Traes_1AL_053F6B12B | Chlorophyll synthase, chloroplastic | x | | |
| | Traes_1AL_1A98A80B4 | Chlorophyll synthase, chloroplastic | | | x |
| | Traes_6AS_E114D6D5B | Delta-aminolevulinic acid dehydratase | x | | |
| | Traes_6BS_0726A0A88 | Delta-aminolevulinic acid dehydratase | x | | |
| | Traes_6AS_F205EE33F | Delta-aminolevulinic acid dehydratase | x | | |
| | Traes_7AL_3C614133C | Delta-aminolevulinic acid dehydratase | x | | |
| | Traes_7BL_C35CD97E7 | Delta-aminolevulinic acid dehydratase | x | | |
| | Traes_7AL_7C99238B0 | Ferredoxin | | | x |
| | Traes_2BL_363CA4ED8 | Ferredoxin-3 | | | x |
| | Traes_2AL_688678052 | Ferredoxin-3 | | | x |
| | Traes_5AL_B0E91B780 | Ferredoxin--NADP reductase | x | | |
| | Traes_5AL_B0E91B780 | Ferredoxin--NADP reductase | x | | |
| | Traes_5AL_23B24917C | Ferrochelatase | | | x |
| | Traes_5BL_FF9AA4B58 | Ferrochelatase | | | x |
| | Traes_5AL_3EED2020C | Magnesium-chelatase subunit ChlD, chloroplastic | | | x |
| | Traes_5AL_F490CE585 | Magnesium-chelatase subunit ChlD, chloroplastic | x | | |
| | Traes_5BL_1DE088A30 | Magnesium-chelatase subunit ChlD, chloroplastic | x | | |
| | Traes_5BL_CFC7BB483 | Magnesium-chelatase subunit ChlD, chloroplastic | x | | |
| | Traes_7BL_82850CC15 | Mg-protoporphyrin IX chelatase | | | x |
| | Traes_7AL_8481A6FD7 | Mg-protoporphyrin IX chelatase | x | | |

| | | | | | |
|---|---|---|---|---|---|
| | Traes_7BL_12EBCA739 | Mg-protoporphyrin IX chelatase | x | | |
| | Traes_3AL_6BDD6F090 | NAD(P)H-quinone oxidoreductase subunit O, chloroplastic | | | x |
| | Traes_5AL_3C4AACD91 | Probable zinc metalloprotease EGY1, chloroplastic | x | | |
| | Traes_3AS_922982272 | Probable zinc metalloprotease EGY2 | x | | |
| | Traes_5AS_3354E01BA | Protoporphyrinogen oxidase 1, chloroplastic (PPOX1) | x | | |
| | Traes_5BS_5C3BA5328 | Protoporphyrinogen oxidase 1, chloroplastic (PPOX1) | x | | |
| | Traes_2AL_1AD2DDF83 | Protoporphyrinogen oxidase 2, chloroplastic/mitochondrial | | | x |
| | Traes_5BS_69F251543 | PsbP domain-containing protein 1, chloroplastic | X | | |
| | Traes_2BL_ACD272701 | PsbP domain-containing protein 2, chloroplastic | X | | |
| | Traes_2AL_57B2ACCC0 | PsbP domain-containing protein 2, chloroplastic | | | x |
| | Traes_6AS_FD6AE4DD4 | Red chlorophyll catabolite reductase, chloroplastic | | | x |
| | Traes_2AS_2B3603918 | Ribulose bisphosphate carboxylase small chain (RUBISCO) | | | x |
| | Traes_2AS_C3EF3A27F | Ribulose bisphosphate carboxylase small chain (RUBISCO) | | | x |
| | Traes_2AS_DDDAF2E70 | Ribulose bisphosphate carboxylase small chain (RUBISCO) | | | x |
| | Traes_2BS_0AF61748C | Ribulose bisphosphate carboxylase small chain (RUBISCO) | | | x |
| | Traes_2BS_E68F1B47C | Ribulose bisphosphate carboxylase small chain (RUBISCO) | | | x |
| | Traes_2AS_7425ED39F | Ribulose bisphosphate carboxylase small chain (RUBISCO) | | | x |
| | Traes_3AL_D2C3599B4 | Uroporphyrinogen decarboxylase (UROD) | x | | |
| | Traes_4BL_B902E85F5 | Uroporphyrinogen decarboxylase 2, chloroplastic (UROD2) | x | | |
| | Traes_4AS_90CC29CAA | Uroporphyrinogen decarboxylase 2, chloroplastic (UROD2) | | | x |
| Photorespiration | Traes_7AS_95F4E4729 | Carboxyl-terminal-processing peptidase 3, chloroplastic | x | | |
| | Traes_7BS_462DD3C32 | Carboxyl-terminal-processing peptidase 3, chloroplastic | x | | |
| | Traes_7BS_7E9C070B8 | Carboxyl-terminal-processing peptidase 3, chloroplastic | x | | |
| | Traes_1AL_C544E6828 | Plastidal glycolate/glycerate translocator 1, chloroplastic | | | x |
| Fatty acids metabolism | Traes_4AS_E1B651375 | 3-ketoacyl-CoA synthase | | x | |
| | Traes_4BL_A71648154 | 3-ketoacyl-CoA synthase 10 | | | x |
| | Traes_4AS_D58530D14 | 3-ketoacyl-CoA synthase 10 | | | x |
| | Traes_5AL_4ACF1AEBB | Acyl-[acyl-carrier-protein] desaturase | x | | |
| | Traes_5BL_3CEA3D2B4 | Acyl-[acyl-carrier-protein] hydrolase | | x | |
| | Traes_5AL_CEEDF43FC | Acyl-[acyl-carrier-protein] hydrolase | | x | |

| | | | | | |
|---|---|---|---|---|---|
| | Traes_3AS_0EA847466 | Fatty acyl-CoA reductase | | | x |
| | Traes_5AL_DC104F3FE | Fatty acyl-CoA reductase | | | x |
| | Traes_7BS_2DA5DB033 | Fatty acyl-CoA reductase | | | x |
| | Traes_4BS_DA955986B | Fatty acyl-CoA reductase | | | x |
| | Traes_4BS_DA955986B | Fatty acyl-CoA reductase | | | x |
| | Traes_6AL_0C5F4D84E | Fatty-acid-binding protein 2 | | | x |
| | Traes_2AL_F1C4FF20C | Lecithin-cholesterol acyltransferase-like 4 | x | | |
| | Traes_2BL_3DBAA0726 | Lecithin-cholesterol acyltransferase-like 4 | | | x |
| | Traes_2BL_3DBAA0726 | Lecithin-cholesterol acyltransferase-like 4 | | | x |
| | Traes_1AS_522839777 | Long chain acyl-CoA synthetase 1 (LCAS 1) | x | | |
| | Traes_1AS_0EAA8A0B4 | Long chain acyl-CoA synthetase 1 | | x | |
| | Traes_1BS_3C86AE038 | Long chain acyl-CoA synthetase 1 | | x | |
| | Traes_5AL_3803F504A | Long chain acyl-CoA synthetase 8 | | | x |
| | Traes_2AS_2980E0300 | Omega-6 fatty acid desaturase, chloroplastic | | | x |
| | Traes_4AS_D2A85E4CA | Patatin | | x | |
| | Traes_7AL_CB50E0F46 | Patatin | x | | |
| | Traes_2BL_D1BE1544E | Protein S-acyltransferase 21 | | x | |
| | Traes_6BL_D1538B2A7 | Very-long-chain (3R)-3-hydroxyacyl-CoA dehydratase | | | x |
| Other metabolism | Traes_7AL_1E9FBDD47 | 1,4-alpha-glucan-branching enzyme 3, chloroplastic/amyloplastic | x | | |
| | Traes_7AL_5D8D7CB60 | 1,4-alpha-glucan-branching enzyme 3, chloroplastic/amyloplastic | | | x |
| | Traes_3AL_A3B273BAC | 1,4-dihydroxy-2-naphthoyl-CoA synthase, peroxisomal | | | x |
| | Traes_2AL_73EA196D5 | 2-phytyl-1,4-beta-naphthoquinone methyltransferase, chloroplastic | | | x |
| | Traes_3AS_6F9355B92 | 3-epi-6-deoxocathasterone 23-monooxygenase | | | x |
| | Traes_2BS_8D01B66C0 | 4-alpha-glucanotransferase DPE1, chloroplastic/amyloplastic (maltotriose metabolism) | x | | |
| | Traes_2AS_3C5CB1C2C | 4-alpha-glucanotransferase DPE1, chloroplastic/amyloplastic (maltotriose metabolism) | x | | |
| | Traes_3AL_8C6F4F66A | 4-diphosphocytidyl-2-C-methyl-D-erythritol kinase, chloroplastic | x | | |
| | Traes_4AS_80A9FCCF4 | Alpha-1,3-glucosyltransferase | | | x |

| | | | | |
|---|---|---|---|---|
| Traes_1AL_ED72E167A | Alpha-galactosidase | | x | |
| Traes_2AS_65297DABA | Alpha-galactosidase 3 | x | | |
| Traes_5AL_ED21C722B | Alpha-glucan phosphorylase 1 | | x | |
| Traes_5AL_D86A878AF | Alpha-mannosidase | x | | |
| Traes_5BL_BF60CBEDA | Alpha-mannosidase | x | | |
| Traes_2AS_E0AFA6D18 | ATP sulfurylase 2 | | | x |
| Traes_2BS_F70FE410B | ATP sulfurylase 2 | | | x |
| Traes_3B_55C9357A4 | Beta-amylase | | | x |
| Traes_2AS_61CA61AD3 | Beta-amylase | x | | |
| Traes_2BS_A1909D302 | Beta-amylase | x | | |
| Traes_7AL_AFC2A9D5D | Beta-1,6-galactosyltransferase GALT31A | | x | |
| Traes_4AS_4C25DF6F4 | Beta-galactosidase | | x | |
| Traes_4AS_BB59116B6 | Beta-galactosidase 8 | | x | |
| Traes_1BL_C955569B8 | Beta-hexosaminidase | x | | |
| Traes_1BS_014C64CC2 | Beta-hexosaminidase 1 | | x | |
| Traes_1AL_8E8E7401B | Beta-hexosaminidase | | x | |
| Traes_5AL_05D58A0C4 | Cellulose synthase A catalytic subunit 7 | | | x |
| Traes_3AL_FFE4C92D8 | Cellulose synthase A catalytic subunit 8 [UDP-forming] | | | x |
| Traes_5BL_E86097AA2 | Chalcone-flavonone isomerase family protein | | | x |
| Traes_6BL_3C04C3D2F | Chlorophyll(ide) b reductase NOL, chloroplastic | | | x |
| Traes_6BL_3C04C3D2F | Chlorophyll(ide) b reductase NOL, chloroplastic | | | x |
| Traes_5BS_012B7F86A | Cyclic pyranopterin monophosphate synthase, mitochondrial | | | x |
| Traes_4AL_BB919E3D0 | Cysteine desulfurase 1, chloroplastic | | | x |
| Traes_3AL_883A4805C | Cytokinin riboside 5'-monophosphate phosphoribohydrolase | | x | |
| Traes_7BL_EA8DE756D | Flavin-containing monooxygenase | | | x |
| Traes_2AL_06EF31003 | Galacturonokinase | | x | |
| Traes_2BL_4DDAB77A8 | Galacturonokinase | | x | |
| Traes_7BS_4FBE4B00A | Glucose-1-phosphate adenylyltransferase | x | | |
| Traes_3AL_F3125B233 | Glycerol-3-phosphate dehydrogenase [NAD(+)] 1, chloroplastic | | x | |
| Traes_3AL_12CDB0C85 | Glycerol-3-phosphate dehydrogenase [NAD(+)] | | x | |
| Traes_3AL_30294D0A5 | Glycerol-3-phosphate dehydrogenase [NAD(+)] | | x | |

| | | | | |
|---|---|---|---|---|
| Traes_6AS_626A3D38C | Glycosyltransferase | | x | |
| Traes_5BL_BD269CC23 | Glycosyltransferase | | | x |
| Traes_3AL_829F70144 | Glycosyltransferase | x | | |
| Traes_3AL_A0C2D775D | Glycosyltransferase | x | | |
| Traes_5BL_406A1C48D | Glycosyltransferase | | | x |
| Traes_2AL_06AD35739 | Granule-bound starch synthase 1, chloroplastic/amyloplastic | | | x |
| Traes_2BL_25DAD57C9 | Granule-bound starch synthase 1, chloroplastic/amyloplastic | | | x |
| Traes_2BL_4470DF166 | Granule-bound starch synthase 1, chloroplastic/amyloplastic | | | x |
| Traes_7AL_3F7F0AA7B | Granule-bound starch synthase 1, chloroplastic/amyloplastic | | | x |
| Traes_2AL_BE5540B1E | Malate synthase (pyruvate metabolism) | | | x |
| Traes_3AL_C0D1C4403 | Malic enzyme (pyruvate metabolism) | | | x |
| Traes_5AL_C1E3FCB4F | Monogalactosyldiacylglycerol synthase 1, chloroplastic | | | x |
| Traes_2AS_DEDC612AE | Nicotianamine synthase | | x | |
| Traes_5BS_E7ADA47A4 | Pectinesterase | x | | |
| Traes_1AL_E9BA105D1 | Pectinesterase 31 | | | x |
| Traes_5AS_F24765BD1 | Pectinesterase | | x | |
| Traes_4BL_2465F30E0 | Pheophorbide a oxygenase, chloroplastic | x | | |
| Traes_4BL_4447E55C5 | Pheophorbide a oxygenase, chloroplastic | x | | |
| Traes_7AL_72A908F12 | Pheophytinase, chloroplastic | x | | |
| Traes_1BS_4A63D1DBC | Phosphatidate cytidylyltransferase | | | x |
| Traes_1AS_9D4506E73 | Phosphatidate cytidylyltransferase | | | x |
| Traes_1BS_4A63D1DBC | Phosphatidate cytidylyltransferase | | | x |
| Traes_1AS_9D4506E73 | Phosphatidate cytidylyltransferase | | | x |
| Traes_5AL_636F300F6 | Phosphoglucan phosphatase LSF2, chloroplastic | x | | |
| Traes_2AL_81CAF6C30 | Phosphomannomutase | x | | |
| Traes_4BL_D08D5FFB5 | Phosphomannomutase | x | | |
| Traes_2BL_EB792A35E | Phosphomannomutase | | | x |
| Traes_6BS_A80520954 | Phosphopantetheine adenylyltransferase | | | x |
| Traes_2AL_62160B4F7 | Probable alpha-amylase 2 | | | x |
| Traes_7AL_6A30E603B | Probable galacturonosyltransferase 3 | | x | |
| Traes_7BS_092F4241C | Pullulanase 1, chloroplastic | | x | |
| Traes_5AS_7E93CEA52 | Purple acid phosphatase (PAP) | | x | |

| | | | | | |
|---|---|---|---|---|---|
| | Traes_5BL_6D2126519 | Purple acid phosphatase (PAP) | x | | |
| | Traes_5AS_D42C83AB8 | Purple acid phosphatase (PAP) | | | x |
| | Traes_4BL_840A609C1 | Purple acid phosphatase (PAP) | | | x |
| | Traes_4BL_598C0826B | Purple acid phosphatase (PAP) | | | x |
| | Traes_4BL_99574CDAC | Purple acid phosphatase (PAP) | | | x |
| | Traes_5AL_394CDB9BC | Purple acid phosphatase (PAP) | | | x |
| | Traes_7AL_9BC56072C | Putative beta-glucosidase 41 | | | x |
| | Traes_7AL_AA28D70BF | Starch branching enzyme I | x | | |
| | Traes_7AS_A80BE362A | Sucrose synthase | | x | |
| | Traes_4AL_FE0C5AEF4 | Sucrose:fructan 6-fructosyltransferase | | | x |
| | Traes_2BS_8800ED4EE | Terpene cyclase/mutase family member | | | x |
| | Traes_7AS_E82D29D97 | Terpene cyclase/mutase family member | | | x |
| | Traes_6AS_F3CC61A36 | Terpene cyclase/mutase family member | | x | |
| | Traes_1AL_0FB80B128 | Uracil phosphoribosyltransferase, chloroplastic | x | | |
| | Traes_6AL_006779D22 | Xyloglucan endotransglucosylase/hydrolase | | x | |
| | Traes_4BS_64FB912EE | Xyloglucan endotransglucosylase/hydrolase | x | | |
| | Traes_7AL_1B1FBCDE4 | Xyloglucan endotransglucosylase/hydrolase (XTHs) | | | x |
| | Traes_4AL_320703FD2 | Xyloglucan endotransglucosylase/hydrolase (XTHs) | | | x |
| | Traes_7AL_8C4A9BEBF | Xyloglucan endotransglucosylase/hydrolase (XTHs) | | | x |
| | Traes_2BS_D4E86AAA6 | Xylose isomerase | | x | |
| | Traes_7AL_AFA2C29C9 | Photosystem II stability/assembly factor HCF136, chloroplastic | | | x |
| | Traes_7BL_CA98655DC | Photosystem II stability/assembly factor HCF136, chloroplastic | | | x |
| **Transcription factors** | **Gene ID** | **Description** | **common** | **Molise Sel. Colli** | **Simeto** |
| | Traes_3AL_7A2CED8E7 | Auxin response factor (ARFs Family) | x | | |
| | Traes_3AL_E34DF2F08 | Auxin response factor (ARFs Family) | x | | |
| | Traes_3AL_5935773EA | Auxin response factor (ARFs Family) | | | x |
| | Traes_1BL_A9AC1340E | Auxin response factor (ARFs Family) | | | x |
| | Traes_1AL_147CF243C | Auxin response factor (ARFs Family) | | | x |
| | Traes_5BL_C86AC392A | Auxin-responsive protein | | x | |
| | Traes_5BL_9301BD154 | Auxin-responsive protein | | | x |

| | | | | |
|---|---|---|---|---|
| Traes_5BL_EC006AD0C | Auxin-responsive protein | | | x |
| Traes_4BL_7DA93696E | CCAAT-binding transcription factor B (NF-Y family) | x | | |
| Traes_6AL_C0E74F959 | CCAAT-binding transcription factor B (NF-Y family) | x | | |
| Traes_6BL_953171851 | CCAAT-binding transcription factor B (NF-Y family) | x | | |
| Traes_5AL_3270A6690 | Enhanced ethylene response protein 5 | | | x |
| Traes_5BL_7F84602F3 | Ethylene responsive transcription factor 5a | | | x |
| Traes_7BS_B1A8B5B8A | Ethylene-responsive element binding protein 1 | | | x |
| Traes_2BL_94E5996F7 | FD-like 15 protein | | | x |
| Traes_7BL_EA8DE756D | Flavin-containing monooxygenase | | | x |
| Traes_6AS_2B85F4104 | Mediator of RNA polymerase II transcription subunit 18 | | | x |
| Traes_5AL_13E2DEC48 | MIKC-type MADS-box transcription factor WM6 | | | x |
| Traes_4AL_98EA87DFA1 | MYB-related protein (MYB family) | | | x |
| Traes_4BL_4D370C9FA | Probable mediator of RNA polymerase II transcription subunit 37c | | x | |
| Traes_4BL_4D370C9FA1 | Probable mediator of RNA polymerase II transcription subunit 37c | | x | |
| Traes_3AS_E18F065A3 | Protein ABCI7, chloroplastic | | | x |
| Traes_3AL_E5CB6F48A | Protein PLASTID TRANSCRIPTIONALLY ACTIVE 12 (PTAC family) | | | x |
| Traes_3AL_89EF640CE | Protein PLASTID TRANSCRIPTIONALLY ACTIVE 12 (PTAC family) | x | | |
| Traes_1AL_D8BA4FB5D | Protein PLASTID TRANSCRIPTIONALLY ACTIVE 14 (PTAC family) | x | | |
| Traes_1BL_E1AD16CAC | Protein PLASTID TRANSCRIPTIONALLY ACTIVE 14 (PTAC family) | x | | |
| Traes_3AL_C25CC8AB1 | Protein PLASTID TRANSCRIPTIONALLY ACTIVE 7 (PTAC family) | x | | |
| Traes_2AL_5BA7E2623 | TaAP2-A | | | x |
| Traes_7AL_6279FCE8B | Transcription activator GLK2 | | | x |
| Traes_7AL_6279FCE8B | Transcription activator GLK2 | | | x |
| Traes_6BS_05264DAEA | Transcription factor HBP-1a | | | x |
| Traes_1BL_AF69F4610 | Transcription factor UNE10 | | x | |
| Traes_1AL_914726769 | Transcription factor UNE10 | | x | |
| Traes_7AS_3BAA88E9E | Transcription termination factor MTERF6, chloroplastic/mitochondrial | x | | |

| | | | | | |
|---|---|---|---|---|---|
| | Traes_3B_07317CB79 | Transcriptional corepressor LEUNIG | | x | |
| | Traes_4AL_26117D669 | RNA polymerase sigma factor | x | | |
| | Traes_4AS_26117D669 | RNA polymerase sigma factor | x | | |
| | Traes_4AS_DE644E0CF | RNA polymerase sigma factor | | | x |
| | Traes_7AS_3BAA88E9E | Transcription termination factor MTERF6, chloroplastic/mitochondrial | x | | |
| | Traes_2AS_94603BB74 | Translation initiation factor | x | | |

| **Kinases** | **Gene ID** | **Description** | **common** | **Molise Sel. Colli** | **Simeto** |
|---|---|---|---|---|---|
| | Traes_4AS_A795672BE | Adenylyl-sulfate kinase | x | | |
| | Traes_1AL_928DC3A3C | Bifunctional riboflavin kinase/FMN phosphatase (FHY) | x | | |
| | Traes_1AL_7C3E4A07E | Diacylglycerol kinase (DGK) | | | x |
| | Traes_5AS_C28641518 | Diacylglycerol kinase (DGK) | | | x |
| | Traes_3AL_3BC2FF0ED | Fructokinase-like 1, chloroplastic | | | x |
| | Traes_2BS_959A4E58A | Fructokinase-like 2, chloroplastic | x | | |
| | Traes_4AL_8C4927CF5 | Lectin receptor kinase | | x | |
| | Traes_6AS_691D1AC82 | Non-specific serine/threonine protein kinase | | x | |
| | Traes_1AL_5F0F4FAFB | Non-specific serine/threonine protein kinase | | x | |
| | Traes_4AS_538F2E3BD | Non-specific serine/threonine protein kinase | | | x |
| | Traes_1BS_C385DE03B | Non-specific serine/threonine protein kinase | x | | |
| | Traes_2BS_0ABED462E | Non-specific serine/threonine protein kinase | x | | |
| | Traes_3AL_66EAB4B4F | Non-specific serine/threonine protein kinase | x | | |
| | Traes_5BL_12214988F | Non-specific serine/threonine protein kinase | x | | |
| | Traes_2AS_6345FCE27 | Non-specific serine/threonine protein kinase | | | x |
| | Traes_1AL_1538AC680 | Nucleoside diphosphate kinase (NDK) | | x | |
| | Traes_1BL_ACC9DDA86 | Nucleoside diphosphate kinase (NDK) | | x | |
| | Traes_5AS_8A3C47D2F | Nucleoside diphosphate kinase (NDK) | x | | |
| | Traes_1AL_6F2E87864 | Nucleoside diphosphate kinase 1 | x | | |
| | Traes_7BS_BEA71AF3F | Receptor-like serine/threonine protein kinase 2 | x | | |
| | Traes_6BS_C44E33CC9 | Receptor-like serine/threonine-protein kinase NCRK | | | x |
| | Traes_3AL_DBED56B67 | Serine/threonine-protein kinase | x | | |
| | Traes_5BL_DFFA40472 | Serine/threonine-protein kinase | x | | |

| | | | | | |
|---|---|---|---|---|---|
| | Traes_2BS_EA063BB91 | Serine/threonine-protein kinase | | x | |
| | Traes_2AL_BDC8DD6C7 | Serine/threonine-protein kinase | | | x |
| | Traes_2BL_07796B6EA | Serine/threonine-protein kinase | | | x |
| | Traes_2BL_6FB5FEFC61 | Serine/threonine-protein kinase | | | x |
| | Traes_2BL_9B20E7E7D | Serine/threonine-protein kinase | | | x |
| | Traes_6AL_7D22244E7 | Serine/threonine-protein kinase | | | x |
| | Traes_2BL_09098E11D | Serine/threonine-protein kinase | | | x |
| | Traes_2BL_E1C37619B | Serine/threonine-protein kinase | | | x |
| | Traes_5AL_BC7DEEA21 | Serine/threonine-protein kinase TOUSLED | | | x |
| | Traes_2BL_D195BD715 | Serine/threonine-protein kinase | | | x |
| | Traes_7AS_7F2987981 | Uridine kinase | x | | |
| | Traes_7BS_C57477AA31 | Uridine kinase | | | x |

| Stress | Gene ID | Description | common | Molise Sel. Colli | Simeto |
|---|---|---|---|---|---|
| | Traes_6BL_4DB266957 | 2-C-methyl-D-erythritol 2,4-cyclodiphosphate synthase, chloroplastic | | x | |
| | Traes_7AL_772A49098 | 2-succinylbenzoate--CoA ligase, chloroplastic/peroxisomal | x | | |
| | Traes_7AS_C73647B58 | 3-hydroxy-3-methylglutaryl coenzyme A reductase | x | | |
| | Traes_2BL_657D93794 | 6,7-dimethyl-8-ribityllumazine synthase (riboflavin synthase) | x | | |
| | Traes_1AS_BEFEF1613 | Acetylornithine aminotransferase, chloroplastic/mitochondrial | | | x |
| | Traes_2AL_CD28AB70E | Amine oxidase | | | x |
| | Traes_2AL_F4F6F518D | Amine oxidase | | | x |
| | Traes_2BS_65836FEE4 | Amine oxidase | x | | |
| | Traes_2AS_E0AFA6D18 | ATP sulfurylase 2 | | | x |
| | Traes_2BS_F70FE410B | ATP sulfurylase 2 | | | x |
| | Traes_1AL_52E56C8D3 | ATP synthase subunit beta | | x | |
| | Traes_3B_00CF2A894 | ATP-dependent Clp protease proteolytic subunit 3, chloroplastic | | | x |
| | Traes_4AS_0B56CEBD5 | ATP-dependent Clp protease proteolytic subunit-related protein 3, chloroplastic | x | | |
| | Traes_2AL_1E96BBDD4 | Autophagy-related protein 101 | | | x |
| | Traes_3AL_27DCCC109 | Autophagy-related protein 7c | | | x |

| | | | | |
|---|---|---|---|---|
| Traes_3AL_81D411955 | Autophagy-related protein 7c | | | x |
| Traes_5BL_763F3FEA8 | Autophagy-related protein 8i | | | x |
| Traes_7BL_6097E6ADE | Beta-carotene isomerase D27, chloroplastic | | | x |
| Traes_5BL_F0FC14B2A | Bifunctional monothiol glutaredoxin-S16, chloroplastic | | | x |
| Traes_7BS_8B18BE79B | Caleosin | x | | |
| Traes_6AS_FC9E9843D | Caleosin | x | | |
| Traes_4BL_53CBCD014 | Calmodulin TaCaM1-1 | x | | |
| Traes_2BS_D4CB3FB08 | Carboxypeptidase | | | x |
| Traes_3AS_B605964C1 | Carboxypeptidase | | | x |
| Traes_1AL_16EAFFACD | Carboxypeptidase | | | x |
| Traes_4BS_AD3C7F651 | Carboxypeptidase | | x | |
| Traes_1BS_FEA181281 | Carboxypeptidase | | x | |
| Traes_3AL_A306585EF | Carboxypeptidase | x | | |
| Traes_1AL_B55CC2568 | Carotene epsilon-monooxygenase, chloroplastic | | | x |
| Traes_1BL_714F4E4AC | Carotene epsilon-monooxygenase, chloroplastic | | | x |
| Traes_4BL_664A41517 | Catalase | | | x |
| Traes_5AL_34C62DB80 | CDGSH iron-sulfur domain-containing protein NEET | | | x |
| Traes_5BL_E86097AA2 | Chalcone-flavonone isomerase family protein | | | x |
| Traes_3AL_883A4805C | Cytokinin riboside 5'-monophosphate phosphoribohydrolase | | x | |
| Traes_7BS_5781FD481 | Dihydroxy-acid dehydratase, chloroplastic | | | x |
| Traes_5AL_288D3A2F4 | Diphosphomevalonate decarboxylase | | | x |
| Traes_5BL_AC016C8E2 | Diphosphomevalonate decarboxylase | | x | |
| Traes_7AL_F516C7DFF | Dolichyl-diphosphooligosaccharide--protein glycosyltransferase subunit DAD1 | | | x |
| Traes_2AS_7D3EC9FC2 | E3 ubiquitin-protein ligase | | x | |
| Traes_7AS_C38093E5C | ECPT-type aminoalcoholphosphotransferase | | | x |
| Traes_4AS_94B8E6F8C | FAD-linked sulfhydryl oxidase ERV1 | | | x |
| Traes_7AS_34AF6845D1 | F-box protein MAX2 | | x | |
| Traes_7BS_3285CDC24 | F-box protein MAX2 | x | | |
| Traes_4AL_497775E8F | ferritin | | | x |
| Traes_4BS_FFF12BA9E | ferritin | | | x |
| Traes_4AL_498512281 | ferritin | x | | |

| | | | | |
|---|---|---|---|---|
| Traes_4BS_BA655294C | ferritin | x | | |
| Traes_7BL_EA8DE756D | Flavin-containing monooxygenase | | | x |
| Traes_7AS_F91D8701E | GA-induced protein | | | x |
| Traes_6AS_4C413AC53 | Gamma-glutamylcyclotransferase | | x | |
| Traes_6BS_A679AA5FF | Gamma-glutamylcyclotransferase | | x | |
| Traes_1AL_F18EC96DF | GBF1 | | | x |
| Traes_2AL_E06A248EE | Glutathione peroxidase | | | x |
| Traes_4AS_519B67360 | Glutathione reductase, chloroplastic | x | | |
| Traes_4BL_9BA5AB82C | Glutathione reductase, chloroplastic | x | | |
| Traes_7AS_0BB13DD79 | Glutathione S-transferase DHAR3, chloroplastic | | | x |
| Traes_6AS_D537A116B | Glutathione synthetase | | x | |
| Traes_6BS_AA7101C2C | Glutathione synthetase | | | x |
| Traes_3AL_1D9B2CE84 | Glyoxylate/succinic semialdehyde reductase 2, chloroplastic | | | x |
| Traes_2BL_088F3C674 | Guard cell S-type anion channel SLAC1 | | | x |
| Traes_5BL_37ECD3B1E | Heat shock protein 90-5, chloroplastic | | | x |
| Traes_5AL_1DA3B4631 | Heat shock protein 90-5, chloroplastic | | x | |
| Traes_4AS_53B185E1A | HVA22-like protein | x | | |
| Traes_2BL_7BB7F9970 | HVA22-like protein | | | x |
| Traes_1BL_7F9006C51 | Hyperosmolality-gated Ca2+ permeable channel | | x | |
| Traes_5AL_07C125B1C | Hypersensitive induced reaction protein 2 | | | x |
| Traes_5AL_C0A407A93 | Hypersensitive induced response protein 3 | | | x |
| Traes_3AL_E5520535A | Laccase | | | x |
| Traes_2AS_B2C72994E | Lactoylglutathione lyase | | | x |
| Traes_4AL_A5AC2BDF2 | Lipoxygenase | | | x |
| Traes_2AL_5BAB26827 | Lipoxygenase | | x | |
| Traes_5BS_060785740 | Lipoxygenase | | x | |
| Traes_2BL_77148B8D8 | Lipoxygenase | x | | |
| Traes_5BL_304FAFA26 | Lipoxygenase | | | x |
| Traes_4BL_BD1E1BEFA | MLO-like protein | | | x |
| Traes_2AS_1FC67C1BD | MLO-like protein | | x | |
| Traes_5AL_268DF2E10 | MLO-like protein | | x | |
| Traes_5AL_E7A3EDC5F | MLO-like protein | | x | |

| | | | | |
|---|---|---|---|---|
| Traes_2AS_9F72E517F | NADPH-dependent thioredoxin reductase 3 | | | x |
| Traes_6AS_101337AE9 | NO-associated protein 1, chloroplastic/mitochondrial | | x | |
| Traes_4BL_96C906D61 | Non-specific lipid-transfer protein | | x | |
| Traes_3AS_5A72CA3A7 | Non-specific lipid-transfer protein | | | x |
| Traes_7BL_2DBEEEBC2 | Nudix hydrolase 10 | x | | |
| Traes_5AL_43F9E9C1D | Nudix hydrolase 23, chloroplastic | | | x |
| Traes_5BL_F0D3D6D2E | Nudix hydrolase 23, chloroplastic | x | | |
| Traes_5AL_64517027B | Obg-like ATPase 1 | x | | |
| Traes_5BL_907D7067C | Obg-like ATPase 1 | x | | |
| Traes_5AL_4A88E0A3A | Pectin acetylesterase | | | x |
| Traes_3AL_11D85AEF9 | Pectin acetylesterase | | | x |
| Traes_5AL_955D2899A | Pectin acetylesterase | | | x |
| Traes_6BL_C83912BDE | Pectin acetylesterase | | | x |
| Traes_2AL_6A0320615 | Pectin acetylesterase | | x | |
| Traes_1BL_46FB3D95E | Peptide chain release factor PrfB3, chloroplastic | | | x |
| Traes_7BL_8930CD1E5 | Peptide methionine sulfoxide reductase B1, chloroplastic | | x | |
| Traes_1AS_C70D49E2E | Peroxidase | | | x |
| Traes_2AL_20BC426B9 | Peroxidase | | | x |
| Traes_2AL_2224DCC1A | Peroxidase | | | x |
| Traes_2AL_520618712 | Peroxidase | | | x |
| Traes_7AL_3E96F374E | Peroxidase | | | x |
| Traes_7AS_020CCE3DB | Peroxiredoxin Q, chloroplastic | | | x |
| Traes_5AL_2F97350D2 | Phospholipase D delta | x | | |
| Traes_5BL_A0ED5623C | Phospholipase D delta | | x | |
| Traes_4AS_1D24D36C4 | Photosystem II repair protein PSB27-H1, chloroplastic | | | x |
| Traes_7AL_E9733B552 | Poly [ADP-ribose] polymerase (PARP) | | | x |
| Traes_6AS_0F0809F96 | Probable inactive linolenate hydroperoxide lyase | | | x |
| Traes_1AS_4C784AA40 | Protease Do-like 2, chloroplastic | x | | |
| Traes_6BS_4A0C34E63 | Protein ACCUMULATION AND REPLICATION OF CHLOROPLASTS 6, chloroplastic | x | | |
| Traes_1BS_8C0F2821A | Protein ALUMINUM SENSITIVE 3 | | x | |
| Traes_5AL_5AF616715 | Protein C2-DOMAIN ABA-RELATED 11 | | x | |

| | | | | |
|---|---|---|---|---|
| Traes_2BS_7093A1D1A | Protein DETOXIFICATION | | | x |
| Traes_5BL_5A648D29D | Protein DETOXIFICATION | | x | |
| Traes_7AS_85178E0D7 | Protein DETOXIFICATION | | x | |
| Traes_3B_189FA2C94 | Protein DETOXIFICATION | | | x |
| Traes_7AS_E477E850F | Protein DETOXIFICATION | | | x |
| Traes_7BS_6A8DBF06C | Protein DETOXIFICATION | | | x |
| Traes_5AL_9697E163B | Protein DETOXIFICATION | x | | |
| Traes_4AL_5EC1ABB77 | Protein DETOXIFICATION | x | | |
| Traes_7AL_970814850 | Protein DETOXIFICATION | x | | |
| Traes_1AL_A970EBB27 | Protein DETOXIFICATION 42 | | | x |
| Traes_1BL_DCC58CE85 | Protein DETOXIFICATION 42 | | x | |
| Traes_4AS_28DA71FF6 | Protein DETOXIFICATION 43 | | | x |
| Traes_4BL_06F78B23C | Protein DETOXIFICATION 43 | | | x |
| Traes_6AL_763EE2AED | Protein DETOXIFICATION 45, chloroplastic | | | x |
| Traes_1AL_4B7636BC9 | Protein LOL1 | | x | |
| Traes_7BL_E2E7AA383 | Protein OVEREXPRESSOR OF CATIONIC PEROXIDASE 3 | | x | |
| Traes_1AL_FE094DCAF | Protein PALE CRESS, chloroplastic | | | x |
| Traes_1BL_74FC9D6D1 | Protein PALE CRESS, chloroplastic | | | x |
| Traes_1AL_AF222553D | Rac-like GTP-binding protein ARAC3 | | | x |
| Traes_6AL_6CD6A9215 | Rhodanese-like domain-containing protein 11, chloroplastic | | | x |
| Traes_6BL_F4AF4CD45 | Rhodanese-like domain-containing protein 11, chloroplastic | | | x |
| Traes_6AS_FD8F6B539 | Rhodanese-like domain-containing protein 7 | x | | |
| Traes_6AL_E70725BF0 | Riboflavin biosynthesis protein PYRR, chloroplastic | | | x |
| Traes_6AL_D13D9D951 | Serine carboxypeptidase-like 34 | | | x |
| Traes_4AS_060E22DA0 | Serine/arginine-rich splicing factor SR45a | | x | |
| Traes_4BL_7BFE58ED9 | Serine/arginine-rich splicing factor SR45a | | x | |
| Traes_4BL_F6A0E1CCB | Signal recognition particle 43 kDa protein, chloroplastic | | | x |
| Traes_6AL_5537AC81D | Soluble inorganic pyrophosphatase 6, chloroplastic | x | | |
| Traes_6BL_92B298585 | Soluble inorganic pyrophosphatase 6, chloroplastic | x | | |
| Traes_2BL_CCD296233 | Stress enhanced protein 2, chloroplastic | x | | |
| Traes_2AS_0C3C91D16 | Stress responsive protein | | x | |

| | | | | | |
|---|---|---|---|---|---|
| | Traes_2BS_762E60A0B | Stress responsive protein 1 | | x | |
| | Traes_6BS_1C714CE3E | Succinate-semialdehyde dehydrogenase, mitochondrial | | | x |
| | Traes_2AL_D0D84176E | Superoxide dismutase | x | | |
| | Traes_7BL_70F07C889 | Superoxide dismutase [Cu-Zn] 2, chloroplastic | | | x |
| | Traes_7BL_A42D6C984 | Superoxide dismutase [Cu-Zn] 2, chloroplastic | | | x |
| | Traes_4AL_47DDDE0C1 | Thioredoxin-like protein CITRX, chloroplastic | | | x |
| | Traes_4BS_D9A4219F2 | Thioredoxin-like protein CITRX, chloroplastic | | x | |
| | Traes_6BL_462C2179A | Tripeptidyl-peptidase 2 | | x | |
| | Traes_6BL_4DB1AC6CC | Tripeptidyl-peptidase 2 | | x | |
| | Traes_1AL_0C964B45F | Tubulin alpha chain | | x | |
| | Traes_2BS_AA6D8150A | Tubulin alpha chain | | x | |
| | Traes_2AS_34272605C | Tubulin alpha chain | | x | |
| | Traes_4BS_0BDB0D05C | Tubulin beta chain | | | x |
| | Traes_1BS_2F6DD776E | Tubulin-folding cofactor E | | | x |
| | Traes_6AL_4048F89A6 | Ubiquitin carboxyl-terminal hydrolase | | | x |
| | Traes_3AL_6AF4297B2 | Vacuolar proton ATPase subunit E | | | x |
| | Traes_2BL_5E868851D | Violaxanthin de-epoxidase, chloroplastic | | | x |
| | Traes_2AL_FA3E7C60B | Violaxanthin de-epoxidase, chloroplastic | | | x |
| | Traes_5BS_2983EE249 | V-type proton ATPase proteolipid subunit | | | x |
| | Traes_5BL_2983EE249 | V-type proton ATPase proteolipid subunit | | | x |
| | Traes_3AL_182FFAF75 | Ycf3-interacting protein 1, chloroplastic | | | x |

| **Others** | **Gene ID** | **Description** | **common** | **Molise Sel. Colli** | **Simeto** |
|---|---|---|---|---|---|
| | Traes_5AL_A8C598914 | 20 kDa chaperonin, chloroplastic | x | | |
| | Traes_5BL_5041B968E | 20 kDa chaperonin, chloroplastic | x | | |
| | Traes_4AS_C420DEB6D | 30S ribosomal protein S1, chloroplastic | | | x |
| | Traes_4BL_C3C6C11C4 | 30S ribosomal protein S1, chloroplastic | | | x |
| | Traes_4AS_6875248F8 | 30S ribosomal protein S10, chloroplastic | x | | |
| | Traes_4BL_EFFA96D15 | 30S ribosomal protein S10, chloroplastic | x | | |
| | Traes_4BS_0ECAB85C2 | 30S ribosomal protein S13, chloroplastic | x | | |
| | Traes_7AL_A1ECABC74 | 30S ribosomal protein S31, chloroplastic | x | | |
| | Traes_7BL_E67311208 | 30S ribosomal protein S31, chloroplastic | | | x |

| | | | | |
|---|---|---|---|---|
| Traes_4AL_FB6127E85 | 30S ribosomal protein S6 alpha, chloroplastic | x | | |
| Traes_5BL_BB5612CA4 | 30S ribosomal protein S6 alpha, chloroplastic | x | | |
| Traes_5AL_55D7BFD57 | 30S ribosomal protein S9, chloroplastic | | | x |
| Traes_2AS_105B1A29A | 40S ribosomal protein S12 | x | | |
| Traes_2BS_7700613D4 | 40S ribosomal protein S12 | x | | |
| Traes_4BS_7700613D4 | 40S ribosomal protein S12 | x | | |
| Traes_5BL_8601EC2E0 | 40S ribosomal protein S15a | | x | |
| Traes_6BL_EFAF71592 | 40S ribosomal protein S24 | | x | |
| Traes_7AL_E7E665596 | 40S ribosomal protein S24 | | x | |
| Traes_7BL_0F1408688 | 40S ribosomal protein S24 | x | | |
| Traes_5AL_106BEC5D4 | 40S ribosomal protein S26 | | | x |
| Traes_5BL_EC6277E0E | 40S ribosomal protein S26 | | | x |
| Traes_4AS_8ED92AF77 | 40S ribosomal protein S3a | | | x |
| Traes_4BL_7D2DDCA26 | 40S ribosomal protein S3a | | | x |
| Traes_4BL_66E76E6B0 | 40S ribosomal protein S3a | x | | |
| Traes_1AS_E58B0B23A | 40S ribosomal protein S4 | | x | |
| Traes_1BS_DD36A7A9B | 40S ribosomal protein S4 | | x | |
| Traes_2AS_9CFA03D22 | 40S ribosomal protein S6 | | | x |
| Traes_2AS_E08BF60FE | 40S ribosomal protein S6 | | | x |
| Traes_2BS_C41E04839 | 40S ribosomal protein S6 | | | x |
| Traes_2AS_C7813CD47 | 40S ribosomal protein S6 | | | x |
| Traes_2AL_1B2CC9926 | 40S ribosomal protein S8 | x | | |
| Traes_2BL_7BC3942D6 | 40S ribosomal protein S8 | x | | |
| Traes_2AS_3F458D2CF | 40S ribosomal protein SA | x | | |
| Traes_2BS_ECC74F149 | 40S ribosomal protein SA | x | | |
| Traes_3AS_7F20E18E2 | 50S ribosomal protein 5, chloroplastic | x | | |
| Traes_1BL_8AC8D124C | 50S ribosomal protein L1, chloroplastic | | | x |
| Traes_1AL_3B838C090 | 50S ribosomal protein L1, chloroplastic | x | | |
| Traes_1AL_B01B26670 | 50S ribosomal protein L1, chloroplastic | x | | |
| Traes_5BL_AA5911949 | 50S ribosomal protein L17, chloroplastic | x | | |
| Traes_5BL_AA5911949 | 50S ribosomal protein L17, chloroplastic | x | | |
| Traes_6AS_EB8BA8FE2 | 50S ribosomal protein L21, chloroplastic | x | | |

| | | | | |
|---|---|---|---|---|
| Traes_7AL_EA6B88D3F | 50S ribosomal protein L24, chloroplastic | x | | |
| Traes_7BL_C99BF5CA9 | 50S ribosomal protein L24, chloroplastic | x | | |
| Traes_3AL_C134C28BC | 50S ribosomal protein L27, chloroplastic | | | x |
| Traes_1BS_71C6FDF1A | 50S ribosomal protein L28, chloroplastic | x | | |
| Traes_6BL_A66A6DCB2 | 50S ribosomal protein L29, chloroplastic | | | x |
| Traes_6AL_33DE178C5 | 50S ribosomal protein L29, chloroplastic | | | x |
| Traes_6AS_478618F75 | 50S ribosomal protein L3-1, chloroplastic | x | | |
| Traes_6BS_AEB9C88B3 | 50S ribosomal protein L3-1, chloroplastic | x | | |
| Traes_3AS_DF92A1FC5 | 50S ribosomal protein L3-2, chloroplastic | | x | |
| Traes_3AL_4EC74A033 | 50S ribosomal protein L34, chloroplastic | x | | |
| Traes_7BL_7A7BC0D15 | 50S ribosomal protein L35 | x | | |
| Traes_4BS_6EAD366CD | 50S ribosomal protein L6, chloroplastic | x | | |
| Traes_4BS_6EAD366CD | 50S ribosomal protein L6, chloroplastic | x | | |
| Traes_4AS_3CD9834D6 | 50S ribosomal protein L6, chloroplastic | | | x |
| Traes_4AS_49782B510 | 50S ribosomal protein L6, chloroplastic | | | x |
| Traes_5AL_4E0638B3E | 50S ribosomal protein L6, chloroplastic | | | x |
| Traes_7BL_86BB13ED9 | 60S ribosomal protein L13 | x | | |
| Traes_7AL_4A3378136 | 60S ribosomal protein L13 | x | | |
| Traes_3AL_7BCADDE08 | 60S ribosomal protein L18a | | x | |
| Traes_5AL_0C11FAA4D | 60S ribosomal protein L21 | x | | |
| Traes_3AL_350DFFE84 | 60S ribosomal protein L36 | x | | |
| Traes_5BS_26A43A8A0 | 60S ribosomal protein L36 | x | | |
| Traes_5AS_6D0CC3B1F | 60S ribosomal protein L36 | | | x |
| Traes_1BL_0FC423766 | 60S ribosomal protein L36 | | | x |
| Traes_3AL_8131EBAD8 | 60S ribosomal protein L37a, expressed | | x | |
| Traes_6BL_168034B0D | 60S ribosomal protein L6 | x | | |
| Traes_6AL_88DE049AB | ACT domain-containing protein ACR3 | | x | |
| Traes_4BL_5AF52F028 | Actin-depolymerizing factor 5 | | | x |
| Traes_4AS_811353465 | Actin-depolymerizing factor 5 | x | | |
| Traes_7AS_545BDB6F1 | Alanine--tRNA ligase, chloroplastic/mitochondrial | | | x |
| Traes_5BL_E123D8905 | ALBINO3-like protein 1, chloroplastic | | | x |
| Traes_6AL_CFE60316E | Annexin | x | | |

| | | | | |
|---|---|---|---|---|
| Traes_6AL_23B31796F | Aquaporin 7 | | | x |
| Traes_6BL_0C17AE0D2 | Armadillo repeat-containing protein LFR | | | x |
| Traes_2AS_0F4EA930D | Asparagine--tRNA ligase, chloroplastic/mitochondrial | x | | |
| Traes_2AS_7D65E81D0 | Asparagine--tRNA ligase, chloroplastic/mitochondrial | | | x |
| Traes_2BS_FF8887BE5 | Asparagine--tRNA ligase, chloroplastic/mitochondrial | | | x |
| Traes_1AL_975CB04EA | ATP-dependent Clp protease proteolytic subunit | | | x |
| Traes_4AL_1F49A9D0D | ATP-dependent Clp protease proteolytic subunit | x | | |
| Traes_7AS_090FDB9B1 | ATP-dependent Clp protease proteolytic subunit | x | | |
| Traes_4AS_C67AE6944 | ATP-dependent Clp protease proteolytic subunit | x | | |
| Traes_4BL_1D9E2AC36 | ATP-dependent Clp protease proteolytic subunit | x | | |
| Traes_1AL_3D9EA5952 | ATP-dependent Clp protease proteolytic subunit-related protein 1, chloroplastic | | | x |
| Traes_7AL_356F6FF0E | Blue-light photoreceptor PHR2 | | | x |
| Traes_3AL_12DCD0158 | Cell division protein FtsY homolog, chloroplastic | | | x |
| Traes_2BL_6097FE67E | Cell division protein FtsZ homolog 1, chloroplastic | x | | |
| Traes_2AL_086336B60 | Chaperonin 60 subunit beta 4, chloroplastic | x | | |
| Traes_2BL_5A94F1657 | Chaperonin 60 subunit beta 4, chloroplastic | x | | |
| Traes_2AL_64E61AB1B | CRM-domain containing factor CFM2, chloroplastic | | | x |
| Traes_2AL_49BBA1645 | CRM-domain containing factor CFM2, chloroplastic | x | | |
| Traes_2BL_8CCB9C88D | CRM-domain containing factor CFM2, chloroplastic | x | | |
| Traes_6AL_86BABC721 | Cryptochrome-2 | | x | |
| Traes_6AL_86BABC721 | Cryptochrome-2 | | x | |
| Traes_5AL_EF0D9834F | Cysteine--tRNA ligase, chloroplastic/mitochondrial | x | | |
| Traes_5BL_41F9FAA36 | Cysteine--tRNA ligase, chloroplastic/mitochondrial | x | | |
| Traes_5BL_0684B3A02 | Cytochrome c biogenesis protein CCS1, chloroplastic | x | | |
| Traes_2BS_642DC63EC | DEAD-box ATP-dependent RNA helicase 10 | | | x |
| Traes_5AL_D952BA6DB | DEAD-box ATP-dependent RNA helicase 22 | | | x |
| Traes_4AL_199EE1839 | DEAD-box ATP-dependent RNA helicase 3 | | | x |
| Traes_5BL_BD895CF84 | DEAD-box ATP-dependent RNA helicase 3 | | | x |
| Traes_4AL_199EE1839 | DEAD-box ATP-dependent RNA helicase 3, chloroplastic | x | | |
| Traes_5BL_BD895CF84 | DEAD-box ATP-dependent RNA helicase 3, chloroplastic | x | | |
| Traes_3AS_D097872BE | DEAD-box ATP-dependent RNA helicase 39 | x | | |

| | | | | |
|---|---|---|---|---|
| Traes_3AS_F527FB55B | DEAD-box ATP-dependent RNA helicase 39 | x | | |
| Traes_6AL_D153AC732 | DEAD-box ATP-dependent RNA helicase 47, mitochondrial | x | | |
| Traes_6BL_3C4549AB1 | DEAD-box ATP-dependent RNA helicase 47, mitochondrial | x | | |
| Traes_5AL_B72E1DD70 | DEAD-box ATP-dependent RNA helicase 50 | x | | |
| Traes_3B_F4870ACB3 | DEAD-box ATP-dependent RNA helicase 50 | x | | |
| Traes_3B_420994D45 | DEAD-box ATP-dependent RNA helicase 58, chloroplastic | x | | |
| Traes_3AS_30D175BEB | DNA gyrase subunit B | | x | |
| Traes_2AL_61B7762C7 | DNA ligase 4 | | | x |
| Traes_5BL_A60442138 | DNA mismatch repair protein MSH6 | | | x |
| Traes_3AL_E77A5F03A | DNA-directed RNA polymerase subunit | | | x |
| Traes_1BS_9F8A8021B | DNA-directed RNA polymerase subunit | | | x |
| Traes_6BS_54B2B1260 | DNA-directed RNA polymerase subunit | | | x |
| Traes_6BS_C85FE9482 | DNA-directed RNA polymerase V subunit 1 | | | x |
| Traes_1AS_AEEC8D740 | E3 ubiquitin-protein ligase UPL6 | | | x |
| Traes_2AS_7D3EC9FC2 | E3 ubiquitin-protein ligase | | x | |
| Traes_5AL_87C8496BE | Elongation factor 1-alpha | x | | |
| Traes_2AL_837BCD21C | Elongation factor G, chloroplastic | | | x |
| Traes_5AS_91BEB6A34 | Elongation factor Ts, mitochondrial | x | | |
| Traes_6AL_32FADD06A | Elongation factor Tu | x | | |
| Traes_6BL_A1759ED17 | Elongation factor Tu | x | | |
| Traes_6AL_54260E477 | Epimerase family protein SDR39U1 homolog, chloroplastic | | | x |
| Traes_2AL_95D918793 | Eukaryotic translation initiation factor 3 subunit H | x | | |
| Traes_2BS_4A4C9BD80 | Eukaryotic translation initiation factor 3 subunit M | | x | |
| Traes_5AS_2C7433827 | Eukaryotic translation initiation factor 5A | x | | |
| Traes_5BS_160C44A80 | Eukaryotic translation initiation factor 5A | x | | |
| Traes_5BS_8FB0A0D95 | Eukaryotic translation initiation factor 5A | x | | |
| Traes_2AL_2DCC96CD9 | Eukaryotic translation initiation factor 5A | | | x |
| Traes_4AL_E4470B79C | Flap endonuclease 1 | | | x |
| Traes_5BL_213C8B59D | Formin-like protein | | x | |
| Traes_3AS_CD8EF3E96 | FRIGIDA-like protein | | x | |
| Traes_7AL_492C9C1E4 | Glutamate receptor (GLR) | x | | |
| Traes_1BL_EC3E899FE | Glutamyl-tRNA reductase | | | x |

| | | | | |
|---|---|---|---|---|
| Traes_2AL_4D97AE321 | Glutamyl-tRNA(Gln) amidotransferase subunit A, chloroplastic/mitochondrial | x | | |
| Traes_2BL_0118B58B3 | Glutamyl-tRNA(Gln) amidotransferase subunit A, chloroplastic/mitochondrial | | | x |
| Traes_2BL_EEE4AF800 | Glutamyl-tRNA(Gln) amidotransferase subunit A, chloroplastic/mitochondrial | | | x |
| Traes_2AL_783420E8B | Glutamyl-tRNA(Gln) amidotransferase subunit B, chloroplastic/mitochondrial | x | | |
| Traes_6BS_C73BCAF43 | Glutamyl-tRNA(Gln) amidotransferase subunit B, chloroplastic/mitochondrial | x | | |
| Traes_4BL_9BA5AB82C | Glutamyl-tRNA(Gln) amidotransferase subunit B, chloroplastic/mitochondrial | x | | |
| Traes_5AL_F55E09494 | Glutamyl-tRNA(Gln) amidotransferase subunit C, chloroplastic/mitochondrial | | | x |
| Traes_6AL_96665E11C | GrpE protein homolog | x | | |
| Traes_6BL_2CD26F5E7 | GrpE protein homolog | x | | |
| Traes_2BL_F3EE344D7 | GrpE protein homolog | x | | |
| Traes_2AL_18913037F | GrpE protein homolog | | | x |
| Traes_7BL_91C7CD5C5 | GTP-binding nuclear protein | | x | |
| Traes_4AS_F1F72C66C | GTP-binding protein At2g22870 | x | | |
| Traes_1BL_0A996D4A2 | Histone H2B | | | x |
| Traes_2BL_BB2C0F695 | Histone H3 | | | x |
| Traes_5BL_71F88DB88 | Histone H4 | | | x |
| Traes_7BS_955956707 | Histone-lysine N-methyltransferase CLF | | | x |
| Traes_2BL_392793376 | Katanin p80 WD40 repeat-containing subunit B1 homolog | | x | |
| Traes_2AL_A3F599522 | Kinesin-like protein | | | x |
| Traes_3AS_BCC969DFF | Leucine--tRNA ligase, chloroplastic/mitochondrial | | | x |
| Traes_5AS_FC3C5CA46 | LIM domain-containing protein WLIM1 | | x | |
| Traes_4AL_603B6DC64 | LR34 | | | x |
| Traes_6AL_0CD92304D | Lysine--tRNA ligase | x | | |
| Traes_6BL_2936220F5 | Lysine--tRNA ligase | x | | |
| Traes_5BL_1DE088A30 | Lysine--tRNA ligase | x | | |
| Traes_5BL_CFC7BB483 | Lysine--tRNA ligase | x | | |
| Traes_6AS_2B85F4104 | Mediator of RNA polymerase II transcription subunit 18 | | | x |
| Traes_4AS_372712125 | Methionine--tRNA ligase, chloroplastic/mitochondrial | x | | |

| | | | | |
|---|---|---|---|---|
| Traes_4BL_B19528CE1 | Methionine--tRNA ligase, chloroplastic/mitochondrial | x | | |
| Traes_4AS_6CE64CB5B | Microtubule-associated protein 70-5 | | x | |
| Traes_4BL_1C570A289 | Microtubule-associated protein 70-5 | | | x |
| Traes_7AL_D7CCFE55C | Multiple organellar RNA editing factor 9, chloroplastic | x | | |
| Traes_7BL_FD98DF9E5 | Multiple organellar RNA editing factor 9, chloroplastic | x | | |
| Traes_1BL_BAA0AE22B | Nascent polypeptide-associated complex subunit beta | x | | |
| Traes_4BL_70F08F293 | Nascent polypeptide-associated complex subunit beta | x | | |
| Traes_5AL_AB1566CF9 | Nascent polypeptide-associated complex subunit beta | x | | |
| Traes_1AL_61DFD90BF | Nascent polypeptide-associated complex subunit beta | | | x |
| Traes_7AL_48612DFF5 | Novel plant SNARE 11 | | x | |
| Traes_4AL_4A747B462 | Nuclear poly(A) polymerase 1 | | | x |
| Traes_5BL_A9BFB8F58 | Pectin acetylesterase | x | | |
| Traes_5AL_85C8D32D0 | Pectin acetylesterase | x | | |
| Traes_3AL_0A528281C | Pectin acetylesterase | x | | |
| Traes_1BL_698AC1335 | Pentatricopeptide repeat-containing protein At1g15510, chloroplastic | | | x |
| Traes_2BL_B4DCEEB99 | Pentatricopeptide repeat-containing protein At2g41720 | | | x |
| Traes_4BS_8D508569E | Pentatricopeptide repeat-containing protein At3g06430, chloroplastic | | | x |
| Traes_7AS_CA6519CDB | Pentatricopeptide repeat-containing protein At3g18110, chloroplastic | | x | |
| Traes_2AL_09781134C | Pentatricopeptide repeat-containing protein At3g22150, chloroplastic | | | x |
| Traes_2BL_EB51A1B4A | Pentatricopeptide repeat-containing protein At3g22150, chloroplastic | | | x |
| Traes_5AL_B84365C59 | Pentatricopeptide repeat-containing protein At3g22150, chloroplastic | | | x |
| Traes_4AL_17C4B47B3 | Pentatricopeptide repeat-containing protein At4g31850, chloroplastic | | x | |
| Traes_5AL_64A3B1812 | Pentatricopeptide repeat-containing protein At4g39620, chloroplastic | x | | |
| Traes_6BL_34B508140 | Pentatricopeptide repeat-containing protein At4g39620, chloroplastic | x | | |
| Traes_2AL_9B7FE4EC3 | Pentatricopeptide repeat-containing protein At5g50280, chloroplastic | | | x |
| Traes_2BL_18A640F3B | Pentatricopeptide repeat-containing protein At5g50280, chloroplastic | | | x |

| | | | | |
|---|---|---|---|---|
| Traes_4BL_097C26EB0 | Peptide chain release factor APG3, chloroplastic | | | x |
| Traes_1BS_F7B1A6268 | Peptidyl-prolyl cis-trans isomerase | x | | |
| Traes_3AS_12D74A531 | Peptidyl-prolyl cis-trans isomerase CYP20-3, chloroplastic | | | x |
| Traes_6BS_387A98C03 | Peptidyl-prolyl cis-trans isomerase FKBP18, chloroplastic | | | x |
| Traes_5AL_F8FD20EF8 | Polyadenylate-binding protein | x | | |
| Traes_2BS_23B5153F9 | Polyribonucleotide nucleotidyltransferase 1, chloroplastic | | | x |
| Traes_2BS_F66779538 | Polyribonucleotide nucleotidyltransferase 1, chloroplastic | x | | |
| Traes_7AL_569D0BB1B | Pre-mRNA cleavage factor Im 25 kDa subunit 1 | x | | |
| Traes_1BL_A127DF6A9 | Probable transmembrane GTPase FZO-like, chloroplastic | | | x |
| Traes_1AL_4BF6C25E1 | Probable transmembrane GTPase FZO-like, chloroplastic | | | x |
| Traes_1BL_241A3B9EF | Probable ubiquitin-conjugating enzyme E2 24 | | | x |
| Traes_5BL_196F297B6 | Probable ubiquitin-conjugating enzyme E2 24 | | | x |
| Traes_3AS_4C0261CD5 | Probable zinc metalloprotease EGY2, chloroplastic | | | x |
| Traes_3AS_922982272 | Probable zinc metalloprotease EGY2, chloroplastic | x | | |
| Traes_5BL_5F8443EFD | Protein ACCUMULATION AND REPLICATION OF CHLOROPLASTS 3 | | | x |
| Traes_1BS_CE144DF81 | Protein ACCUMULATION AND REPLICATION OF CHLOROPLASTS 6, chloroplastic | x | | |
| Traes_6AS_62C8C07B6 | Protein ACCUMULATION AND REPLICATION OF CHLOROPLASTS 6, chloroplastic | x | | |
| Traes_6AS_87906149C | Protein CHLORORESPIRATORY REDUCTION 6, chloroplastic | | | x |
| Traes_6BS_C7BF687C2 | Protein CHLORORESPIRATORY REDUCTION 6, chloroplastic | | | x |
| Traes_6BS_63FCF7A14 | Protein CHLORORESPIRATORY REDUCTION 6, chloroplastic | | x | |
| Traes_2BL_985C16D8A | Proteinaceous RNase P 1, chloroplastic/mitochondrial | | | x |
| Traes_2AL_D31914DE6 | Proteinaceous RNase P 1, chloroplastic/mitochondrial | | | x |
| Traes_2AL_D31914DE6 | Proteinaceous RNase P 1, chloroplastic/mitochondrial | x | | |
| Traes_4AL_345A5A89F | Putative 30S ribosomal protein S13 | | | x |
| Traes_5AL_EDE003529 | Putative ribosomal protein S18 | | x | |
| Traes_3AL_31E49AA0F | Putative translation initiation factor | | | x |
| Traes_7BL_8F226AC0C | Reticulon-like protein | x | | |
| Traes_2AL_366D94C0B | Reticulon-like protein | x | | |

| | | | | |
|---|---|---|---|---|
| Traes_7AS_CDF144498 | Rhomboid-like protein | | x | |
| Traes_7AS_235A8B4D8 | Ribonuclease E/G-like protein, chloroplastic | | x | |
| Traes_7AS_97BCB0125 | Ribonuclease E/G-like protein, chloroplastic | | x | |
| Traes_5AL_CFEF57F4A | Ribonuclease Z, chloroplastic | x | | |
| Traes_5BL_19748672B | Ribonuclease Z, chloroplastic | x | | |
| Traes_3AL_137E874E4 | Ribosomal protein | x | | |
| Traes_3AL_74778D9DF | Ribosomal protein | x | | |
| Traes_2AS_76163A005 | Ribosomal protein L15 | x | | |
| Traes_2BS_100054707 | Ribosomal protein L15 | | x | |
| Traes_1BL_C706FAA2B | Ribosomal protein L17 | | x | |
| Traes_6AL_288540CF3 | Ribosomal protein L17 | x | | |
| Traes_1AL_D20D648FD | Ribosomal protein L17 | | | x |
| Traes_6BL_30EFB4704 | Ribosomal protein L17 | | | x |
| Traes_2AS_C750EF91E | Ribosomal protein L19 | | x | |
| Traes_2BS_A9BF58434 | Ribosomal protein L19 | | x | |
| Traes_2BS_CB8DF8A02 | Ribosomal protein L19 | | x | |
| Traes_2AS_E1D0B2CE3 | Ribosomal protein L19 | | x | |
| Traes_2AS_467FD5266 | Ribosomal protein L19 | | | x |
| Traes_2BS_7135D68C5 | Ribosomal protein L19 | | | x |
| Traes_6AL_D5111DA34 | Ribosomal protein L37 | x | | |
| Traes_5AS_8785C0B49 | Ribosomal protein L3-A3 | x | | |
| Traes_7AL_A1EA63AD2 | Ribosomal protein P1 | | | x |
| Traes_4AL_001CD0D65 | Ribosomal protein S20 | | x | |
| Traes_2AL_16CF7A3BA | Ribosomal protein S20 | x | | |
| Traes_4AS_98267C6F6 | Ribosomal protein S7 | x | | |
| Traes_4BL_BF17A941B | Ribosomal protein S7 | x | | |
| Traes_2AL_EC2CE85AC | RING-type E3 ubiquitin transferase | | | x |
| Traes_2AL_F5A6D2E35 | RNA-binding protein CP33, chloroplastic | x | | |
| Traes_2BL_8CC7DE748 | RNA-binding protein CP33, chloroplastic | x | | |
| Traes_3AS_ECAACF24F | RNA-dependent RNA polymerase | x | | |
| Traes_3B_2C6DB84FB | RNA-dependent RNA polymerase | x | | |
| Traes_6AL_13BC97E04 | RNA-dependent RNA polymerase | | | x |

| | | | | | |
|---|---|---|---|---|---|
| | Traes_4AL_5D769AED8 | RuvB-like helicase | | | x |
| | Traes_5BS_5361C483D | Synonyms | x | | |
| | Traes_2BL_FE46FC938 | TaAP2-B | x | | |
| | Traes_7AS_378A12AA9 | TGW-7A | | x | |
| | Traes_5BL_369BF7271 | Thioredoxin-like protein HCF164, chloroplastic | | | x |
| | Traes_1BL_F1640B356 | Thylakoid lumenal 17.4 kDa protein, chloroplastic | | | x |
| | Traes_1AL_5558B7A4C | Thylakoid lumenal 17.4 kDa protein, chloroplastic | x | | |
| | Traes_1AL_5558B7A4C | Thylakoid lumenal 17.4 kDa protein, chloroplastic | x | | |
| | Traes_4AS_553C18F16 | Translocase of chloroplast | x | | |
| | Traes_4BL_E48C704F1 | Translocase of chloroplast | x | | |
| | Traes_5AL_A9C9D731A | Triticain gamma | | | x |
| | Traes_2BS_E5A5144E0 | tRNA (guanine(26)-N(2))-dimethyltransferase | | | x |
| | Traes_7AL_6ED296411 | tRNA dimethylallyltransferase 9 | | x | |
| | Traes_3AL_7494F4783 | Tryptophan--tRNA ligase, chloroplastic/mitochondrial | | | x |
| | Traes_1AL_FE1097B22 | Tyrosine--tRNA ligase, chloroplastic/mitochondrial | x | | |
| | Traes_6AS_E011BC5BB | Ubiquitin-conjugating enzyme E2 29 | | x | |
| | Traes_2AL_496A24411 | Ubiquitin-like-conjugating enzyme ATG10 | x | | |
| | Traes_4BL_5FCA04C3B | YlmG homolog protein 2, chloroplastic | | | x |

**Table S3.** Number of DEGs (up-/down-regulated) under two N conditions, enriched GO terms before and after FDR correction in emmer (Molise Sel. Colli), durum wheat (Simeto), and in total.

| Genotypes | up/ down | DE genes | GO enriched | Filtered GO (FDR) |
|---|---|---|---|---|
| emmer (Molise Sel. Colli) | up | 1202 | 486 | 390 |
| | down | 586 | 116 | 94 |
| durum wheat (Simeto) | up | 1846 | 621 | 477 |
| | down | 1283 | 239 | 178 |
| total | up | 3048 | 1107 | 867 |
| | down | 1869 | 355 | 272 |

**Table S4** Most significant GO categories in emmer (Molise Sel. Colli) and durum wheat (Simeto)

| | categories | GO Terms | p_value emmer | Number of DEGs per GO emmer | p_value durum wheat | Number of DEGs per GO durum wheat |
|---|---|---|---|---|---|---|
| Biological Process up-regulated | cellular component organization or biogenesis | plastid organization | | | 1,21E-14 | 67 |
| | | chloroplast organization | | | 3,16E-10 | 50 |
| | cellular process | translation | 1,40E-47 | 181 | 1,12E-82 | 273 |
| | | peptide biosynthetic process | 2,50E-46 | 181 | 1,61E-81 | 274 |
| | | amide biosynthetic process | 1,08E-45 | 185 | 2,08E-82 | 283 |
| | | peptide metabolic process | 1,81E-45 | 182 | 1,80E-79 | 275 |
| | | cellular amide metabolic process | 8,71E-43 | 186 | 2,98E-77 | 284 |
| | | cellular nitrogen compound biosynthetic process | 8,64E-18 | 283 | 1,25E-29 | 417 |
| | | cellular biosynthetic process | 4,41E-16 | 366 | 4,15E-25 | 535 |
| | | cellular nitrogen compound metabolic process | 6,07E-11 | 367 | 3,48E-19 | 540 |
| | | carboxylic acid metabolic process | 1.45E-05 | 148 | 2,21E-07 | 203 |
| | | plastid organization | | | 1,21E-14 | 67 |
| | | chloroplast organization | | | 3,16E-10 | 50 |
| | | ncRNA metabolic process | | | 2,05E-08 | 102 |
| | | cellular amino acid metabolic process | | | 2,34E-08 | 134 |
| | | oxoacid metabolic process | | | 1,24E-06 | 204 |
| | | organic acid metabolic process | | | 1,61E-06 | 204 |
| | | tetrapyrrole biosynthetic process | | | 5,50E-06 | 39 |
| | | isopentenyl diphosphate metabolic process | | | 9,20E-06 | 40 |
| | | isopentenyl diphosphate biosynthetic process | | | 9,20E-06 | 40 |
| | | cellular macromolecule biosynthetic process | | | 1,49E-05 | 349 |
| | | isopentenyl diphosphate biosynthetic process, methylerythritol 4-phosphate pathway | | | 1,55E-05 | 39 |
| | metabolic process | organonitrogen compound biosynthetic process | 2,07E-68 | 286 | 2,82E-100 | 412 |
| | | organonitrogen compound metabolic process | 3,33E-49 | 310 | 2,21E-78 | 459 |
| | | translation | 1,40E-47 | 181 | 1,12E-82 | 273 |
| | | peptide biosynthetic process | 2,50E-46 | 181 | 1,61E-81 | 274 |
| | | amide biosynthetic process | 1,08E-45 | 185 | 2,08E-82 | 283 |

| | | | | | | |
|---|---|---|---|---|---|---|
| | | peptide metabolic process | 1,81E-45 | 182 | 1,80E-79 | 275 |
| | | cellular amide metabolic process | 8,71E-43 | 186 | 2,98E-77 | 284 |
| | | cellular nitrogen compound biosynthetic process | 8,64E-18 | 283 | 1,25E-29 | 417 |
| | | organic substance biosynthetic process | 8,30E-17 | 372 | 1,24E-21 | 532 |
| | | cellular biosynthetic process | 1,41E-16 | 366 | 4,15E-25 | 535 |
| | | biosynthetic process | 2,21E-14 | 381 | 2,72E-20 | 552 |
| | | nitrogen compound metabolic process | 4,72E-13 | 399 | 6,40E-22 | 591 |
| | | cellular nitrogen compound metabolic process | 6,07E-11 | 367 | 3,48E-19 | 540 |
| | | carboxylic acid metabolic process | 1,45E-05 | 148 | 2,21E-07 | 203 |
| | | gene expression | | | 2,00E-11 | 385 |
| | | ncRNA metabolic process | | | 2,05E-08 | 102 |
| | | cellular amino acid metabolic process | | | 2,34E-08 | 134 |
| | | oxoacid metabolic process | | | 1,24E-06 | 204 |
| | | organic acid metabolic process | | | 1,61E-06 | 204 |
| | | tetrapyrrole biosynthetic process | | | 5,50E-06 | 39 |
| | | isopentenyl diphosphate biosynthetic process | | | 9,20E-06 | 40 |
| | | isopentenyl diphosphate metabolic process | | | 9,20E-06 | 40 |
| | | cellular macromolecule biosynthetic process | | | 1,49E-05 | 349 |
| | | isopentenyl diphosphate biosynthetic process, methylerythritol 4-phosphate pathway | | | 1,55E-05 | 39 |
| | | macromolecule biosynthetic process | | | 6,97E-05 | 351 |
| Molecular Function up-regulated | binding | binding | 1,68E-39 | 507 | 0 | 736 |
| | | RNA binding | 2,22E-19 | 92 | 7,35E-44 | 127 |
| | | heterocyclic compound binding | 4,44E-17 | 330 | 5,31E-80 | 469 |
| | | organic cyclic compound binding | 4,44E-17 | 330 | 5,31E-80 | 469 |
| | | nucleic acid binding | 4,19E-07 | 163 | 1,19E-29 | 222 |
| | | ion binding | | | 9,23E-46 | 404 |
| | | small molecule binding | | | 8,90E-21 | 279 |
| | | nucleoside phosphate binding | | | 3,66E-16 | 267 |
| | | nucleotide binding | | | 3,66E-16 | 267 |
| | | anion binding | | | 2,81E-14 | 249 |
| | | purine ribonucleoside triphosphate binding | | | 6,14E-05 | 196 |
| | catalytic activity | catalytic activity | 4,90E-94 | 518 | 0 | 740 |

| | | | | | | |
|---|---|---|---|---|---|---|
| | | hydrolase activity | 5,99E-09 | 182 | 1,27E-26 | 234 |
| | | oxidoreductase activity | | | 1,27E-26 | 159 |
| | | ligase activity | | | 5,35E-16 | 75 |
| | | ligase activity, forming aminoacyl-tRNA and related compounds | | | 6,69E-14 | 41 |
| | | ligase activity, forming carbon-oxygen bonds | | | 6,69E-14 | 41 |
| | | hydrolase activity, acting on glycosyl bonds | | | 1,47E-13 | 64 |
| | | aminoacyl-tRNA ligase activity | | | 3,00E-13 | 41 |
| | | hydrolase activity, hydrolyzing O-glycosyl compounds | | | 8,83E-12 | 60 |
| | | lyase activity | | | 6,98E-10 | 57 |
| | | transferase activity, transferring one-carbon groups | | | 7,72E-05 | 51 |
| | structural molecule activity | structural constituent of ribosome | 1,16E-92 | 141 | 1,12E-160 | 196 |
| | | structural molecule activity | 7,74E-84 | 146 | 6,84E-141 | 198 |
| Molecular Function down-regulated | binding | binding | | | 0 | 556 |
| | | heterocyclic compound binding | | | 3,70E-90 | 390 |
| | | organic cyclic compound binding | | | 3,70E-90 | 390 |
| | | ion binding | | | 1,92E-74 | 363 |
| | | anion binding | | | 9,34E-50 | 261 |
| | | adenyl ribonucleotide binding | | | 5,67E-45 | 226 |
| | | adenyl nucleotide binding | | | 6,05E-45 | 228 |
| | | carbohydrate derivative binding | | | 5,22E-42 | 238 |
| | | ribonucleotide binding | | | 4,16E-41 | 234 |
| | | ATP binding | | | 1,05E-40 | 209 |
| | | ribonucleoside binding | | | 1,41E-39 | 232 |
| | | nucleoside binding | | | 1,45E-39 | 232 |
| | | purine nucleoside binding | | | 2,18E-39 | 231 |
| | | purine ribonucleoside binding | | | 2,18E-39 | 231 |
| | | purine ribonucleotide binding | | | 2,18E-39 | 231 |
| | | purine nucleotide binding | | | 2,47E-39 | 233 |
| | | nucleoside phosphate binding | | | 7,26E-37 | 252 |
| | | nucleotide binding | | | 7,26E-37 | 252 |
| | | small molecule binding | | | 2,61E-36 | 254 |
| | | purine ribonucleoside triphosphate binding | | | 5,09E-35 | 214 |

| | | | | | | |
|---|---|---|---|---|---|---|
| | catalytic activity | catalytic activity | | | 0 | 566 |
| | | transferase activity | | | 1,23E-36 | 228 |
| | | transferase activity, transferring phosphorus-containing groups | | | 2,45E-28 | 160 |
| | | kinase activity | | | 3,96E-25 | 140 |
| | | phosphotransferase activity, alcohol group as acceptor | | | 1,49E-22 | 134 |
| | | protein kinase activity | | | 1,98E-21 | 122 |
| | | hydrolase activity | | | 2,00E-16 | 174 |
| | | protein serine/threonine kinase activity | | | 1,06E-05 | 58 |
| | transporter activity | transporter activity | | | 3,12E-36 | 120 |
| | | transmembrane transporter activity | | | 1,93E-19 | 84 |
| | | substrate-specific transporter activity | | | 2,79E-06 | 60 |
| | | active transmembrane transporter activity | | | 9,09E-05 | 42 |

**Table S5.** ANOVA for each individual metabolite, relating to their content in the leaves of emmer and durum wheat grown under starvation and optimal N conditions.

| | **G** | **N** | **G x N** |
|---|---|---|---|
| valine | ns | ns | *** |
| glutamic acid | *** | *** | *** |
| Alanine | ns | ns | *** |
| β-alanine | ns | ns | *** |
| Serine | ns | ns | ** |
| glycine | ns | ns | ns |
| Threonine | ns | ns | * |
| aspartic acid | ** | * | *** |
| asparagine | ns | ns | ns |
| thryptophan | * | ns | ns |
| GABA | ns | ns | ** |
| citric acid | ** | ** | ** |
| aconitic acid | ** | ns | *** |
| isocitric acid | ns | ns | ns |
| succinic acid | ns | ns | * |
| fumaric acid | ns | ns | ** |
| malic acid | ns | ns | ** |
| shikimic acid | * | ns | ** |
| quinic acid | ns | ns | *** |
| gluconic acid | ns | ns | ns |
| saccharic acid | ** | * | ** |
| oxalic acid | ns | ns | ns |
| phosphate | ns | ns | ns |
| xylose - lyxose - threalose | ns | ns | ns |
| ribose | ns | ns | ** |
| Fructose | ns | ns | ns |
| mannose - galactose - glucose | ns | ns | ns |
| sucrose | ns | ns | ns |
| isomaltose | ns | ns | * |
| maltose - turanose | ns | ns | ** |
| lactulose | ns | ns | ns |
| maltitol | *** | *** | *** |
| glycerol | ns | ns | ns |
| mannitol | ns | ns | ns |
| sorbitol | ns | ns | ns |
| myo-inositol | ns | ns | ** |
| hexadecanoic acid | ns | ns | ns |
| 1-octacosanol | ns | ns | ns |
| campsterol | ns | ns | ns |
| stigmasterol | ns | ns | ns |
| β-sitosterol | ns | ns | ns |

n.s., not significant; *, $P \leq 0.05$, **, $P \leq 0.01$; ***, $P \leq 0.001$

**Table S6.** Number of edges between central DEGs genes and significantly behaved metabolites under the two N conditions. For each metabolite was highlight in bold the higher number of edges.

| From | To | wheat | Total edge between DEGs and significantly behaved metabolites | Total edge between central DEGs and significantly behaved metabolites |
|---|---|---|---|---|
| DEGs | Aconitic acid | emmer<br>durum wheat | 383<br>1867 | 71<br>**344** |
| DEGs | Alanine | emmer<br>durum wheat | 705<br>1430 | 88<br>**320** |
| DEGs | aspartic acid | emmer<br>durum wheat | 899<br>497 | 143<br>119 |
| DEGs | β-alanine | emmer<br>durum wheat | 37<br>687 | 10<br>**179** |
| DEGs | citric acid | emmer<br>durum wheat | 700<br>464 | 130<br>124 |
| DEGs | fumaric acid | emmer<br>durum wheat | 527<br>548 | 89<br>121 |
| DEGs | GABA | emmer<br>durum wheat | 12<br>2954 | 1<br>**479** |
| DEGs | glutamic acid | emmer<br>durum wheat | 1360<br>101 | **201**<br>19 |
| DEGs | isocitric acid | emmer<br>durum wheat | 44<br>53 | **21**<br>7 |
| DEGs | Isomaltose | emmer<br>durum wheat | 338<br>28 | **55**<br>0 |
| DEGs | malic acid | emmer<br>durum wheat | 213<br>851 | 54<br>**199** |
| DEGs | maltose-turanose | emmer<br>durum wheat | 254<br>1796 | 36<br>**299** |
| DEGs | myo-inositol | emmer<br>durum wheat | 36<br>2725 | 5<br>**460** |
| DEGs | maltitol | emmer<br>durum wheat | 1667<br>1977 | 431<br>391 |
| DEGs | quinic acid | emmer<br>durum wheat | 244<br>2873 | 60<br>**474** |
| DEGs | Ribose | emmer<br>durum wheat | 131<br>1906 | 36<br>**355** |
| DEGs | saccharic acid | emmer<br>durum wheat | 52<br>59 | **23**<br>2 |
| DEGs | Serine | emmer<br>durum wheat | 486<br>36 | **77**<br>1 |
| DEGs | shikimic acid | emmer<br>durum wheat | 244<br>1564 | 52<br>**270** |
| DEGs | succinic acid | emmer<br>durum wheat | 218<br>52 | **47**<br>0 |
| DEGs | Threonine | emmer<br>durum wheat | 266<br>50 | **50**<br>0 |
| DEGs | Valine | emmer<br>durum wheat | 1390<br>2484 | 218<br>**427** |

**Table S7.** Putative functions of common DEGs in both tetraploid wheats with central roles in either of emmer-specific and durum wheat-specific networks.

| DEGs | up/down | metabolites | group | Putative function description | Orthologues | Central role emmer | Central role durum wheat |
|---|---|---|---|---|---|---|---|
| Traes_1BL_AAC4D956E | *down* | Aconitic acid, aspartic acid, citric acid, fumaric acid, glutamic acid, maltitol, valine | C metabolism (glycolysis) | Pyruvate, phosphate dikinase 1(PPDK) (EC 2.7.9.1) | AT4G15530 | **x** | |
| Traes_2AL_F1C4FF20C | *down* | Citric acid, glutamic acid, maltitol, valine | Fatty acid metabolism | Lecithin-cholesterol acyltransferase-like 4 (EC 2.3.1-) | AT4G19860 | **x** | |
| Traes_1AL_053F6B12B | *up* | Alanine, aspartic acid, b-alanine, fumaric acid, glutamic acid, isocitric acid, isomaltose, malic acid, maltose-turanose, maltitol, quinic acid, ribose, saccharic acid, serine, shikimic acid, succinic acid, threonine, valine | C metabolism (photosynthesis) | Chlorophyll synthase (CHLG) (EC 2.5.1.62) | AT3G51820 | **x** | |
| Traes_1AL_FE1097B22 | *up* | Maltitol | Others | Tyrosine--tRNA ligase, chloroplastic/mitochondrial (EC 6.1.1.1) | AT3G02660 | **x** | |
| Traes_2AS_F485758C1 | *up* | Citric acid, glutamic acid, maltitol, valine | C metabolism (glycolysis) | Pyruvatedehydrogenase E1 component subunit alpha-3 (PDH-E1 ALPHA) (EC 1.2.4.1) | AT1G01090 | **x** | |
| Traes_2BS_959A4E58A | *up* | Citric acid, glutamic acid, maltitol, valine | Kinases | Fructokinase-like 2 (FLN 2) | AT1G69200 | **x** | |
| Traes_4AL_45E27280D | *up* | Alanine, serine, valine, maltitol | Amino acid metabolism | Cysteine desulfurase 1 (NIFS) (EC 2.8.1.7) | AT1G08490 | **x** | |
| Traes_3AS_D097872BE | *up* | Aconitic acid, alanine, aspartic acid, fumaric acid, glutamic acid, malic acid, maltitol, valine | Transcription factors | DEAD-box ATP-dependent RNA helicase 39 (EC 3.6.4.13) | AT4G09730 | **x** | |
| Traes_6AS_FD8F6B539 | *up* | Maltitol, valine | Stress | Rhodanese-like domain-containing protein 7 (STR 7) | AT2G40760 | **x** | |
| Traes_7AL_A1ECABC74 | *up* | Alanine, fumaric acid, glutamic acid, isocitric acid, isomaltose, maltitol, saccharic acid, serine, threonine | Others | 30S ribosomal protein S31 | AT2G38140 | **x** | |
| Traes_7BS_3285CDC24 | *up* | Fumaric acid, threonine | Stress | F-box protein MAX2 | AT2G42620 | **x** | |

| | | | | | | | |
|---|---|---|---|---|---|---|---|
| Traes_2BL_CCD296233 | *down* | Emmer (Citric acid, glutamic acid, maltitol, valine)<br><br>Durum wheat (Aconitic acid, alanine, GABA, malic acid, maltose-turanose, myo-inositol, maltitol, quinic acid, ribose, valine) | Stress | Stress enhanced protein 2 (SEP2) | AT2G21970 | **x** | **x** |
| Traes_2AL_DB64E18A1 | *down* | GABA, myo-inositol, maltitol, quinic acid, ribose, valine | Purine metabolism | Allantoinase (ALN) (EC 3.5.2.5) | AT4G04955 | | **x** |
| Traes_7BS_7E9C070B8 | *down* | Aconitic acid, alanine, b-alanine, citric acid, GABA, malic acid, maltose-turanose, myo-inositol, maltitol, quinic acid, shikimic acid, valine | C metabolism (photorespiration) | Carboxyl-terminal-processing peptidase 3 (CTPA3)<br><br>(EC 3.4.21.102) | AT3G57680 | | **x** |
| Traes_1AL_928DC3A3C | *up* | Aconitic acid, alanine, aspartic acid, citric acid, GABA, maltose-turanose, myo-inositol, maltitol, quinic acid, ribose, shikimic acid, valine | Riboflavine metabolism | Bifunctional riboflavin kinase/FMN phosphatase (FMN/FHY)<br>(EC 3.1.3.102/ EC 2.7.1.26) | AT4G21470 | | **x** |
| Traes_3AL_8C6F4F66A | *up* | Aconitic acid, citric acid, GABA, maltose-turanose, myo-inositol, quinic acid, ribose, shikimic acid, valine | Terpenoid backbone biosynthesis | 4-diphosphocytidyl-2-C-methyl-D-erythritol kinase (ISPE)<br>(EC 2.7.1.148) | AT2G26930 | | **x** |
| Traes_3AS_922982272 | *up* | Alanine, b-alanine, fumaric acid | C metabolism (glycolisis) | Probable zinc metalloprotease EGY2 (EC 3.4.24-) | AT5G05740 | | **x** |
| Traes_4AS_0B56CEBD5 | *up* | Aconitic acid, GABA, maltose-turanose, myo-inositol, quinic acid, ribose, shikimic acid, valine | stress | ATP-dependent Clp protease proteolytic subunit-related protein 3 (CLPR3) | AT1G09130 | | **x** |
| Traes_4BS_0ECAB85C2 | *up* | Aconitic acid, GABA, isocitric acid, maltose-turanose, myo-inositol, quinic acid, ribose, valine | others | 30S ribosomal protein S13 | AT5G14320 | | **x** |
| Traes_4AL_199EE1839<br><br>Traes_5BL_BD895CF84 | *up* | Aconitic acid, alanine, citric acid, GABA, malic acid, maltose-turanose, myo-inositol, maltitol, quinic acid, | Transcription factors | DEAD-box ATP-dependent RNA helicase 3 (EC 3.6.4.13) | AT5G26742 | | x |

| | | | | | | | |
|---|---|---|---|---|---|---|---|
| | | ribose, shikimic acid, valine<br><br>Aconitic acid, alanine, b-alanine, citric acid, GABA, malic acid, maltose-turanose, myo-inositol, maltitol, quinic acid, ribose, shikimic acid, valine | | | | | |
| Traes_5BL_0684B3A02 | *up* | Aconitic acid, alanine, citric acid, GABA, maltose-turanose, myo-inositol, maltitol, quinic acid, ribose, shikimic acid, valine | Energy pathways | Cytochrome c biogenesis protein (CCS1) | AT1G49380 | | **x** |
| Traes_5BL_CFC7BB483 | *up* | Aconitic acid, alanine, citric acid, GABA, maltose-turanose, myo-inositol, maltitol, quinic acid, ribose, shikimic acid, valine | Porphyrin and chlorophyll metabolism | Magnesium-chelatase subunit ChlD (EC 6.6.1.1) | AT1G08520 | | **x** |
| Traes_6BL_34B508140 | *up* | Aconitic acid, alanine, GABA, malic acid, maltose-turanose, myo-inositol, maltitol, quinic acid, valine | others | Pentatricopeptide repeat-containing protein At4g39620 | AT4G39620 | | **x** |

**Table S8.** Putative function of DEGs specific to each of the tetraploid wheats with central roles in either of emmer-specific and durum wheat-specific networks

| DEGs | up/down | metabolites | group | Putative function description | Orthologues | Central role emmer | Central role durum wheat |
|---|---|---|---|---|---|---|---|
| Traes_1AL_5F0F4FAFB | *down* | aspartic acid, citric acid glutamic acid, maltitol valine | kinases | Non-specific serine/threonine protein kinase (EC 2.7.11.1) | | **x** | |
| Traes_3AL_883A4805C | *down* | aconitic acid, aspartic acid, citric acid, fumaric acid, glutamic acid, maltitol, valine | other C metabolisms | Cytokinin riboside 5'-monophosphate phosphoribohydrolase (EC 3.2.2.n1) | AT2G37210 | **x** | |
| Traes_4BL_4D370C9FA<br>Traes_4BL_4D370C9FA1 | *down* | Alanine, fumaric acid glutamic acid, maltitol, threonine<br>Alanine, fumaric acid glutamic acid, isomaltose, maltitol serine, valine | stress | Probable mediator of RNA polymerase II transcription subunit 37c | AT3G12580 | x | |
| Traes_4BL_96C906D61 | *down* | citric acid, glutamic acid maltitol, valine | stress | Non-specific lipid-transfer protein | AT3G08770 | **x** | |
| Traes_5AL_6FB526190 | *down* | citric acid, glutamic acid maltitol, valine | N metabolism | Carbonic anhydrase (EC 4.2.1.1) | AT5G14740 | **x** | |
| Traes_5BL_A0ED5623C | *down* | Maltitol, valine | stress | Phospholipase D delta (PLDDELTA**)** (EC3.1.4.4) | AT4G35790 | **x** | |
| Traes_7AS_9A01E6B17 | *down* | aconitic acid, aspartic acid, citric acid, glutamic acid, maltitol valine | transporter | Potassium transporter | AT2G30070 | **x** | |
| Traes_7BL_8930CD1E5 | *down* | aspartic acid, citric acid glutamic acid, palatinose -maltitol, valine | Stress | Peptide methionine sulfoxide reductase B1, chloroplastic (EC 1.8.4.12) | AT1G53670 | **x** | |
| Traes_1AL_4B7636BC9 | *up* | Valine, maltitol | Stress | Protein LOL1 | AT1G32540 | **x** | |
| Traes_1AL_090571678<br>Traes_1BL_5F74C58ED | *up* | Alanine, aspartic acid fumaric acid, glutamic acid, isomaltose, malic acid, maltose-turanose, maltitol quinic acid, ribose, serine, shikimic acid, valine<br>alanine, glutamic acid, maltose-turanose, | transporter | Secretory carrier-associated membrane protein (SCAMP) | AT1G61250 | **x** | |

| | | | | | | |
|---|---|---|---|---|---|---|
| | | maltitol | | | | |
| Traes_1AL_1538AC680 | *up* | alanine, citric acid, glutamic acid, maltose-turanose, maltitol, valine | Kinases | Nucleoside diphosphate kinase (EC 2.7.4.6) | AT4G23895 | **x** |
| Traes_1BL_ACC9DDA86 | | citric acid, glutamic acid maltitol, valine | | | | |
| Traes_1AL_8E8E7401B | *up* | alanine, maltose-turanose, maltitol, serine, valine | other C metabolisms | Beta-hexosaminidase (EC 3.2.1.52) | AT3G49850 | **x** |
| Traes_1BL_7F9006C51 | *up* | glutamic acid, maltitol, valine | Stress | Hyperosmolality-gated Ca2+ permeable channel | | **x** |
| Traes_1BL_C706FAA2B | *up* | glutamic acid maltitol, valine | Others | Ribosomal protein L17 | AT3G54210 | **x** |
| Traes_2AL_5BAB26827 | *up* | citric acid, maltitol, valine | Stress | Lipoxygenase (EC 1.13.11-) | AT3G45140 | **x** |
| Traes_2AS_34272605C | *up* | alanine, aspartic acid, fumaric acid, glutamic acid, isocitric acid isomaltose, maltitol, quinic acid, ribose, saccharic acid, serine shikimic acid, succinic acid, threonine, valine | Stress | Tubulin alpha chain | AT4G14960 | **x** |
| Traes_2AS_7D3EC9FC2 | *up* | citric acid, glutamic acid maltitol, valine | Others | E3 ubiquitin-protein ligase | | **x** |
| Traes_3AS_CD8EF3E96 | *up* | citric acid, glutamic acid maltitol, valine | Others | FRIGIDA-like protein | AT5G16320 | **x** |
| Traes_4AL_17C4B47B3 | *up* | aconitic acid, aspartic acid, glutamic acid, maltitol | Others | Pentatricopeptide repeat-containing protein At4g31850, chloroplastic | AT4G31850 | **x** |
| Traes_4AS_4C25DF6F4 | *up* | aspartic acid, citric acid glutamic acid, maltitol valine | others C metabolism | Beta-galactosidase (EC 3.2.1.23) | AT2G32810 | **x** |
| Traes_5AL_CEEDF43FC | *up* | alanine, aspartic acid, isomaltose, maltose-turanose, maltitol, ribose, serine, valine | Fatty acid metabolism | Acyl-[acyl-carrier-protein] hydrolase (EC 3.1.2-) | AXX17-AT1G08340 | **x** |
| Traes_6AL_006779D22 | *up* | aconitic acid, alanine, aspartic acid, citric acid, fumaric acid, glutamic acid, malic acid, maltitol, quinic acid, serine, valine | other C metabolisms | Xyloglucan endotransglucosylase/hydrolase (EC 2.4.1.207) | AT2G06850 | **x** |

| | | | | | | | |
|---|---|---|---|---|---|---|---|
| Traes_6BS_C9B216D97 | *up* | alanine, aspartic acid, citric acid, glutamic acid malic acid, maltitol quinic acid, valine | C metabolism (glycolysis) | Acetyltransferase component of pyruvate dehydrogenase complex (EC 2.3.1.12) | AXX17-AT1G48680 | **x** | |
| Traes_7AS_378A12AA9 | *up* | aconitic acid, aspartic acid, citric acid, fumaric acid, glutamic acid, maltitol, quinic acid, shikimic acid, valine | Others | TGW-7A | | **x** | |
| Traes_7AS_A80BE362A | *up* | aconitic acid, aspartic acid, citric acid glutamic acid, maltitol valine | others C metabolism | Sucrose synthase (EC 2.4.1.13) | AT5G49190 | **x** | |
| Traes_7BL_E2E7AA383 | *up* | alanine, aspartic acid, fumaric acid, isocitric acid, isomaltose, myo-inositol, maltitol, quinic acid, ribose, serine, shikimic acid, succinic acid, threonine | Stress | Protein Overexpressor of Cationic Peroxidase 3 | AT5G11270 | **x** | |
| Traes_1AL_A970EBB27 | *down* | aconitic acid, alanine, b-alanine, fumaric acid, GABA, malic acid, maltose-turanose, myo-inositol, maltitol, quinic acid, ribose, valine | Stress | Protein Detoxification 42 | AT1G51340 | | **x** |
| Traes_1AS_AEEC8D740 | *down* | alanine, b-alanine, GABA, maltose-turanose, myo-inositol, maltitol, quinic acid, valine | Others | E3 ubiquitin-protein ligase UPL6 (EC 2.3.2.26) | AT3G17205 | | **x** |
| Traes_1BL_9772C6F45 | *down* | alanine, aspartic acid beta-alanine, fumaric acid, GABA, malic acid myo-inositol, maltitol quinic acid, ribose, shikimic acid, valine | transporter | Calcium-transporting ATPase (EC 3.6.3.8) | AT1G27770 | | **x** |
| Traes_1BL_A9AC1340E | *down* | aconitic acid, alanine, GABA, malic acid, maltose-turanose, myo-inositol, maltitol quinic acid, ribose, valine | Transcription factors | Auxin response factor (ARF family) | AT5G20730 | | **x** |
| Traes_1BS_4A63D1DBC | *down* | GABA, quinic acid, ribose, shikimic acid, valine | other C metabolisms | Phosphatidate cytidylyltransferase (EC 2.7.7.41) | AT4G22340 | | **x** |
| Traes_2AL_BDC8DD6C7 | *down* | aconitic acid, alanine aspartic acid, beta-alanine, fumaric acid, | kinases | Serine/threonine-protein kinase (EC 2.7.11.1) | | | **x** |

| | | | | | | | |
|---|---|---|---|---|---|---|---|
| | | GABA, malic acid, maltose-turanose myo-inositol, maltitol, quinic acid, ribose, valine | | | | | |
| Traes_2AL_E80950527 | *down* | alanine, GABA, maltose-turanose, myo-inositol, maltitol, quinic acid, valine | transporter | Chloride channel protein | AT5G40890 | | **x** |
| Traes_2AL_EC2CE85AC | *down* | GABA, myo-inositol, quinic acid, ribose | others | RING-type E3 ubiquitin transferase (EC 2.3.2.27) | ATG23140 | | **x** |
| Traes_2AL_F4F6F518D | *down* | alanine, GABA, myo-inositol, quinic acid, ribose, maltitol | stress | Amine oxidase (EC 1.4.3-) | AT4G12290 | | **x** |
| Traes_2BL_19B3E60AA | *down* | aconitic acid, alanine, citric acid, GABA, malic acid, maltose-turanose, myo-inositol, maltitol, quinic acid, ribose, shikimic acid, valine | transporter | Probable copper-transporting ATPase HMA5 (EC 3.6.3.54) | AT1G63440 | | **x** |
| Traes_2BL_BB2C0F695 | *down* | aconitic acid, alanine, beta-alanine, GABA myo-inositol, maltitol, quinic acid, ribose | others | Histone H3 | AT5G65350 | | **x** |
| Traes_3AL_FFE4C92D8 | *down* | aconitic acid, alanine, citric acid, GABA, malic acid, maltose-turanose, myo-inositol, maltitol, quinic acid, shikimic acid, valine | others C metabolism | Cellulose synthase A catalytic subunit 8 [UDP-forming] (EC 2.4.1.12) | AT4G18780 | | **x** |
| Traes_4AL_D7702A458 | *down* | aconitic acid, beta-alanine, fumaric acid, GABA, myo-inositol, maltitol, quinic acid, ribose, shikimic acid, valine | transporter | Plasma membrane ATPase (EC 3.6.3.6) | AT2G18960 | | **x** |
| Traes_5AL_07C125B1C | *down* | alanine, beta-alanine, GABA, quinic acid, maltose-turanose, valine myo-inositol, shikimic acid, maltitol | stress | Hypersensitive induced reaction protein 2 | AT1G69840 | | **x** |
| Traes_5AL_13E2DEC48 | *down* | aconitic acid, GABA myo-inositol, valine, maltitol, quinic acid, ribose | Transcription factors | MIKC-type MADS-box transcription factor WM6 | | | **x** |

| | | | | | | | |
|---|---|---|---|---|---|---|---|
| Traes_5AL_23B24917C | *down* | aconitic acid, alanine, beta-alanine, citric acid GABA, malic acid, maltose-turanose, valine myo-inositol, shikimic acid, maltitol, quinic acid, ribose | C metabolism (photosynthesis) | Ferrochelatase (EC 4.99.1.1) | AT2G30390 | | **x** |
| Traes_5BL_FF9AA4B58 | | aconitic acid, alanine, citric acid, GABA, maltose-turanose, myo-inositol, maltitol, quinic acid, ribose, shikimic acid, valine | | | | | |
| Traes_5AS_BA57B5B56 | *down* | aconitic acid, GABA, myo-inositol, valine maltitol, quinic acid, ribose | Purine/pyrimidine metabolism | Ureidoglycolate hydrolase (allantoate amido hydrolase) (EC 3.5.1.116) | AT5G43600 | | **x** |
| Traes_5BL_369BF7271 | *down* | GABA, myo-inositol, maltitol, quinic acid | others | Thioredoxin-like protein HCF164 | AT4G37200 | | **x** |
| Traes_5BL_E123D8905 | *down* | aconitic acid, alanine, citric acid, GABA, malic acid, maltose-turanose, myo-inositol, maltitol, quinic acid, ribose, valine | others | ALBINO3-like protein 1, chloroplastic | AT1G24490 | | **x** |
| Traes_6AS_0C1D497EA | *down* | aconitic acid, alanine, b-alanine, citric acid, GABA, malic acid, valine, maltose-turanose, myo-inositol, maltitol, quinic acid, ribose, shikimic acid | Amino acid metabolism | S-adenosylmethionine synthase 4 (EC 2.5.1.6) | AT3G17390 | | **x** |
| Traes_6AS_F3CC61A36 | *down* | GABA, quinic acid, shikimic acid, valine | other C metabolisms | Terpene cyclase/mutase family member (EC5.4.99-) | AT1G78950 | | **x** |
| Traes_6BL_2B67C5E30 | *down* | aconitic acid, GABA, maltose-turanose, myo-inositol, quinic acid, valine | transporter | Glycolipid transfer protein 1 (GLTP 1) | AT2G33470 | | **x** |
| Traes_6BS_A80520954 | *down* | aconitic acid, alanine, aspartic acid, b-alanine, citric acid, fumaric acid, GABA, malic acid, maltose-turanose, myo-inositol, maltitol, quinic | other C metabolisms | Phosphopantetheine-adenylyltransferase (PPAT) (EC 2.7.7.3) | AT2G18250 | | **x** |

| | | | | | | | |
|---|---|---|---|---|---|---|---|
| | | acid, ribose, shikimic acid, valine | | | | | |
| Traes_7AS_E477E850F | *down* | aconitic acid, alanine, aspartic acid, beta-alanine, fumaric acid, GABA, malic acid, maltose-turanose, myo-inositol, maltitol, quinic acid, valine | Stress | Protein DETOXIFICATION | AT4G25640 | | **x** |
| Traes_1AL_4BF6C25E1 | *up* | aconitic acid, citric acid, GABA, maltose-turanose, myo-inositol, quinic acid, ribose, shikimic acid, valine | Others | Probable transmembrane GTPase FZO-like | AT1G03160 | | **x** |
| Traes_1AL_1A98A80B4 | *up* | aconitic acid, GABA myo-inositol, quinic acid, ribose, shikimic acid, valine | C metabolism (photosynthesis) | Chlorophyll synthase, chloroplastic (EC 2.5.1.62) | AT3G51820 | | **x** |
| Traes_1BL_0FC423766 | *up* | GABA, myo-inositol, maltitol, quinic acid, valine | Others | 60S ribosomal protein L36 | AT5G02440 | | **x** |
| Traes_1BL_714F4E4AC | *up* | aconitic acid, GABA, maltose-turanose, myo-inositol, quinic acid, ribose, shikimic acid, valine | Stress | Carotene epsilon-monooxygenase, chloroplastic (EC 1.14.99.45) | AT3G53130 | | **x** |
| Traes_1BL_8AC8D124C | *up* | aconitic acid, citric acid, GABA, myo-inositol, quinic acid, ribose, shikimic acid, valine | Others | 50S ribosomal protein L1 | AT3G63490 | | **x** |
| Traes_2AL_E7D1C8AA5 | *up* | GABA, maltose-turanose, myo-inositol, maltitol, quinic acid, shikimic acid, valine | transporter | Probable inositol transporter 2 | AT1G30220 | | **x** |
| Traes_2AL_FA3E7C60B | *up* | aconitic acid, GABA, isocitric acid, maltose-turanose, myo-inositol, maltitol, quinic acid, valine | Stress | Violaxanthin de-epoxidase, chloroplastic (VDE 1) (EC 1.23.5.1) | AT1G08550 | | **x** |
| Traes_2AL_20BC426B9 | *up* | aconitic acid, aspartic acid, citric acid, GABA myo-inositol, maltitol quinic acid, ribose, valine | Stress | Peroxidase (EC 1.11.1.7) | AT5G17820 | | **x** |

| | | | | | | | |
|---|---|---|---|---|---|---|---|
| Traes_2AL_729CB1040 | *up* | GABA, myo-inositol quinic acid, ribose, valine | C metabolism (glycolysis) | Aldose 1-epimerase (EC 5.1.3.3) | AT3G17940 | | **x** |
| Traes_2AL_E06A248EE | *up* | alanine, beta-alanine, fumaric acid, GABA myo-inositol, maltitol, quinic acid, ribose, shikimic acid, valine | Stress | Glutathione peroxidase (EC 1.11.1.9) | AT2G48150 | | **x** |
| Traes_2BL_363CA4ED8 | *up* | aconitic acid, alanine, GABA, maltose-turanose, myo-inositol, maltitol, quinic acid, ribose, shikimic acid, valine | C metabolism (photosynthesis) | Ferredoxin-3 | AT2G27510 | | **x** |
| Traes_2BS_FF8887BE5 | *up* | aconitic acid, citric acid, GABA, maltose-turanose, myo-inositol, quinic acid, ribose, shikimic acid, valine | Others | Asparagine--tRNAligase, chloroplastic/mitochondrial | AT4G17300 | | **x** |
| Traes_2BS_7135D68C5 | *up* | alanine, GABA, myo-inositol, maltitol, quinic acid, ribose | Others | Ribosomal protein L19 | AXX17-AT4G02830 | | **x** |
| Traes_2BS_E5A5144E0 | *up* | alanine, fumaric acid GABA, maltitol, quinic acid, ribose | Others | tRNA (guanine(26)-N(2))-dimethyltransferase | | | **x** |
| Traes_3AS_3CB8A9C01 | *up* | aconitic acid, GABA, myo-inositol, quinic acid ribose, valine | Amino acid metabolism | Glutamate decarboxylase (EC 4.1.1.15) | AT1G65960 | | **x** |
| Traes_3B_00CF2A894 | *up* | GABA, myo-inositol, quinic acid, ribose, shikimic acid, valine | Stress | ATP-dependent Clp protease proteolytic subunit 3 (EC 3.4.21.92) | AT1G66670 | | **x** |
| Traes_4AL_497775E8F | *up* | alanine, beta-alanine GABA, myo-inositol, Maltitol, quinic acid, ribose, shikimic acid, valine | Stress | Ferritin (EC 1.16.3.1) | AT3G56090 | | **x** |
| Traes_4BS_8D508569E | *up* | aconitic acid, GABA, myo-inositol, quinic acid, ribose, shikimic acid, valine | Others | Pentatricopeptide repeat-containing protein At3g06430 | AT3G06430 | | **x** |
| Traes_5AL_955D2899A | *up* | aconitic acid, alanine GABA, maltose-turanose, myo-inositol maltitol, quinic acid, ribose, shikimic acid, valine | Stress | Pectin acetylesterase (EC 3.1.1-) | AT5G23870 | | **x** |

| | | | | | | | |
|---|---|---|---|---|---|---|---|
| Traes_5AS_6D0CC3B1F | *up* | GABA, maltose-turanose, myo-inositol, Maltitol, quinic acid, shikimic acid, valine | Others | 60S ribosomal protein L36 | AT5G02440 | | **x** |
| Traes_5BL_304FAFA26 | *up* | GABA, myo-inositol, maltitol, quinic acid, ribose, valine | Stress | Lipoxygenase (EC 1.13.11-) | AT3G45140 | | **x** |
| Traes_6AL_D13D9D951 | *up* | aconitic acid, GABA, maltose-turanose, myo-inositol, maltitol, quinic acid, ribose, valine | stress | Serine carboxypeptidase-like 34(EC 3.4.16-) | AT5G23210 | | **x** |
| Traes_6AL_58B6B7319 | *up* | aconitic acid, alanine, aspartic acid, citric acid, GABA, malic acid, maltose-turanose, myo-inositol, maltitol, quinic acid, ribose, valine | transporter | Bidirectional sugar transporter SWEET | AT3G16690 | | **x** |
| Traes_6AS_0F0809F96 | *up* | aconitic acid, GABA, maltose-turanose, myo-inositol, maltitol, quinic acid, ribose, shikimic acid, valine | stress | Probable inactive linolenate hydroperoxide lyase | AT4G15440 | | **x** |
| Traes_6AS_87906149C | *up* | aconitic acid, GABA, maltose-turanose, myo-inositol, quinic acid, ribose, shikimic acid, valine | others | Protein Chlororespiratory Reduction 6 (CRR6) | AT2G47910 | | **x** |
| Traes_7AL_356F6FF0E | *up* | aconitic acid, citric acid, GABA, maltose-turanose, myo-inositol, maltitol, valine | others | Blue-light photoreceptor PHR2 | AT2G47590 | | **x** |
| Traes_4AL_320703FD2 | *up* | aconitic acid, alanine, aspartic acid, fumaric acid, GABA, malic acid, myo-inositol, maltitol, quinic acid, ribose, valine | other C metabolisms | Xyloglucan endotransglucosylase/hydrolase (EC 2.4.1.207) | AT2G06850 | | **x** |
| Traes_7AL_1B1FBCDE4 | | aconitic acid, alanine, beta-alanine, citric acid, GABA, malic acid maltose-turanose, myo-inositol, | | | | | |

| | | | | | | | |
|---|---|---|---|---|---|---|---|
| | | maltitol, quinic acid, ribose, valine | | | | | |
| Traes_7AL_8C4A9BEBF | | aconitic acid, alanine, aspartic acid, b-alanine, citric acid, fumaric acid, GABA, malic acid, maltose-turanose, myo-inositol, maltitol, quinic acid, ribose, shikimic acid, valine | | | | | |
| Traes_7AS_020CCE3DB | *up* | aconitic acid, citric acid, GABA, maltose-turanose, myo-inositol, maltitol, quinic acid, ribose, valine | Stress | Peroxiredoxin Q, chloroplastic (EC 1.11.1.15) | AT3G26060 | | **x** |
| Traes_7AS_545BDB6F1 | *up* | aconitic acid, GABA, maltose-turanose, myo-inositol, quinic acid, ribose, shikimic acid, valine | Others | Alanine--tRNA ligase, chloroplastic/mitochondrial (EC 6.1.1.7) | AT5G22800 | | **x** |
| Traes_7BL_A42D6C984 | *up* | aconitic acid, GABA, maltose-turanose, myo-inositol, maltitol, quinic acid, ribose, valine | Stress | Superoxidedismutase [Cu-Zn] 2 (EC 1.15.1.1) SOD2 | AT2G28190 | | **x** |
| Traes_7BL_B5491A48C | *up* | aconitic acid, alanine, citric acid, GABA, malic acid, maltose-turanose, myo-inositol, maltitol, quinic acid, ribose, shikimic acid, valine | Amino acid metabolism | D-3-phosphoglycerate dehydrogenase (EC 1.1.1.95) | AXX17-AT4G39120 | | **x** |
| Traes_7BL_C35CD97E7 | *up* | aconitic acid, alanine, citric acid, GABA, maltose-turanose, myo-inositol, maltitol, quinic acid, ribose, shikimic acid, valine | C metabolism (photosynthesis) | Delta-aminolevulinic acid dehydratase (EC 4.2.1.24) | AXX17-AT1G63860 | | **x** |
